# Insights into bear evolution from a Pleistocene polar bear genome

**DOI:** 10.1101/2021.12.11.472228

**Authors:** Tianying Lan, Kalle Leppälä, Crystal Tomlin, Sandra L. Talbot, George K. Sage, Sean D. Farley, Richard T. Shideler, Lutz Bachmann, Øystein Wiig, Victor A. Albert, Jarkko Salojärvi, Thomas Mailund, Daniela I. Drautz-Moses, Stephan C. Schuster, Luis Herrera-Estrella, Charlotte Lindqvist

## Abstract

The polar bear (*Ursus maritimus*) has become a symbol of the threat to biodiversity from climate change. Understanding polar bear evolutionary history may provide insights into apex carnivore responses and prospects during periods of extreme environmental perturbations. In recent years, genomic studies have examined bear speciation and population history, including evidence for ancient admixture between polar bears and brown bears (*Ursus arctos*). Here, we extend our earlier studies of a 130,000–115,000-year-old polar bear from the Svalbard Archipelago using a 10X coverage genome sequence and ten new genomes of polar and brown bears from contemporary zones of overlap in northern Alaska. We demonstrate a dramatic decline in effective population size for this ancient polar bear’s lineage, followed by a modest increase just before its demise. A slightly higher genetic diversity in the ancient polar bear suggests a severe genetic erosion over a prolonged bottleneck in modern polar bears. Statistical fitting of data to alternative admixture graph scenarios favors at least one ancient introgression event from brown bears into the ancestor of polar bears, possibly dating back over 150,000 years. Gene flow was likely bidirectional, but allelic transfer from brown into polar bear is the strongest detected signal, which contrasts with other published works. These findings may have implications for our understanding of climate change impacts: polar bears, a specialist Arctic lineage, may not only have undergone severe genetic bottlenecks, but also been the recipient of generalist, boreal genetic variants from brown bear during critical phases of Northern Hemisphere glacial oscillations.

**Significance:** Interspecific hybridization is a widespread phenomenon, but measuring its extent, directionality, and adaptive importance remains challenging. Ancient genomes, however, can help illuminate the history of modern organisms. Here, we present a genome retrieved from a 130,000–115,000-year-old polar bear and perform genome analyses of modern polar and brown bears throughout their geographic range. We find that the principal direction of ancient allele sharing was from brown bear into polar bear, although gene flow between them has likely been bidirectional. This inverts the current paradigm of unidirectional gene flow from polar into brown bear, and it suggests that polar bears were recipients of external genetic variation prior to their extensive population decline.

## Introduction

The polar bear (*Ursus maritimus*) has become a symbolic species for ascertaining the impact of climate change on biodiversity and species evolution. With their dependence on sea ice, polar bears owe their continuing survival to the future stability of the vast Arctic regions of the planet. In connection, given Pleistocene oscillations between glacial and interglacial periods, polar bear paleohistory must hold clues to future responses to changing Earth climates. High coverage genomes from ancient polar bear remains can therefore provide invaluable insights regarding prior adaptative resilience of the species to extreme environmental fluctuations in the past. Moreover, should such ancient polar bears be appropriately placed in age, their paleogenomes could illuminate the lineage split from the species’ lower-latitude sister taxon, the brown bear (*Ursus arctos*), in addition to enlightening any post-divergence admixture between the two species. However, polar bear fossils are very rare, with most dating to the Holocene period (1–3). In 2012 (4), extensive genomic data were generated from 23 extant polar bears, and a draft genome was presented from a stratigraphically validated 115,000–130,000-year-old polar bear jawbone of Eemian interglacial age that was recovered from the Svalbard archipelago of Norway (1, 5). At the time, that study successfully pushed the age record of a sequenced vertebrate genome toward the Middle Pleistocene, but the initial draft genome was of low coverage (<1X depth), limiting its utility in genome-scale analyses.

Several additional genomic studies have since sought to trace polar bear evolution, a species that has emerged for uncovering complex speciation processes associated with interspecific admixture and rapid evolutionary adaptation (6–10). Although the polar bear and brown bear are recognized as closely related yet highly distinct species, studies so far strongly point to ancient and even ongoing (11) introgressive hybridization between the two lineages. This work has mostly centered on polar bear admixture with brown bears in Alaska’s Alexander Archipelago, because mitochondrial haplotypes of brown bears in the archipelago today (the so-called ABC brown bears) are more similar to polar bear haplotypes than they are to haplotypes found in most non-ABC brown bears (3, 6, 12–14). The deep nesting of polar bears within the brown bear maternal lineage, along with the fact that several other, both modern and extinct, brown bear populations share mitochondrial haplotypes with polar bears (15–17), implies a much more complex evolutionary history beyond only the Alexander Archipelago. Indeed, analyses of bear nuclear genomes have suggested widespread allele sharing among polar bears and brown bears, including extinct Irish brown bears (7), albeit with the highest proportion of allele sharing found between polar bears and ABC brown bears (8, 9). The nature of this allele sharing has been interpreted to represent multiple polar bear introgressions into various brown bear lineages (7), but this directionality, although broadly accepted, is not conclusively established.

Population genomic analyses have also identified an ancient and drastic decline in polar bear effective population size over the past 300,000 years, reflecting the far lower genetic diversity among extant polar bears compared to brown bears (4, 9). The complex population histories of the two sister species have challenged models for estimating divergence times and left a conundrum concerning the age of the polar bear as a species. Applying an extended coalescence hidden Markov model based on isolation-with-migration, an initial split time between brown and polar bears and American black bear was estimated to be approximately 5–4 million years ago (ma), followed by a period of gene flow before a complete split ~200 thousand years ago (ka) (4). However, other estimates have generally agreed on a much younger split, although spanning a large interval from ~1.6 ma to 200 ka (9, 18).

The complex model for polar bear evolution that suggests multiple introgression events from polar bear into brown bear (7) warrants further scrutiny with a more complete sampling of crucial North American brown bear populations and methodologies that permit explicit testing of alternative hypotheses of admixture directionality. Here, we present a 10X depth genome of the 130,000–115,000-year-old subfossil polar bear from the Norwegian Svalbard archipelago, and ten new polar and brown bear genomes from contemporary zones of overlap in northern Alaska where the species may have come into increasing contact due to recent climatic changes. Here, we use a more complete genome from this ancient polar bear and an extended sampling of extant bear populations, comparing 65 polar bear and brown bear genomes from throughout their geographic ranges to better characterize evolutionary splits and admixture between polar and brown bears.

## Results

### Assembly of an ancient polar bear genome and new modern bear genomes

Using a strategy combining multiple sequencing library construction methods and both Ion Torrent and Illumina sequencing platforms, we assembled a 10.11X depth and 97% width of coverage ancient subfossil polar bear genome from the Svalbard archipelago (hereafter denoted as APB). We mapped a total of almost 5 billion sequence reads to a chromosome-length assembly (19, 20) based on the polar bear draft genome UrsMar_1.0 (GCF_000687225.1) (9) (*SI Appendix*, Table S1 and S2). Postmortem damage profiles of the mapped reads from each library sequenced identified typical patterns of nucleotide misincorporations at the reads ends (*SI Appendix*, Text S4, Fig. S2), as expected with degraded ancient DNA. To confirm the authenticity of the APB sequence data, we also assembled mitogenomes from each of the six separate libraries constructed for APB and added them to an alignment of mitogenomes of all modern bear samples analyzed as part of this study. Mitogenome data from all individual APB libraries grouped with the previously published mitogenome from the same ancient polar bear specimen (3) in a position sister to all modern polar bears (*SI Appendix*, Fig. S3), demonstrating the endogenous nature of all new APB libraries.

To expand the geographic sampling of extant bear genomes (Table 1, *SI Appendix*, Fig. S1), and evaluate contemporary admixture among polar bears and brown bears, we also generated ten new modern bear genomes of 9–28X sequence depth coverage (*SI Appendix*, Table S1). Combining these new genomes with previously sequenced genomes of American black (*Ursus americanus*), brown, and polar bears (4, 9) provided 65 modern bear genomes (*SI Appendix*, Table S3), representing all major contemporary brown bear and polar bear maternal lineages. We aligned the sequence reads from these genomes to the same polar bear draft genome as the APB and called over 90 million nuclear single nucleotide polymorphism (SNP) genotypes that were filtered and prepared for downstream analyses (*SI Appendix*, Table S4 and S5).

**Table 1.** Samples, locality, and genome coverage for the 11 new genomes generated for this study. For a complete list of samples analyzed, see *SI Appendix*, Table S1.

| Species | Geographic locality | Sample ID<br>(population ID) | Average width<br>coverage | Average depth<br>coverage |
| --- | --- | --- | --- | --- |
| *U. maritimus* | Poolepynten, Svalbard | APB (APB) | 0.97 | 10.11 |
| *U. maritimus* | Chukchi Sea, AK | AK017 (AK) | 0.97 | 9.47 |
| *U. maritimus* | S. Beaufort Sea, AK | AK034 (AK) | 0.98 | 28.36 |
| *U. arctos* | Seward Peninsula, AK | BB020 (BB) | 0.97 | 9.26 |
| *U. arctos* | North Slope, AK | BB034 (BB) | 0.97 | 8.57 |
| *U. arctos* | North Slope, AK | BB037 (BB) | 0.97 | 9.19 |
| *U. arctos* | North Slope, AK | BB049 (BB) | 0.97 | 9.43 |
| *U. arctos* | North Slope, AK | BB059 (BB) | 0.97 | 9.14 |
| *U. arctos* | Anchorage, AK | EB027 (BB) | 0.97 | 9.59 |
| *U. arctos* | Douglas River, AK | WB039 (BB) | 0.97 | 8.76 |
| *U. arctos* | Yellowstone NP | CON001 (YB) | 0.97 | 9.32 |

### Genetic relationships among brown and polar bears highlight mitochondrial-nuclear discordance

Phylogenetic analysis of assembled mitochondrial genomes (Fig. 1A) confirms previously reported findings of a close maternal relationship between polar bears and Alexander Archipelago (ABC) brown bears, a clade that is in turn sister to a brown bear from Finland (representing European clade 1) (15, 21). Sister to this larger lineage are the remaining brown bear individuals from three main matrilines: a lineage comprising individuals from Yellowstone and Glacier National Parks (North American clade 4 bears) and two sister lineages comprising Eurasian and Alaskan bears (subclades 3a and 3b), including bears from western Alaska (BB034, BB049, BB059, BB020, WB039, and GRZ) and eastern Alaska (BB037 and EB027), respectively (*SI Appendix*, Fig. S1A). The North Slope of Alaska encompasses bears with either eastern Alaskan (BB037) or western Alaskan (BB034, BB049, BB059) mitochondrial haplotypes, indicating that North Slope brown bears contain considerable matrilineal diversity.

**Figure 1.**
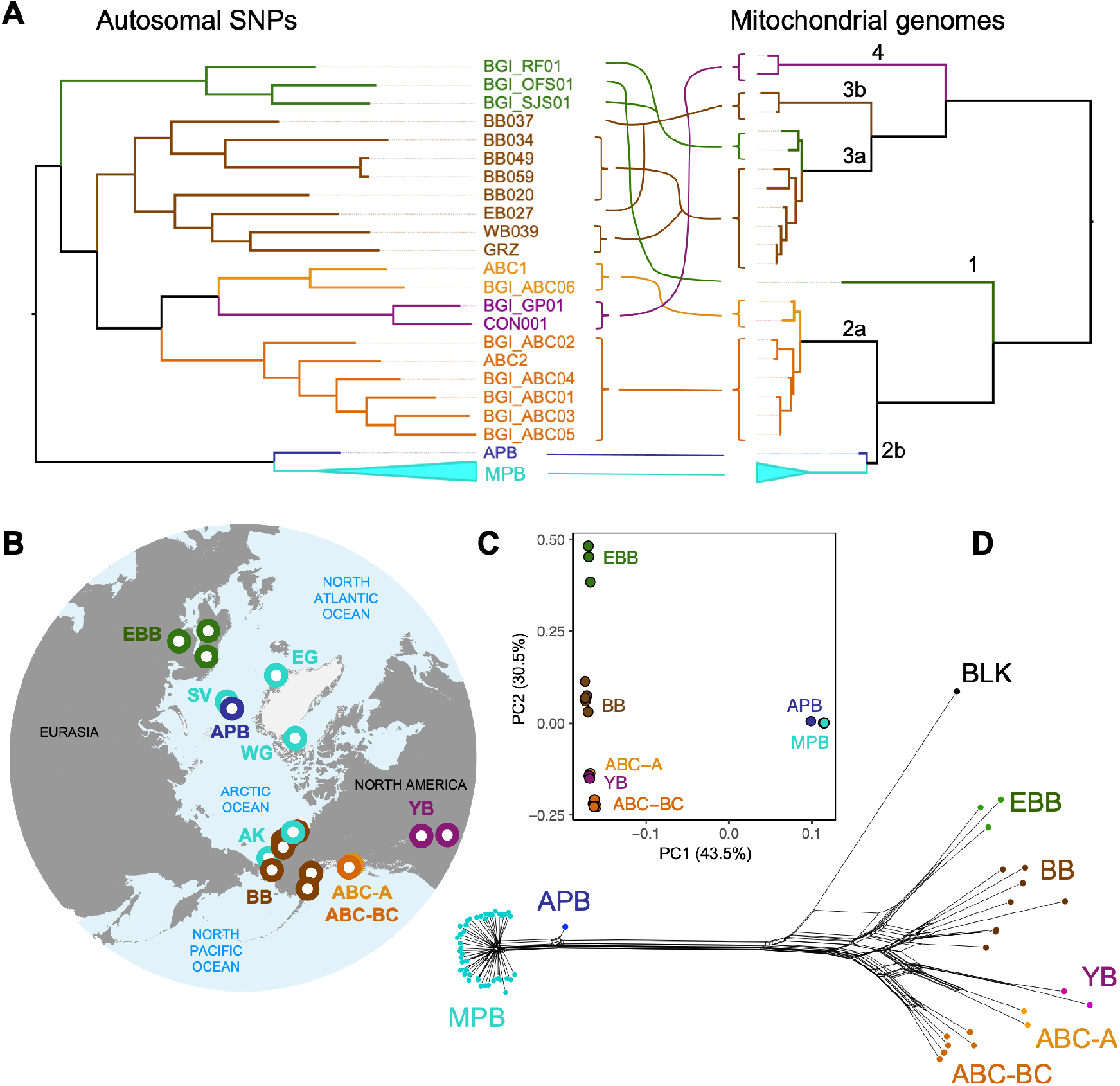
(*A*) Maximum likelihood phylogenetic trees based on autosomal SNPs (left) and complete mitochondrial genomes, with maternal clade names indicated above branches (right). Incongruences between the two phylogenetic topologies are highlighted with colored lines. (*B*) Map showing localities of bear population groupings included in analyses: Alaskan brown bears (BB; brown; BB020, BB034, BB037, BB049, BB059, EB027, WB039, GRZ), continental North American brown bears (YB; CON001, BGI_GP01), Admiralty brown bears (ABC-A; yellow; ABC1, BGI_ABC06), Baranof and Chichagof brown bears (ABC-BC; orange; ABC2, BGI_01, BGI_02, BGI_03, BGI_04, BGI_05), European brown bears (EBB; green; BGI_RF01, BGI_OFS01, BGI_SJS01), and polar bears (dark blue for the ancient polar bear, APB, and light blue for modern polar bears, MPB, from the Svalbard Archipelago, SV, East Greenland, EG, West Greenland, WG, and Alaska, AK). See also *SI Appendix*, Table S1 for provenance of each individual and *SI Appendix*, Fig. S1A, for a map of the geographic localities of the Alaskan bears new to this study. (*C*) principal component analysis of brown and polar bear genomes with genome coverage >8x (DS3). (*D*) Neighbor-Net phylogenetic network based on autosomal SNPs.

The nuclear autosomal SNP phylogenetic tree (Fig. 1A) is highly incongruent with the mitochondrial phylogeny in that polar and brown bears comprise two distinct nuclear lineages, as previously demonstrated (4, 9, 14). In the autosomal tree, all brown bears form a strongly supported clade, with the European brown bears grouping together (hereafter referred to as EBB bears; Fig. 1B), and this clade sister to a lineage containing the remaining brown bears. This latter lineage includes a monophyletic group of Alaskan brown bears (BB bears) that is in turn sister to the Yellowstone and Glacier National Parks brown bears (YB bears) plus the ABC brown bears. Interestingly, the ABC brown bears are paraphyletic, with two individuals from Admiralty Island (ABC-A bears) sister to the YB bears, while the bears from Baranof and Chichagof Islands (ABC-BC bears) form a monophyletic group.

These phylogenetic groupings are recapitulated by principal component analysis (PCA; Fig. 1C), wherein the brown bears form these same five clusters: EBB, BB, YB, ABC-A, and ABC-BC (but note that the PCA includes only individuals with genomes of > 8X coverage; see *SI Appendix*, Table S3). The western and eastern Alaskan brown bear matrilines are not evident from the autosomal phylogenetic tree nor the PCA. It is noteworthy that two Alaskan brown bears (BB049 and BB059) are extremely close relatives in both phylogenetic analysis and PCA, confirming the purported parent-child relationship between a light-colored cub (BB059) and its brown mother (BB049) (*SI Appendix*, Text S3; see also *f*_3_-analysis, below).

Although phylogenetic relationships among modern polar bears are poorly resolved, reflecting their relatively low genetic diversity (4), some mito-nuclear incongruence within polar bears is evident. In the nuclear autosomal phylogenetic tree, the groupings largely follow geographic locality (*SI Appendix*, Fig. S4), and this pattern is largely captured by PCA, although some Svalbard bears appear as outliers (*SI Appendix*, Fig. S6D). On the other hand, the maternal relationships are poorly resolved and do not recapture these general geographic relationships (*SI Appendix*, Fig. S5). In all PCA and phylogenetic analyses, the ancient polar bear is clearly genetically distinct from all modern polar bears (MPB), with the mitochondrial DNA and nuclear autosomal phylogenies supporting a sister-group relationship to all modern polar bear specimens.

To provide a first visualization of discordance among the SNP data that might stem from past admixture among bear species and interspecific populations, we employed the Neighbor-Net approach (22) to generate a distance-based phylogenetic network applied to the same SNP data used for phylogenetic tree reconstruction (Fig. 1D). Character incongruences that are manifest as extra edges in such networks (beyond a perfectly bifurcating tree) have been variously interpreted by other investigators to reflect admixture and/or incomplete lineage sorting phenomena (23, 24). Immediately apparent are three principal findings: (1) modern polar bears form a highly distinct group with only few extra edges separating them from APB and other bear species; (2) brown bear groups are strongly webbed by network edges; and (3) American black bear (BLK) is itself connected through extra edges to brown bears, with a major connection to EBB bears. The impression from the network is one of considerable allele sharing among brown bear groups, as well as between polar bear, brown bear, and American black bear species.

We next explored the level of genomic diversity among the bear genomes. Significantly lower and uniform levels of heterozygosity and nucleotide diversity was found among modern polar bears compared to brown bears (Fig. 2A,B; *SI Appendix*, Fig. S7A,B), as previously reported (9). Interestingly, the ancient polar bear exhibited higher heterozygosity and nucleotide diversity than any of the modern polar bears. Although many alleles unique to the ancient polar bear (*SI Appendix*, Fig. S7C) largely contributed to this difference (*SI Appendix*, Fig. S7A), when SNPs were filtered for private alleles, the level of genetic diversity was still slightly higher in the APB compared to modern polar bears (Fig. 2A; *SI Appendix*, Fig. S8). Among brown bears, the ABC and YB bears (the Yellowstone brown bear, CON001, in particular; *SI Appendix*, Fig. S7) exhibited the lowest levels of genetic diversity. Population differentiation identified from clusters in the PCA analysis (Fig. 1B) demonstrated low differentiation among modern polar bear clusters (*SI Appendix*, Fig. S9; mean weighted F_ST_ = 0.031–0.054) as compared to among brown bear clusters (mean weighted F_ST_ = 0.127–0.276).

**Figure 2.**
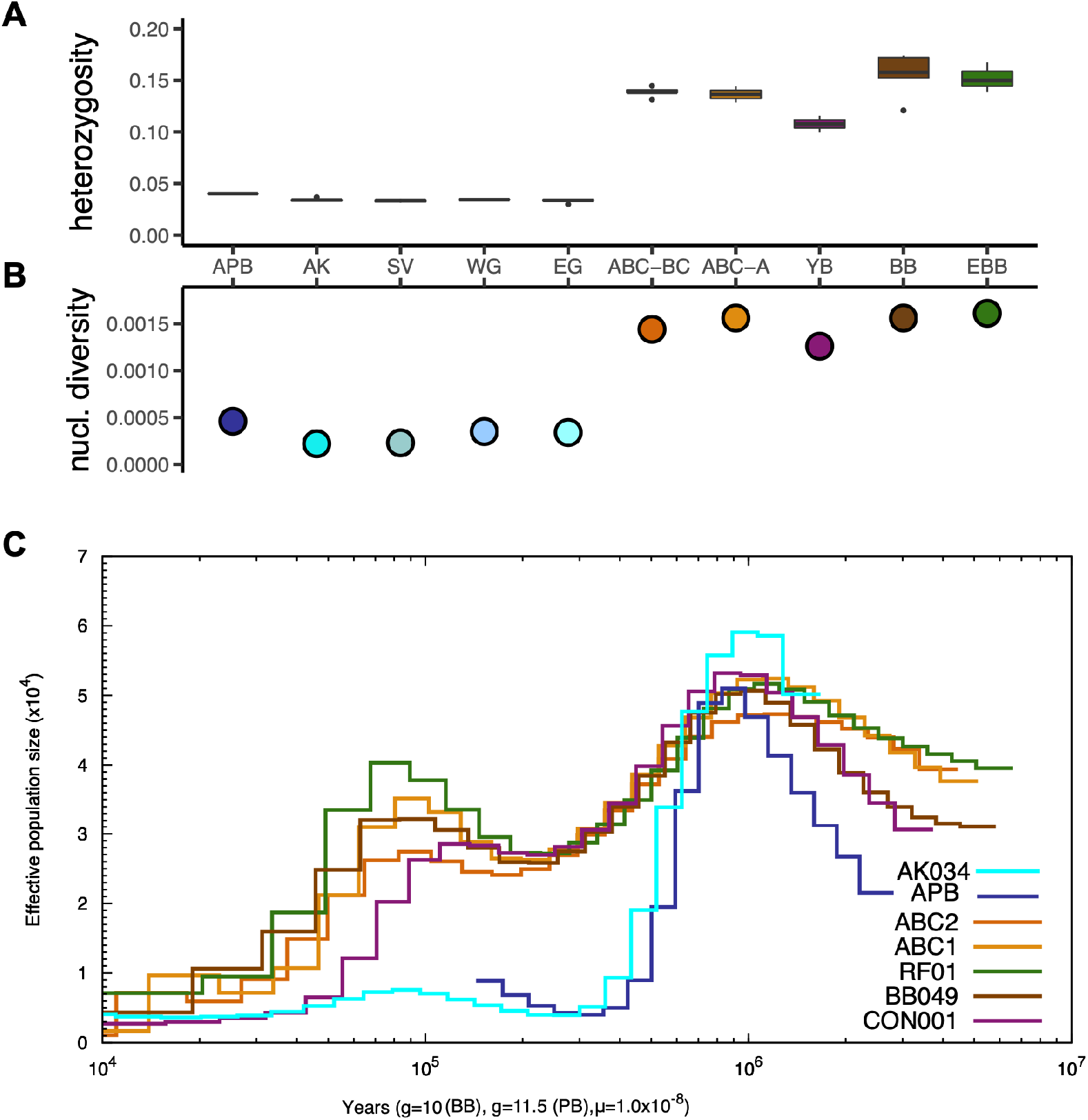
(*A*) Autosomal heterozygosity frequencies and (*B*) and nucleotide diversity, pi, across polar and brown bear populations (dataset DS2). Acronyms of population groupings follow the description in Fig. 1. (*C*) Estimates of effective population size over time shown for one representative individual from each of the brown bear and polar bear populations: AK034 (Alaskan polar bear), APB (the ancient polar bear), ABC2 (Chichagof brown bear), ABC1 (Admiralty brown bear), RF01 (European brown bear), BB049 (Alaskan brown bear), CON001 (Yellowstone brown bear). For provenance of each individual see *SI Appendix*, Table S1.

### Population demographic histories of highly inbred ancient and modern polar bears differ substantially from those of brown bears

We used pairwise sequentially Markovian coalescent (PSMC) analysis (25) to infer the population demographic history for all bear individuals, including the ancient polar bear. As has been demonstrated before (4, 9), we also found evidence for a small long-term effective population size (Ne) in extant polar bears following a dramatic and steep decline, possibly about 500,000 years ago (Fig. 2C). Using an average generation time for polar bear (g = 11.5 years) and brown bear (g = 10 years), following comprehensive assessments of generation lengths in the two species (26, 27), provided comparable demographic histories to previously reported results based on a generation time of g = 10 for all bears (see *SI Appendix*, Fig. S10). A similar sharp decline in Ne was found when demographic history was inferred for the ancient polar bear, but interestingly, a modest increase in Ne for both the ancient as well as extant polar bears was apparent before the APB’s demise about 120,000 years ago (Fig. 2C). It is possible that this Ne increase represents a population expansion, or perhaps a slight increase in heterozygosity following interbreeding with brown bear (see below). In contrast to polar bears, the past decline in Ne among brown bears was more gradual and interrupted by a significant population expansion about 100,000 to 150,000 years ago (*SI Appendix*, Fig. S10C). This marked difference in demographic histories between polar bear and brown bear was also reflected in the reduced heterozygous genetic background observed among polar bears, in particular modern polar bears, compared to brown bears (Fig. 2A and *SI Appendix*, Fig. S7). It is noteworthy that the demographic curves for polar bears display a slight “shift” towards modern times compared to brown bear demographies. Highly heterozygous regions coalesce further back in time in coalescent modeling. Therefore, this shift, implying much more recent coalescence throughout the genome in polar bears than brown bears, could have resulted from phenomena that reduce genome-level heterozygosity, such as high levels of inbreeding, as has been demonstrated in plant systems (28).

We also applied the SMC++ method, which couples the genealogical process for a given diploid individual with the allele frequency information in a collection of other individuals, providing higher Ne resolution in the recent past (29). Similar to that observed with PSMC, we see an ancient sharp decline in polar bear population size (*SI Appendix*, Fig. S11). We also observed a gradual decline in Ne over the past 10^5^ generations among brown bears (*SI Appendix*, Fig. S11), particularly in the continental (YB) and ABC brown bears, likely reflecting differences in heterozygosity levels among these brown bears versus European and mainland Alaskan populations. Shortly after 10^4^ generations, there is an increase in brown bear Ne, while the polar bear Ne decreases, suggesting differential responses to environmental perturbation (*SI Appendix*, Fig. S11C).

### Estimates for the polar-brown bear split time range to 1.6 million years ago

To estimate the average coalescence time between polar bears and brown bears, we followed the strategy applied in (30). Exploring the alleles of each population BLK, EBB, and APB, we denoted N_X_ by the number of SNPs where the allele from population X differs from the two others. Assuming mutations occur at similar rate *ρ* per year on each lineage, we obtain N_EBB_ = N_APB_ + A\**ρ*, where A is 115,000–130,000 years. From the data N_APB_ = 368,285 and N_EBB_ = 402,509, the genetic divergence time between brown bears and polar bears, N_EBB_/*ρ*, is estimated to be 1.3–1.6 ma. As a genetic divergence time, this is an upper bound for the population divergence time. We note that this split estimate is consistent with the coalescence time estimated by PSMC analysis, which inferred that the population divergence between brown bears and polar bears occurred over one million years ago (*SI Appendix*, Fig. S10).

We also estimated split times for polar bear and brown bear populations applying the SMC++ method, which analyzes pairs of populations to infer divergence times jointly with population size histories (29). Again, the populations were identified from the clusters in the PCA analysis (Fig. 1B). The ancient polar bear sample was excluded from this analysis, because SMC++ was not able to incorporate an age from an ancient sample. Reflecting the cladistic progression of our populations (*SI Appendix*, Fig. S4), the youngest split time was between the PB and AK modern polar bear populations, followed by the split between the ABC-A and ABC-BC brown bear populations (~11.8 ka), the ABC brown bear split from the mainland Alaskan brown bears (28–36 ka), and the continental (YB) brown bear split from the Alaskan bears (43–46 ka) (*SI Appendix*, Fig. S12 and Table S6). The split time between North American brown bear populations and EBBs ranged from 95 ka to 150 ka. The split time between all brown bear populations and modern polar bears was estimated to be ~264 ka, much younger than the 1.3–1.6 ma split estimate above. Importantly, the clean-split model currently implemented in SMC++ (31) assumes that two subpopulations are descended from a common ancestral population, with no gene flow occurring more recently than the subpopulation split. Hence, in the case of polar bears and brown bears, which have clearly demonstrated gene flow following their divergence (see below), the split time estimated using SMC++ is most likely underestimated, possibly associated instead with cessation of gene flow between the two lineages. Likewise, the ~11.8 ka split time estimate for ABC-A and ABC-BC brown bears more likely reflects interpopulational gene flow termination accompanying fragmentation of Alexander Archipelago landmasses by sea-level rise at the end of the last Ice Age.

### Polar bear genomes preserve the strongest evidence for past admixture with brown bears

Because we do not expect simple, bifurcating patterns to fully describe bear population interrelationships, we applied multiple measures to estimate genome-wide admixture, some similar to those used for previous work (8, 9, 32), but now including more brown bear genomes from North America and an ancient, and higher-quality, polar bear genome to provide a better temporal framework.

#### ADMIXTURE analysis suggests 2% shared ancestry between the ancient polar bear and brown bears

First, we performed a model-based clustering analysis employing the program ADMIXTURE, which estimates ancestry from large autosomal SNP genotype datasets (33). Although no admixture was detected between modern polar bears and brown bears at *K* = 4, which was determined by cross validation to be the optimal number of hypothetical ancestral source populations, slight shared ancestry (~2%) with brown bears from mainland Alaska and Europe was found in the ancient polar bear (Fig. 3A). This same level of shared ancestry between APB and brown bears was also recovered with *K* = 2 and *K* = 3 (Fig. 3A and *SI Appendix*, Fig. S13). Inside polar bears and brown bears, considerable population structure was apparent. For example, polar bears sort into several ancestral groups at *K* = 5 to 7, suggesting significant population structure among polar bears that largely corresponds with clusters observed in PCA (*SI Appendix*, Fig. S6D).

**Figure 3.**
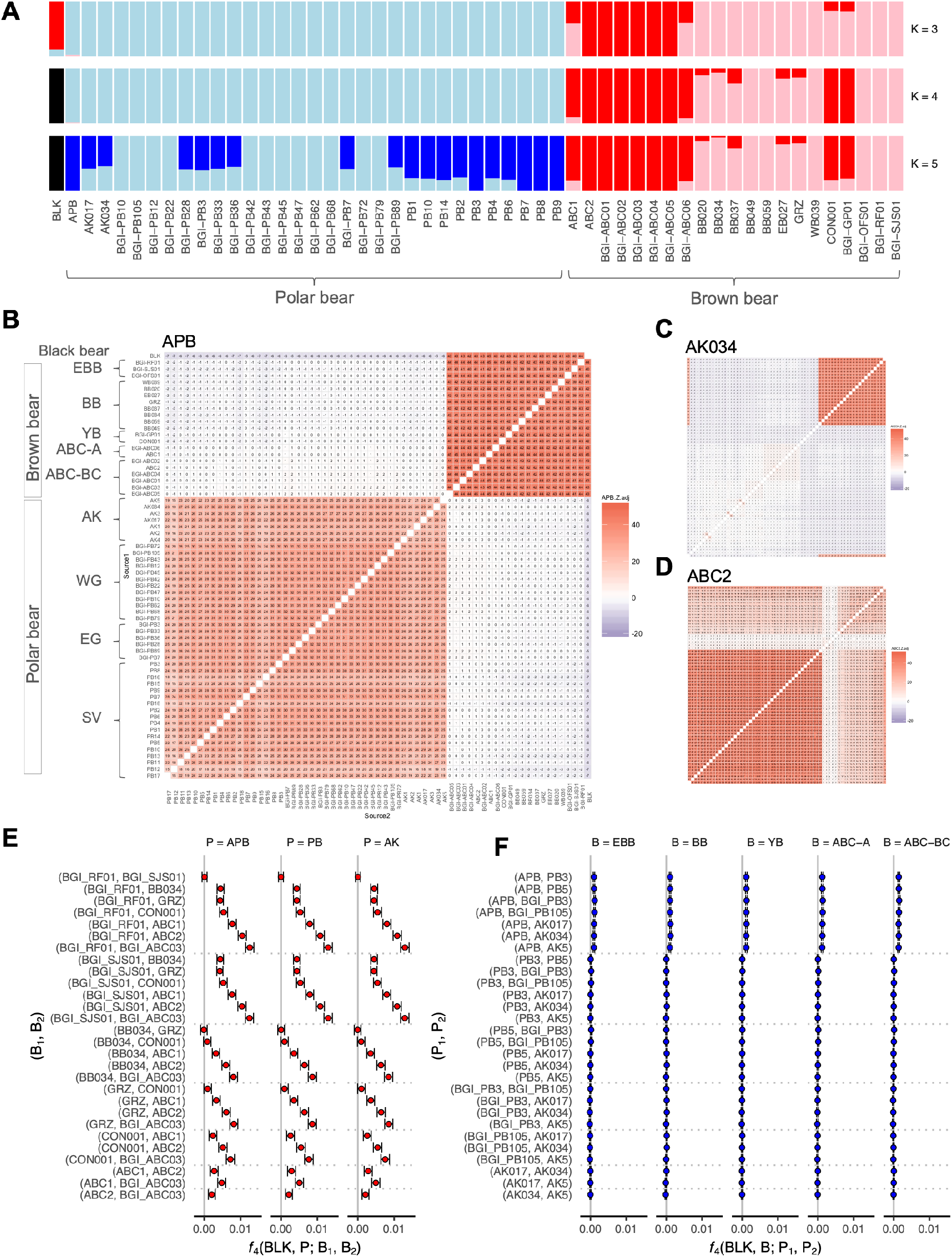
(*A*) ADMIXTURE analysis of brown bear, polar bear, and American black bear genomes with coverage >8x (ancestral population clusters, K = 3-5, is shown). (*B*) *f*_3_ statistics results (adjusted Z scores) showing target population APB, (*C*) AK034 (Alaskan polar bear), and (*D*) ABC2 (Chichagof brown bear). The bear group acronyms follow the descriptions in Fig. 1. *f*_4_ statistics showing *f*_4_ values and their 95% confidence intervals of (*E*) *f*_4_(BLK, X; B_1_, B_2_) and (*F*) *f*_4_(BLK, X; P_1_, P_2_).

#### f_3_-statistics show evidence for interspecies admixture in polar bears only

Next, we used the *f*_3_-statistic, *f*_3_(C; A, B), to provide definitive evidence of admixture (34). We calculated *f*_3_ for all possible combinations of bear individuals and populations, with populations following clusters identified with PCA, to consider potential introgression among polar, brown and American black bear in all directions (*SI Appendix*, Fig. S14). We conducted the *f*_3_-statistics using both all substitutions and transversion substitutions only. At the population level, when all substitutions were considered, with APB as the target and black bear as one source and modern polar bear the other, *f*_3_ was significantly negative, which is indicative of APB being admixed (*SI Appendix*, Fig. S14). This pattern was repeated at the individual bear level with APB as a target (Fig. 3B and *SI Appendix*, Fig. S15). When only transversions were included, no three-way comparisons had significantly negative *f*_3_ values at the population level (*SI Appendix*, Fig. S14), while they were only marginally negative at the individual level (*SI Appendix*, Fig. S16), suggesting that this signal may not be robust or that power was lost with the reduced number of SNPs in the transversion-only dataset. All other *f*_3_ results at the individual level exhibited similar patterns for both datasets; hence, only results obtained when all substitutions are considered are further mentioned (*SI Appendix*, Fig. S15).

Significantly negative *f*_3_(C; A, B) values were observed for multiple polar bear individuals. In some cases, they merely indicated close relationships (population sharing) between the target C and one of the sources, because the presence of segments that are identical by descent will inflict a downwards deviation in the *f*_3_-statistic estimators (for the exact derivation of the deviation on the level of single individuals, see *SI Appendix*, Text S15). Such cases included polar bear individuals from the Svalbard Archipelago (PB3/PB8, PB5/PB14, and PB7/PB9), where the highly significantly negative *f_3_* values may indicate familial relationships or inbreeding among Svalbard Archipelago polar bears (35). West Greenland polar bears also appeared to be close relatives of one another, particularly BGI-PB47/BGI-PB10, although less so compared to the aforementioned Svalbard individuals. Some Alaskan polar bear individuals also exhibited admixture with brown bear, particularly AK034, a female from the Southern Beaufort Sea that exhibited many significantly negative *f*_3_ values when one source included a brown bear individual (Fig. 3C). The only brown bear individuals that exhibited significantly negative *f*_3_ values were BB059 and BB049, and only when one of these individuals represented the target individual and the other individual the source population (*SI Appendix*, Fig. S15), confirming their purported familial relationship suggested by kinship analyses conducted using microsatellite loci (*SI Appendix*, SI Text S3). Some of the ABC bears from Baranof and Chichagof also appeared to be close relatives, e.g., BGI-ABC01 and BGI-ABC05. Importantly, however, no other brown bears, including ABC (Fig. 3D) and mainland Alaska brown bears (*SI Appendix*, Fig. S15), exhibited negative *f_3_* values from any source combinations. Although *f*_3_ may be positive under some admixture scenarios (34, 36), these results suggest that polar bears are the only taxon still containing alleles obtained through admixture with other bear species.

#### The f_4_-statistics demonstrate gradually increasing allele sharing from EBB to ABC brown bears

The *f*_4_-statistic (closely related to the *D*-statistic) is a four-taxon test of admixture that has become an important tool for estimating gene flow in population genomics (30, 34, 37). We evaluated levels of shared drift by computing *f*_4_(BLK, P; B_1_, B_2_) (Fig. 3E) and *f*_4_(BLK, B; P_1_, P_2_) (Fig. 3F), where P and B are populations of polar bears and brown bears, respectively, Bi are brown bear individuals, and Pi polar bear individuals (see *SI Appendix*, Table S1). The significantly positive values of *f*_4_(BLK, P; B_1_, B_2_) (Fig. 3E, *SI Appendix*, Table S7) indicate gene flow between brown bears and all polar bears, including APB, with an increasing trend from EBB to ABC-BC. It is important to note that this EBB to ABC-BC *f*_4_ trend does not necessarily require multiple gene flow events into polar bear, as suggested by other investigators; rather, it may simply represent a gradient of relatedness (drift) extending through brown bear cladogenesis that would exist regardless of any admixture events. Our rationale for this is illustrated in the next section. Further, in *f*_4_(BLK, B; P_1_, P_2_) tests (Fig. 3F, *SI Appendix*, Table S8) with APB included as one of the polar bear populations (P_1_), the *f*_4_ values were slightly significantly positive, with an increase as B goes from EBB to ABC-BC, although much less pronounced than before. When only modern polar bears were evaluated instead, values did not significantly deviate from zero. These results indicate some admixture between brown bears and modern polar bears after the APB-PB split. On the other hand, modern polar bears appear highly homogenous, with the gene flow between brown bears and modern polar bears (Fig. 3E) affecting all modern polar bear individuals equally, thereby suggesting its occurrence before they shared a common ancestor.

#### Lengths of introgressed segments are less than 1 Mb in polar bears, consistent with admixture being ancient

The *f*_2_-statistic measures the amount of drift separating two populations (34, 38). To search for fragments where ABC brown bears and polar bears might resemble each other as a signature of admixture, we examined genetic distances within 50 kb blocks between ABC bears and modern polar bears versus between ABC bears and the ancient polar bear (*SI Appendix*, Text S17). The two distances, as well as their difference, are plotted in *SI Appendix*, Fig. S17, and 18 potentially introgressed regions are highlighted in *SI Appendix*, Fig. S18. These segments are less than 1 Mb in length, with most of them only ~250 kb. This result is consistent with a previous study that estimated any admixture between the two species must have occurred at least hundreds of generations (or thousands of years) ago (9).

### The predominant direction of gene flow was from brown bear into polar bear

#### f_4_-ratio estimation is inadequate to infer gene flow direction among brown and polar bears

The current paradigm of unidirectional gene flow from polar bears into brown bears is largely based on *f*_4_-ratio estimation (30, 34), which has previously been applied to study the direction and proportion of gene flow among brown bears and polar bears (7, 8). That work proposed that varying levels of admixture among brown bears are best explained via multiple gene flow events from polar bears into brown bears. We therefore sought to replicate and reassess these results using our data (*SI Appendix*, Text S18). Similar to previous observations of 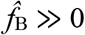 and 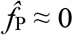 (8), we estimated 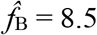 and 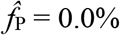. Importantly, however, this approach requires the assumption that some polar bear populations (here AK) are unadmixed, while others (here PB) are admixed, which violates our finding that *f*_4_(BLK, ABC-BC; P_1_, P_2_) = 0 (Fig. 3F), irrespective of which modern polar bear population is included. In any case, ancient gene flow in either direction between ABC-BC and the ancestors of all modern polar bears fits the observation 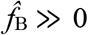 and 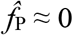 just as well as potential modern gene flow from polar bears into ABC-BC (*SI Appendix*, Fig. S20, see also Text S18). Using APB as the polar bear sister population instead, in case it may better represent an unadmixed polar bear, these values are 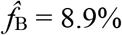 and 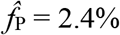. However, the assumptions of *f*_4_-ratio estimation are still not met, because we already detected that APB is also involved in brown bear admixture, which occurred before the ancient-modern polar bear divergence (Fig. 3E). Even if APB was unadmixed, the two non-zero numbers 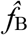 and 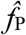 could not be interpreted as admixture proportions, because each contradicts the assumptions made when computing the other (see *SI Appendix*, Fig. S19). However, under the current paradigm of unidirectional gene flow from polar bears into ABC bears (either before or after the APB-PB split) (6–8), we should nevertheless have expected 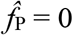 with APB as well. In summary, we determine that (*i*) recent gene flow only into brown bear populations is not the only possible explanation for the *f*_4_-ratio observations, as the fractions 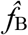 and 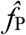 represent admixture proportions under assumptions that may not be met, and (ii) various ancient gene flow scenarios fit the observations just as well (*SI Appendix*, Fig. S20). Hence, *f*_4_-ratio analysis is inappropriate for our sample. As such, we instead explored extensions of *f*_2_ and *f*_4_-statistics in admixture graph statistical fitting.

#### Graph fitting applied selectively indicates ancient bidirectional gene flow

Admixture graph fitting provides a rigorous test for whether a proposed evolutionary model fits the data (34, 39). Whereas the *f*- and D-statistics on four populations usually detect only the *presence* of admixture, introducing a fifth population is informative on the *direction* of gene flow (34). With the aim of assessing the timing of gene flow, we also added a sixth population, APB, and studied the six populations BLK, EBB, BB, ABC-BC, PB, and APB. We used the admixturegraph package (39) to fit all 105 trees with six populations to 4319 *f*_2_-data sets computed within 500 kb windows (Fig. 4A and *SI Appendix*, Fig. S21 and Text S19). The idea was that gene flow events in different directions between ABC-BC and polar bears could create segments that locally resemble different trees, as may also be expected from incomplete lineage sorting (ILS).

**Figure 4.**
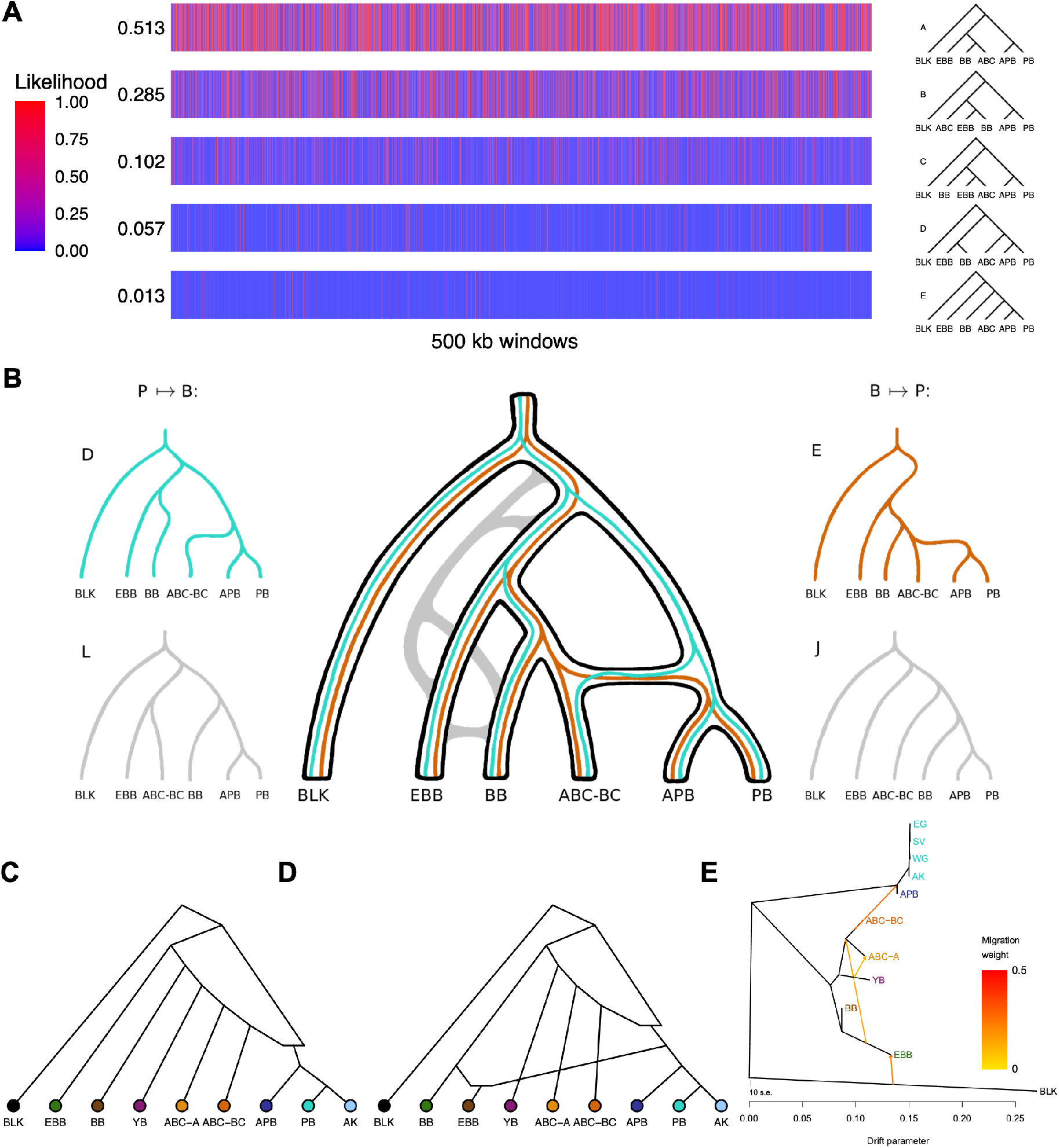
(*A*) The likelihood of the top five best fitting trees among 105 different trees according to admixturegraph analysis using *f*_2_-statistics within 500 kb windows. Values to the left are averages over all the windows. (*B*) Scenarios of gene flow and incomplete lineage sorting among best-fitting trees in (*A*). Best fitting admixture graphs after (*C*) Stage 3 of the *f*_4_-analysis (sum of squared errors, C = 9192) and (*D*) Stage 4 of the *f*_4_-analysis (C = 4055). (*E*) TreeMix showing 4 migration edges for populations of brown and polar bears based on genomes >8x coverage. The bear group acronyms follow the descriptions in Fig. 1.

In most regions, tree A, which recapitulates the expected population relationships, provides the best overall fit (Fig. 4A). Trees B and C, the next best fitting, show rearrangements among brown bear populations, possibly reflecting admixture and/or ILS within brown bears. Trees D and E (Fig. 4B) are the fourth and fifth best fitting trees. In D, drift from the polar bear clade contributes alleles to brown bear phylogeny only via entry into the ABC-BC edge. In E, in contrast, brown bear phylogenetic drift, only from ABC-BC, contributes alleles to polar bear at the edge subtending APB+PB. Hence, gene flow from ancient polar bears predating the PB-APB-split into the ancestors of ABC-BC bears would result in modern ABC-BC bears carrying small segments of DNA that would appear to belong to a sister group of PB+APB rather than a sister group of BB. In such segments, the tree D is likely to fit well. In contrast, gene flow in the opposite direction would result in all polar bears carrying segments that would appear to belong to a sister group of ABC-BC rather than a sister group of all brown bears; here, tree E is likely to fit well. In other words, trees D and E (Fig. 4B) represent gene flow from ancient polar bears (predating the APB-PB split) into ABC-BC brown bears, and the inverse, respectively. Ruling out a simple ILS argument for either pattern of allele sharing, ILS, through its expected symmetry, should favor alternative trees L and J (Fig. 4B) equally often as D and E, whereas the likelihoods of L and J are considerably worse than those of D or E (*SI Appendix*, Fig. S21).

Due to differences in how gene flow, ILS and phylogenetic drift might conspire to impact the likelihood scores of trees D and E, their direct comparison is inapt (see *SI Appendix*, Text S19). Therefore, we conclude, in the absence of other plausible explanations, that bidirectional gene flow between ancestors of all polar bears and ancestors of ABC bears is the most likely evolutionary scenario. Furthermore, we find no evidence for modern gene flow (not involving APB) between these lineages because trees where PB and APB are not sisters fit the data only poorly.

#### Graph fitting applied to all populations shows a preference for gene flow from brown bears into the ancestor of ancient and modern polar bears

Given that gene flow appears to have been bidirectional between ancient polar bears and ancient brown bears and considering that the *f*_4_-ratio estimation is not applicable, we next applied admixturegraph to find models that are consistent with the full set of *f*-statistics. In a manner close to exhaustive, we tested admixture graphs including three additional populations: YB, ABC-A and AK, adding an admixture event to the best fitting trees, and a second admixture event to the best fitting single-admixture graphs (see *SI Appendix*, Text S20). The results showed a clear preference for gene flow from brown bears into polar bears, with the most recurrent feature among well-fitting admixture graphs being an admixture edge from ABC-BC into the ancestors of all polar bears (Fig. 4C and *SI Appendix*, Fig. S22B and S23B). The direction from polar bears into brown bears also fit well among some 1-admixture graphs (*SI Appendix*, Fig. S22B and S23B). The outstanding best fit among the 2-admixture graphs (Fig. 4D and *SI Appendix*, Fig. S22C and S23C) features bidirectional gene flow, including the admixture edge from ABC-BC into the ancestors of all polar bear, as seen in the 1-admixture graph, and an edge from the ancestors of polar bear into EBB. However, because the position of EBB is inconsistent with the species tree, and since there was no evidence for admixture between EBB and polar bears in other analyses, this is likely a result of ILS and/or ghost admixture into EBB, possibly coming from cave bears (as discussed below). The second admixture event in most other bestfitting, 2-admixture graphs, when consistent with the species tree, typically concerned brown bears only. It is worth noting that EBB appeared admixed in many graphs, including the bestfitting graph (Fig. 4D), and therefore its role as a model of an unadmixed brown bear (6, 9, 18) must be reconsidered.

#### TreeMix analyses suggest admixture from ABC-BC bears into the ancestral node of polar bears

Finally, we generated a maximum likelihood drift tree using TreeMix (40) to infer patterns of population splits and mixtures among multiple populations. Although the initial tree with no migration edges largely recapitulated the splits already seen in the RAxML autosomal SNP analysis (*SI Appendix*, Fig. S24), 0.23% of the variance was residual to the model’s fit, i.e., was not captured by the tree (*SI Appendix*, Table S9). Hence, we sequentially added one to five admixture events to the tree (*SI Appendix*, Fig. S24). Several admixture events stood out, particularly admixture from ABC-BC bears into the ancestral node of the polar bear lineage (Fig. 4E), which was the first admixture edge found consistently throughout all TreeMix results. Other admixture events included migration edges among brown bears, e.g., EBB to the ancestor of the ABC brown bears, and admixture edges from outside the brown bear / polar bear lineage into EBB (Fig. 4E) and APB (*SI Appendix*, Fig. S24), respectively. Hence, the TreeMix results are consistent with the admixturegraph results.

## Discussion

It has been established that widespread and rapid global climate changes have occurred at unprecedented scales in recent years (41). Associated with these climate changes are well-documented impacts to the ecologies and life histories of plants and animals (42), including shifts in latitudinal and elevational ranges (43–45), local extinctions (46), and changes in morphologies (47). Furthermore, colonizing species have in some cases been shown to capture local adaption by hybridizing with closely related resident lineages (48). Because hybridization may catalyze adaptive evolutionary change (49), its potential role among the responses to global climate change should not be underestimated.

Although contemporary ranges of polar bears and their lower-latitude closest relatives are discrete across most of the Arctic, latitudinal shifts in their distributions in recent years are likely caused by the altered Arctic environment (50–52). For example, brown bears and American black bears appear to be moving northward into the Canadian Arctic Archipelago (53, 54). Polar bears are increasingly summering in nearshore terrestrial and barrier island habitats in the central Beaufort Sea, likely due to the loss of nearshore Beaufort Sea ice during summer, and possibly facilitated by the presence of fall subsistence-harvested bowhead whale (*Balaena mysticetus*) remains (55, 56). Such increased range overlaps may permit increased interactions between these closely related species, including competition and hybridization.

Despite evidence for brown bear-polar bear hybrids in the Canadian Arctic (53), contemporary hybridization seems sparse and therefore its potential impact limited. For example, recent genotyping and parentage analysis of numerous bears in the western Canadian Arctic Archipelago traced eight hybrid individuals to a single female polar bear who mated with two brown bears (11). These findings suggested that although the evolutionary importance of breakdown of species barriers should not be underestimated, recent hybridization between the two species could merely be caused by uncommon and atypical mating preferences of select individuals. In keeping with this hypothesis, an expansive genetic analysis of a large, circumpolar sample of polar bear subpopulations failed to find genetic signatures of recent hybridization between the two species, suggesting that recently observed hybrids represent localized events (57).

Recent research based on genomic data, however, has pointed to considerable ancient introgressive hybridization between bear lineages (4, 8, 9). The current consensus scenario in the literature for brown and polar bear admixture, referred to as the “population conversion model” (6, 8), involves multiple polar bear introgressions into brown bear lineages, possibly also including extinct Irish brown bears (7). The direction of gene flow has implications for how climate change and range overlap may have influenced adaptive evolution. On the one hand, gene flow from polar bear into brown bear suggests that generalist, boreal predators were the recipients of high-Arctic specialist alleles, with a selective barrier to gene flow possibly acting in the opposite direction (8). The converse, gene flow from brown bear into polar bear, would implicate capture of generalist, boreal-adapted alleles by Arctic specialists known to be highly sensitive to climate change.

Similar to the difficulty in timing the split between the polar and brown bear lineages, however, the complex and highly disparate population histories of the two species complicate resolving their intertwined evolutionary past. We show here that fossil DNA evidence may hold the necessary clues. Our analyses of a genome from a ~120,000-year-old subfossil polar bear and an extended sampling of extant polar and brown bear populations from throughout their geographic range suggest that the two lineages diverged more than one million years ago, which is consistent with earlier estimates from SNP and Y chromosome marker analyses (58, 59), as well as with our comparative PSMC analysis, which inferred that population divergence between brown bears and polar bears occurred over one million years ago. Although some studies have estimated younger split times (18), repeated hybridization may have led to an underestimation of coalescence times (60). We find that gene flow into the polar bear lineage, wherein polar bears also captured a brown bear mitochondrial genome, was likely the predominant admixture direction before most gene flow between the lineages ceased around 200,000 years ago (ka). This latter estimate of a complete split is consistent with the “clean” split time of ~264 ka produced with the SMC++ method employed in this study, our previous estimate based on a coalescence hidden Markov model (4), and a divergence between the maternal lineages of the two species ca. 150,000 years ago (3).

Although several studies demonstrate a consistent signal of gene flow between the polar bear and brown bear lineage, we find that previous use of the *f*_4_-ratio estimation to infer gene flow direction between polar and brown bears (8) has been inadequate, and even inappropriate by its required use of unadmixed populations. Instead, we find that admixture graph fitting, using the methods of admixturegraph and TreeMix analyses, favors predominant gene flow into the polar bear lineage from ancestors of Alexander Archipelago brown bears, whose matriline likely was once more geographically widespread, at least ranging also to Haida Gwaii and interior Alaska (61). This admixture would have occurred before the split between the ~120,000-year-old polar bear and modern polar bear, an inference that is supported by negative *f*_3_ values observed for multiple polar bear individuals, including APB, but no definitive evidence for admixed brown bear individuals. It is also the most parsimonious explanation for the gradually increasing positive *f*_4_ values from EBB to ABC brown bears, i.e., gene flow to polar bear (including the ancient polar bear) from a relative of ABC brown bears, and a gradient of drift paths through their phylogenetic tree, extending from ABC to EBB brown bears and the black bear outgroup. It would require multiple, less-parsimonious gene flow events from polar bears into brown bears to fit these gradually increasing *f*_4_ trends. Furthermore, a brown bear into polar bear principal directionality is entirely consistent with a scenario wherein a brown bear mitochondrial genome was captured by polar bears, reconciling the highly paraphyletic nature of brown bear maternal lineages (3, 16, 17). Importantly, though, our admixture graph fitting analyses also indicate significant gene flow from polar bears into brown bears, and gene flow into brown bears from a population ancestral to both brown bears and polar bears. Our latter finding supports previous reports of gene flow between brown bears and cave bears (62, 63) as well as gene flow involving American black bears (64). Therefore, despite a predominant pattern of gene flow from brown bears into polar bears, it seems likely that the true history of gene flow between these species has been multidirectional, as has been recognized recently for admixture patterns between modern and archaic humans (65).

Our study supports, if not an inverted paradigm shift in the current understanding of gene flow between these bear species, then a new emphasis on its complexity and multidimensionality. These new perspectives may have relevance for our understanding of potential adaptive responses to climate change. Our data suggest that following the divergence between ancestors of brown and polar bears, introgression events between these species predominantly involved gene flow *into* the Arctic lineage from ancestors of extant brown bears, possibly facilitating the capture of novel genes by Arctic specialists (polar bears) from colonizing boreal brown bear generalists. Although there is likely strong purifying selective pressure on polar bear phenotypic features adapted to extreme Arctic life, novel, heritable traits transferred from brown to polar bears could have become selectively advantageous during certain past periods of climatic change. Complete clarity on admixture scenarios is confounded by a complex brown bear phylogeographic history comprising distinct, lineage-specific geographic expansions of brown bears into the New World. Insight into the potential adaptive importance of ancient polar-brown bear admixture could come from genomic regions where modern polar bear alleles are more similar to those of brown bears than they are to the ancient polar bears predating any admixture events. However, such inference of any potential genome-wide adaptive signals, as well as proper modeling and timing of admixture events and demographic expansions and contractions, which in turn will better inform their correlation with events during Earth history, would require the collection of more complete ancient DNA data than presented here, including from unadmixed ancient polar bear remains.

What can our genomic findings contribute to understanding how future climate changes might impact polar bears? Given the evidence for past and current interbreeding between polar bear and brown bear species, hybridization is likely to have been an important element in their evolutionary history, and presumably it may also have an impact in the species’ response to future climate change. It is expected that polar and brown bears will come into more frequent contact due to the loss of sea ice, potentially providing increased opportunities for interbreeding. Importantly, however, the selective pressures incurred by habitat loss and other climate-related impacts will most likely outweigh any potential for adaptive evolutionary change catalyzed by hybridization. The current fragmentation of sea ice habitat is predicted to reduce gene flow among polar bear populations, resulting in increased local inbreeding and overall diversity loss (35). The marked differences between brown and polar bear population histories and genetic diversities, wherein polar bears show the signature of an ancient steep decline in population size with adverse effects to their genetic diversity, may be a testament to the impact of similar responses in the past.

## Materials and Methods

The ancient polar bear DNA was extracted from a canine in a cleanroom facility dedicated to ancient DNA work at the University at Buffalo (UB). Libraries were prepared and sequenced at Daicel Arbor Biosciences and Nanyang Technological University (NTU). DNA was extracted from tissue and blood samples of modern Alaskan bears at UB and sequenced at NTU. In addition to standard quality control procedures, the sequence reads of the ancient sample were end trimmed to alleviate cytosine deamination. For a detailed description of the sample collection, DNA extraction, sequencing, mapping, and SNP calling, as well as all genome evolution analyses, we refer to the *SI Appendix, SI Materials and Methods*.

## Data Availability

Genomic data have been deposited in NCBI (new mitogenome accessions OM732473-OM732482 and bioproject ID PRJNA804505).

## ACKNOWLEDGMENTS

The authors are grateful to Todd Attwood and Karyn Rode (U.S. Geological Survey, Alaska Science Center) for providing polar bear tissue samples. This work was supported by the National Fish and Wildlife Foundation and National Science Foundation (awards no. 1556565 and 1854550 to C.L.), Alaska Department of Fish and Game (to S.F. and R.T.S.), and U.S. Geological Survey’s Changing Arctic Ecosystems (to S.L.T.), which is supported by funding from the Wildlife Program of the USGS Ecosystem Mission Area. Mention of trade names or commercial products does not constitute endorsement or recommendation for use by the U.S. Government.

