## Supplemental Text and Figures for "Insights into bear evolution from a Pleistocene polar bear genome"

**Supplementary information for**
### Insights into bear evolution from a Pleistocene polar bear genome

Tianying Lan[a,b,1], Kalle Leppälä[c,1,2], Crystal Tomlin[a], Sandra L. Talbot[d], George K. Sage[d], Sean D. Farley[e], Richard T. Shideler[f], Lutz Bachmann[g], Øystein Wiig[g], Victor A. Albert[a], Jarkko Salojärvi[h,i], Thomas Mailund[j], Daniela I. Drautz-Moses[k], Stephan C. Schuster[k], Luis Herrera-Estrella[l,m,2], Charlotte Lindqvist[a,2]

*[a]Department of Biological Sciences, University at Buffalo, Buffalo, NY 14260; [b]Daicel Arbor Biosciences, 5840 Interface Dr. Suite 101, Ann Arbor, MI 48103; [c]Department of Mathematical Sciences, University of Oulu, 90014 Oulu, Finland; [d]Far Northwestern Institute of Art and Science, Alaska Office, Studio B, 429 D Street, Anchorage, AK 99501; [e]Alaska Department of Fish and Game, Anchorage, AK 99518; [f]Alaska Department of Fish and Game, Fairbanks, AK 99701; [g]Natural History Museum, University of Oslo, 0318 Oslo, Norway; [h]School of Biological Sciences, Nanyang Technological University, Singapore 637551; [i]Organismal and Evolutionary Biology Research Programme, University of Helsinki, 00014 Helsinki, Finland; [j]Bioinformatics Research Centre, Aarhus University, 8000 Aarhus, Denmark; [k]Singapore Centre for Environmental Life Sciences Engineering, Nanyang Technological University, Singapore 637551; [l]Laboratorio Nacional de Genómica para la Biodiversidad/Unidad de Genómica Avanzada, Centro de Investigación y Estudios Avanzados del Instituto Politecnico Nacional, 36500 Irapuato, Guanajuato, México; [m]Institute of Genomics for Crop Abiotic Stress Tolerance, Department of Plant and Soil Science, Texas Tech University, Lubbock, TX 79430*

[1]T.L. and K.L. contributed equally to this work.

**This file includes:**

Supplementary materials and methods, Text S1 to S21
Figures S1 to S25
SI References

**Other supplementary materials for this manuscript include the following:**

External documents SI-FigureS14.pdf, SI-FigureS15.pdf and SI-FigureS16.pdf.
Supplementary tables in SI-Appendix_TablesS1-7.xlxs.

### Supplementary Materials and Methods

#### S1. Geological settings and dating of the ancient Poolepynten polar bear mandible

The ancient polar bear mandible is very well preserved. The only tooth remaining is the canine, which is pre-mortem worn at the apex, suggesting that it belonged to an adult bear (1, 2). As previously reported, the polar bear mandible was excavated in-situ at the Poolepynten coastal cliffs, Prins Karls Forland, western Svalbard (1). The lithostratigraphy, depositional history, and environmental development of these coastal cliffs have been studied extensively by geologists (1, 3-5). The polar bear mandible was discovered in the lowest unit A (4) that consists of three units A1–A3, representing two glacio-isostatically controlled regression cycles and implying two separate deglaciation sequences (3). An Eemian (c. 130–115 ka) interglacial origin for the polar bear mandible that was excavated from the lowermost unit A1 is supported by multiple lines of evidence (3, 4):

1. Unit A1, similarly to A3, is a thin bed of massive to laminated fine sand with scattered pebbles, shell fragments and occasionally occurring subfossil kelp on bed surfaces, containing an abundant and diversified foraminiferal fauna, dominated by Arctic species similar to modern fauna in shallow sites near Svalbard, clearly suggesting it is marine, deposited in a shallow marine to sublittoral setting.
2. $^{14}$C radiocarbon dating of seaweed from unit A yielded an infinite age, while Infrared Stimulated Luminescence (IRSL) dating of a sediment sample suggested an age between 150–80 ka.
3. Optically Stimulated Luminescence (OSL) dating of two sediment (laminated sand) samples from A1 returns dates of 118 ± 13 and 105 ± 10 ka, respectively, which are on the young side of the Eemian, but underestimation of quartz OSL ages from Eemian deposits has been described from other sites, and the age underestimation may be of the order of 10% (3).
4. Shells from unit A1 have an average amino acid ratio of 0.045 ± 0.001, while shells from unit A3 (the upper part of unit A) have significantly lower ratios of 0.025 ± 0.001 (4), which suggests that the two sampling points are separated by a sufficiently long and relatively warm time interval to allow for this amount of racemization. Such a pattern could be explained if the lower part of unit A was Eemian in age, and the upper part was Early Weichselian, which is consistent with the OSL ages.
5. A total of nine OSL ages from Poolepynten (ranging from ~1000 to ~120 000 years) correspond well with within the described stratigraphic units E–A with successively older ages for lower units, and largely match previous chronological data ($^{14}$C, IRSL) of the site, demonstrating relative dating by lithostratigraphy.
6. The units A1, A3, C and D2 all represent marine or littoral sedimentation during high relative sea-level events, which were preceded by regional glaciations, are recorded in the Poolepynten succession. They date to the Eemian, the Early Weichselian, the Middle Weichselian and the Late Weichselian/early Holocene, respectively.

#### S2. Sampling of modern Alaskan bears

For the new bear genome sequencing, and for microsatellite analyses associated with kinship analyses, we obtained tissue samples from wild populations collected by and held at the Alaska Science Center (USGS) and Alaska Department of Fish and Game. Blood and tissue samples were collected following standard procedures (6, 7). Briefly, blood samples were stored in

EDTA, whole blood Vacutainer® tubes, or a blood preservation buffer(8). All tissue and skin biopsies were placed in a preservation buffer (4 M Urea, 0.2 M NaCl, 100 mM Tris HCL pH 8.0, 0.5% n-lauroyl-sarcosine, 10 mM EDTA).

The samples selected for the new bear genome sequencing (Table S1) comprised two polar bears from the Chukchi Sea (AK017) and Southern Beaufort Sea (AK034), one individual (CON001) from Yellowstone National Park and seven individuals from throughout Alaska (Fig. S1A). Three of these Alaskan brown bear samples were preliminarily confirmed as belonging to the western Alaska (Seward Peninsula BB020 and Douglas River WB039) and the eastern Alaska (Anchorage EB027) maternal lineages. The remaining four samples were from the North Slope of Alaska, where polar and brown bears have recently been observed to come into contact, possibly attracted by subsistence-harvested bowhead whale (*Balaena mysticetus*) remains during the fall open-water period (9): a light-colored brown bear male (BB034; Fig. S1B) that is thought to have fathered several light-colored cubs in the region (D. Shideler, unpubl.), a brown bear female (BB049) and her light-colored cub (BB059), and one individual from Kaktovik (BB037). We wanted to test if light pelage might be inherited through a relationship with the light-colored male (BB034) and performed a microsatellite-based kinship analysis that tests the probability that two alleles sampled from each individual of a pair are identical by descent, as described below. This analysis confirmed the parent-child relationship between the light-colored cub (BB059) and its brown mother (BB049). It also determined that the light-colored adult male bear (BB034) is not a close relative of this cub. It is worth noting that both of the light-colored brown bears (BB034 and BB059) hold typical brown bear mitochondrial haplotypes (Fig. S5). The North Slope of Alaska encompasses bears with either eastern Beringian or western Beringian mitochondrial haplotypes (Fig. S5), suggesting that North Slope brown bears contain considerable matrilineal phylogeographic structure.

**S3. Kinship analysis**

We extracted DNA from blood or muscle ($n$ = 34) and hair ($n$ = 99) tissue samples from brown bears sampled between 1992 and 2013 from the Kaktovik area and other locales along the North Slope of Alaska—including the cub with light pelage (BB059), his putative mother (BB049) and father (BB034), as well as the putatively unrelated BB037—using a salt extraction procedure described in (10) and modified as described in (11) for blood and muscle samples, and, for hairs, procedures reported in (12), developed specifically for low-quality and low-quantity tissue sources. Detailed sample information is included in (13). We quantified total DNA using fluorometry and diluted all DNA to 50 ng $\mu L^{-1}$ working concentrations for subsequent analyses.

We genotyped DNA extracted from all specimens at 22 microsatellite loci isolated in ursid and canid species described in (14) (loci G1A, G1D, G10B, G10L, G10O), (15) (loci G10C, G10M, G10P, G10X), (16) (loci G10H, G10J, G10U), (17) (loci UarMU23, UarMU26, UarMU50, UarMU51, and UarMU59), (18) (CXX20, CXX110), (19) (locus MSUT-2), (20) (locus CPH9) and REN145P07 (GenBank Accession AJ411284, F: TGGAAAGGTTTGCACTCTGA; R: AGCCTCCCCATTTCACAGAT). PCR amplifications were carried out in 8 multiplex reactions, each in a final volume of 10 mL and contained 2–100 ng genomic DNA, 0.2 mM dNTPs, [3.6–4.0 *p*moles unlabeled primers, 0.06–0.4 *p*moles IRD-labeled primer], 1.0 mg BSA, 1X PCR buffer (Perkin Elmer Cetus I), and 0.3 units Amplitaq DNA polymerase (PE Biosystems, Forest City, CA). PCR reactions began with 94°C for 2 min and continued with 40 cycles each of 94°C for 15 sec; 50°C for 15 sec; 72°C for 30 sec. A 30 min extension at 72°C concluded each reaction.

The sex of the bear that left each hair sample was determined using PCR amplification of the *amelogenin* (AMEL) X and Y gametologs, using the following primers: CST1836: 5'-TCCCAGCCCCAGCCCGTCCAG-3'; CST1837R: GCTTCCAGAGGCAGGTCAGGA-3'). The primer CST1837 was synthesized with a universal M13R tail (GATTTAGGTGACACTATAG) to facilitate universal tailed PCR reactions following (21). All molecular sexing PCR reactions were performed on a Bio-Rad (Hercules, California) thermalcycler. PCR amplifications were carried and using procedures and parameters identical to those described for species determination. PCR products were electrophoresed on a 48-well 18-cm 6% polyacrylamide gel on a LI-COR 4200LR automated sequencer (LI-COR, Inc., Lincoln, Nebraska). The sample was interpreted to derive from a female if a single band (AMELX/AMELX, 150/150 bp, including the universal tail sequences) was amplified and a male if two bands (AMELX/AMELY, 150/196-205 bp, including the universal tail sequences) were amplified.

Because both brown and polar bears would be expected to leave hair samples around Kaktovik, where both species visit bowhead whale bone sites, we used sequence data from the mitochondrial DNA control region that is diagnostic of each species to verify the hairs derived from brown bears and not polar bears. Primers, amplification and sequencing methods followed those reported in (22, 23).

We electrophoresed fluorescently labeled PCR products on a 48-well 6% polyacrylamide gel on a LI-COR 4200 LR or $IR^2$ DNA automated sequencer (LI-COR, Lincoln NE). For allele size standardization we included 2-4 samples as standards of known size (22, 24), initially sized against a fluorescently-labeled M13 sequence ladder of known size. These samples were used in each subsequent gel as size standards, occupying at least six lanes across each gel. Based on these standards, genotypes for each individual were determined using GeneImagIR 4.05 software (Scanalytics, Inc.). For genotype quality control, >18% of the samples were extracted, amplified, and genotyped in duplicate; in some instances, some individuals were known to be represented by multiple specimens. In addition, data from 14 loci collected from BB034 in an independent laboratory as part of allied research (unpublished data) was examined and found to correspond precisely with genotype data collected by us. Overall, based on quality control reprocessing as well as duplicate and independent analyses, locus error rate was < 0.0001%. MICROCHECKER 2.2.3 (25) was used to identify genotyping errors and check for null alleles. We used sterile technique in the handling of all DNA, and all PCR procedures were done with positive and negative controls to verify amplification without contamination.

*Genetic diversity and marker resolution*

We used the Microsatellite Toolkit Version 2.1 (26) to identify matching samples among genotyped individuals to generate a dataset comprised only of unique individuals. There were seven unique genotypes among the 99 hair samples collected from brown bears sampled at Kaktovik; all were determined to be males. To estimate levels of genetic diversity metrics representing brown bears sampled from the Kaktovik/North Slope region, we removed the data associated with the ivory-colored cub (BB059) and putative first order (parent-offspring) relatives (BB049, BB034), as well as other hypothesized first-order relatives estimated from field observations. We also removed any sample with <60% of the microsatellite data, yielding a dataset comprised of 28 individuals representing the Kaktovik area and elsewhere on the North Slope of Alaska. This dataset was then augmented with genotype data collected from 14 of the 22 loci from 3 bears sampled from locales on the North Slope of Alaska as part of independent collection efforts (SF), resulting in a final baseline dataset for estimation of genetic diversity as part of this study. Details of the samples are given in (13).

Mean number of alleles per locus was 6.91, observed heterozygosity ($H_O$) was 0.75, and expected heterozygosity ($H_E$) was 0.75. All microsatellite loci were tested for gametic phase disequilibrium and for deviations from Hardy-Weinberg equilibrium (HWE) using the Fisher's Exact Test in GENEPOP'007 (27, 28). The Arctic coastal plain population showed no deviation from HWE overall ($\chi 2 = 40.05$, df = 44; P = 0.64) or at any locus (P >0.08). No locus showed evidence of null alleles, allelic dropout, or scoring error, based on analysis using MICROCHECKER. There were 16 locus-by-locus pairwise comparisons of significant linkage disequilibrium among 231 possible pairwise comparisons, higher than the number (11) expected to occur randomly. Probabilities of identity (PID; the probability that two identical genotypes from two different individuals could be obtained by chance, (29); PIDsib, (30)) in a randomly-breeding population were obtained using GIMLET and were very low ($PID_{biased} = 1.24x10^{-22}$; $PID_{unbiased} = 1.11x10^{-24}$; PIDsib = $2.65x10^{-9}$). Overall combined probability of parental exclusion (Q)—calculated using locus-specific Q values obtained using PowerStats Version 1.3.2 (31) and employing methods described in Jamieson and Taylor (1997, eq. 4)—were maximal (Q = 1.0) and exceeded the theoretical threshold of 0.99% recommended for when excluding incorrect parentage (32). Following analyses of baseline representative data, the dataset was augmented with with genotypes from the putative first- and second-order relatives, including two sets of ivory-colored brown bear cubs/yearlings, and subjected to analyses of relatedness and relationship. Microsatellite data from all samples assayed, including the 31 individuals used to generate genetic diversity information, the two ivory-colored yearlings and putative mother and all hypothesized first- and second-order relatives, are provided in (13).

*Relatedness and family relationship analyses*
We used ML-RELATE (33) to determine average relatedness of the ivory-colored cub (BB059) to their putative mother (BB034) and putative father (BB034). We also check relatedness of all three of these individuals to BB037. Relatedness was estimated between all pairs using the maximum likelihood estimator (reviewed in 34) of relatedness (*r*), to estimate common pedigree relationships (unrelated, U; half-siblings, HS; full siblings, FS, and parent-offspring, PO) between individuals, and for estimated PO dyads, evaluate the possibility that the dyad represented a HS rather PO dyad. Results of the kinship analysis estimated a first-order relationship (PO, LnL(R) = -97.21, *r* = 0.5926) between cub BB059 and the putative mother (BB049), and the likelihood that the first-order relationship was PO rather than FS was significant (P = 002, based on 500 simulations). However, analyses failed to find a first- or second-order relationship between cub BB059 and the putative father (BB034); the two individuals were unrelated (U, LnL(R) = -103.17, *r* = 0.0000), with the likelihood that the dyad was unrelated was significantly more likely (P <0.001) than in a first- (PO: P = 0.964, FS: P = 0.886) or even second-order (HS: P = 0.596). Similarly, but as expected, we failed to find a close

familial relationship between BB037 and the cub BB059 (U, LnL(R) =-113.22, $r = 0.00$), mother BB049 (U, LnL(R) =-110.52, $r = 0.00$) or BB034 (U, LnL(R) =-101.55, r = 0.00).

**S4. DNA extraction and genome sequencing**

For the ancient Poolepynten polar bear specimen, powder was obtained from the canine by using a dental drill equipped with low-temp drill bits and split into 50–100 mg aliquots. Genomic DNA was extracted following the column-based method described in (35). All extractions were carried out in a cleanroom dedicated to ancient DNA work, physically separate from any processing, including nucleic acid extraction and amplification, of modern samples. Negative controls were prepared alongside each batch of DNA extraction.

To maximize its genomic coverage, multiple libraries were generated from DNA extracted from the ancient Poolepynten polar bear and sequenced using different methods. Table S2 describes sequencing and mapping statistics for the individual and combined libraries from the ancient Poolepynten polar bear.

First, we used data that were previously published and had been generated using the Illumina HiSeq 2000 platform (2). Of the total 164 Gb, we restricted our sampling to the following NCBI SRA accessions: SRX156143–156150 (representing 113 Gb). This subsample was selected due to concerns over three particular Illumina HiSeq lanes (accessions SRX156151–156153) that were processed from DNA extracts that potentially carried low-grade contamination. Hence, to be safe, we here only included sequence data available from SRX156143–156150, and we denote this subsample APBv2 (Table S2).

In addition, new sequencing libraries were prepared and processed using shotgun sequencing and whole genome enrichment on the Ion Torrent and Illumina sequencing platforms, as described below.

The ancient polar bear DNA extracts (EL03, EL04, EL05, EL06) were processed by Daicel Arbor Biosciences (http://www.arborbiosci.com) for preparation of Ion Torrent sequencing libraries, whole-genome enrichment (WGE) and sequencing. Biotinylated RNA baits for WGE were synthesized at Daicel Arbor Biosciences from DNA extracted from three modern samples representing polar, brown, and American black bear, where ~1.3 ng of genomic DNA from each species was pooled. The Ion Torrent sequencing libraries were constructed with the NEBNext Fast DNA library preparation kit (New England Biolabs, E6270L) following the manufacturer's protocol with modifications applied as follows: doubling of time in end repair and ligation step, using 2.5X SPRI beads for purification, and Kapa HiFi Uracil+ PCR enzyme (Roche) for amplification. Each library was then individually target-enriched using 500 ng of the WGE RNA baits. The standard myBaits v. 3.0 protocol was applied with hybridization for 49 h at 65 °C. The enriched libraries were quantified with spectrofluorometry, which indicated between 63.7 and 219.3 ng dsDNA per library. Three non-enriched libraries (EL03, EL04, EL05) and all four enriched libraries were sequenced on the Ion Proton platform using the Ion PI Chip Kit v2 chemistry. Following sequencing, reads were de-multiplexed, quality trimmed and filtered using the default settings on the Ion Torrent Suite v. 4.4.3. WGE and shotgun sequencing reads were then combined for downstream data processing (denoted "WGE" in Table S2).

Illumina Truseq dual-barcoded shotgun libraries were prepared from three DNA extracts (EL09, EL10, and EL11) at Daicel Arbor Biosciences without sonication. Libraries were constructed using the blunt-end ligation module from NEBNext Fast DNA library preparation kit (New England Biolabs, E6270L) with an extended double-time treatment and blunt-end adapters synthesized by Daicel Arbor Biosciences. The Kapa HiFi Uracil+ PCR enzyme (Roche) was used for PCR amplification. Spectrofluorometry quantification of sample libraries indicated between 167 and 977 ng of DNA. The library from EL11 was then target enriched using the Daicel Arbor Biosciences WGE RNA baits described above and a total of 60 ng DNA was obtained from the enrichment. Both the two shotgun libraries (EL09 and EL10) and the WGE library (EL11) were paired-end sequenced on the Illumina HiSeqX platform at SCELSE, NTU, Singapore.

Two DNA extracts (EL14 and EL15) were processed and sequenced at SCELSE-NTU, Singapore. The libraries were prepared using the Accel-NGS 2S Plus DNA Kit (Swift Biosciences), a method previously employed for metagenomic sequencing of low-biomass air samples (36). For library preparation, a total DNA input amount of 2.2ng and 2.7ng were used for EL14 and EL15, respectively, and both libraries were amplified with 12 PCR cycles. In addition, DNA damage repair was performed on EL15 prior to library preparation, using the PreCR Repair Mix (New England BioLabs), following the manufacturer's instructions (sequential reaction protocol). The shotgun libraries were then sequenced in two lanes on the Illumina HiSeqX platform at a read-length of 150bp paired-end.

For the modern bears, DNA was extracted from tissue and blood samples following either the protocols listed in Medrano et al. (10), with the modifications reported in Sonsthagen et al. (11), or using the DNeasy Blood and Tissue Kit (QIAGEN, USA) according to the manufacturer's specifications. DNA libraries were prepared and sequenced at SCELSE, NTU, using the Illumina TruSeq Nano DNA Library Kit. The DNA samples were sheared on a Covaris E220 to ~450bp, following the manufacturer's recommendation, and tagged with one of Illumina's TruSeq LT DNA barcodes to enable sample pooling for sequencing. Finished libraries were quantitated using Invitrogen's Picogreen assay and the average library size was determined on Bioanalyzer 2100, using a DNA 7500 chip (Agilent). Library concentrations were then normalized to 4nM and validated by qPCR on a ViiA-7 real-time thermocycler (Applied Biosystems), using the Kapa library quantification kit for Illumina platforms (Kapa Biosystems). The libraries were then pooled at equimolar concentrations and sequenced in four flowcells (32 lanes) on the Illumina HiSeqX platform at a read-length of 150bp paired-end. See Table S3 for a summary of the sequencing and mapping statistics for the new modern bear genomes.

**S5. Sequence data processing and mapping**

We processed the raw sequencing data from each ancient polar bear library separately. First, we removed adapter sequences and trimmed low-quality bases using Trimmomatic as described below, except we discarded reads that were less than 20 bp, then trimmed any T in the first one or two bases and any A in the last one or two bases ("end trimming") from the reads to reduce the effects of cytosine deamination in ancient DNA. Next, for Illumina sequencing reads, we merged the paired-end Illumina sequencing reads with ≥ 11 nt overlap into a single sequence using FLASH (37), then aligned the merged reads and the unmerged reads to the reference using BWA (38) mem single-end mode and paired-end mode, respectively. For Ion Torrent single-end reads, we directly aligned the reads to the reference using BWA mem single-end mode after trimming the ends. PCR duplicated reads were removed using the MarkDuplicates tool in the Picard software suite v. 2.7.1 (http://broadinstitute.github.io/picard/) applying the "LENIENT" validation stringency. The generated BAM files from all libraries were merged using SAMtools v. 1.3 (39). See Table S2 for a summary of the mapping statistics of each APB library.

To remove adapter sequences and trim low-quality bases from the raw modern bear sequence reads, we used Trimmomatic v. 0.38 (40) with the following settings:

1. remove leading and trailing low-quality bases (quality score ≤ 3) and N bases,
2. remove low-quality bases (quality score ≤ 10) within a 4-base wide sliding window, and
3. discard reads that are less than 30 bases long after trimming low-quality bases.

We then aligned the trimmed reads to the chromosome-length HiC-assembly (41, 42) based on the polar bear draft genome assembly UrsMar_1.0 (GCF_000687225.1) (43) using BWA mem v. 0.7.17 with default parameters for paired-end reads mapping, and generated the BAM files using SAMtools with option "-F 4 -q 30" to only output reads that are mapped in proper pairs and have mapping quality over 30. We then performed PCR duplicates removal as described above. Individual genome coverage was calculated using the genomeCoverageBed tool in the BEDtools software suite v. 2.23.0 (44). See Table S3 for a summary of the mapping statistics of the modern bears.

To assemble the mitochondrial genomes from the new brown bear data, we used a brown bear mitochondrial genome as a reference (GenBank accession AF303110.1) and performed mapping of the trimmed reads and PCR duplicate removal as described above. A consensus sequence was called using the "mpileup" command in SAMtools in haploid mode.

**S6. Postmortem DNA damage profiling and authentication of the ancient polar bear data**

Postmortem, DNA is rapidly degraded by exogenous microorganisms that feed on and degrade macromolecules, as well as endogenous nucleases and hydrolytic and oxidative processes, resulting in DNA fragmentation, base modifications, and a reduction of overall DNA amount (45). Although this post-mortem DNA damage challenges attempts to recover and sequence ancient DNA, characterization and quantification of damage patterns can serve as authentication of ancient DNA. We computed post-mortem DNA damage profiles for each sequencing library of the ancient polar bear using mapDamage2 (46) and PMDtools (47). As expected, we observed elevated levels of deaminated nucleotide residues accumulated toward the ends of the molecules for all the libraries (Fig. S2). We also generated DNA damage profiles with mapDamage2 of libraries where ends of sequence reads had been trimmed (see above) and compared the damage patterns between pre-end trimming and post-end trimming dataset. The C to T and G to A substitution frequencies were substantially reduced after end trimming (not shown), confirming that end trimming successfully had removed deaminated cytosine residues.

We also assembled the mitogenomes from each library constructed for APB (Table S2) using the pipeline described above and reconstructed a phylogenetic tree adding these mitogenomes to the alignment of complete mitogenomes presented here (Fig. S5). The results (Fig. S3) clearly shows that all individual APB libraries acquired in this study grouped with the previously published mitogenome from the same ancient polar bear specimen (48) (GenBank accession GU573488) in a position sister to all modern polar bears, indicating the endogenous nature of all new APB libraries.

#### S7. SNP calling and filtering

We divided the scaffolds of the reference genome into 204 partitions to speed up the process of nuclear SNP calling and performed SNP calling across all bear samples in parallel for each partition using default settings in HaplotypeCaller, followed by GenomicsDBImport and GenotypeGVCFs in GATK toolkit v. 4.1.1.0 (49). We then concatenated the output VCF files from all partitions into one final VCF file using gatherVCFs in GATK.

We first excluded insertions/deletions (indels) sites using SelectVariants in GATK, and following the GATK Best Practices recommendations (50, 51), we applied the standard hard filtering for Fisher strand bias (FS > 60.0), mapping quality (MQ < 40.0), variant confidence normalized by depth (QD < 2.0), mapping quality rank sum test (MQRankSum < -12.5), and site position within reads rank sum test (ReadPosRankSum < -8.0). Next, to identify sex chromosomes in the reference, we respectively aligned the 12 X chromosome scaffolds and 5 Y chromosome scaffolds identified from a previously published polar bear genome (43) to the reference using nucmer from MUMmer 3.0 (52). As a result, scaffold 1 was identified as the X chromosome, while a few scaffolds (scaffold297, scaffold318, scaffold369, scaffold579 and scaffold605) that are less than 1 Mb were identified containing Y-chromosome sequences. We then extracted autosomal scaffolds and X chromosomal scaffold from the VCF file respectively, and applied the additional filtering using VCFtools v. 0.1.13 (53): (1) extract scaffolds ≥ 1 Mb and (2) include sites with a depth of coverage 5–500 with option "--minDP 5" and "--maxDP 500", resulting in our dataset DS1 (Table S5).

Additionally, for some analyses (DS2–3 and DS6–8; Table S5) we also removed singletons, using option minor allele count greater than or equal to 2 "--mac 2", because they provide extremely low statistical power (54-56). Singletons are SNPs that have only one copy of the minor allele among all samples, including rare mutations, sequencing errors, and DNA damage residues in the ancient sample. For datasets DS3 and DS8 we excluded individuals that have less than 8X depth of coverage and excluded SNPs that contain any missing data ("--max-missing 1"). To explore if increased deamination in the ancient polar bear sample had any effect on the results from certain analyses, we also generated datasets that only contained transversion sites (DS5, DS6, DS8).

For further analyses, PED and MAP files were generated from the filtered VCF files using the option "--plink" in VCFtools. The PED and MAP files were converted into GENO, SNP and INDV files using CONVERTF in the ADMIXTOOLS software suite (57).

#### S8. Phylogenetic analyses

We performed phylogenetic analyses based on complete mitochondrial genomes (Table S4) and autosomal SNPs (Fig. 1, Figs. S4–S5). We used SNPhylo v. 20160204 (58) on the autosomal SNP DS7 to reduce SNP redundancy and generated a FASTA input for tree reconstruction. All SNPhylo parameters were set to default except the LD threshold option "-l" which was set to 0.5. Maximum likelihood phylogenetic trees were reconstructed using RAxML-HPC on XSEDE v. 8.2.12 in CIPRES (59).

To provide a first visualization of discordance among the SNP data that might stem from past admixture among bear species and interspecific populations, we also employed the Neighbor-Net method, a distance-based method based on neighbor-joining that generates phylogenetic networks (60). To generate this distance-based phylogenetic network using Logdet distances we applied the same SNP data used for phylogenetic tree reconstruction. Character incongruence that is manifest as extra edges in such networks (beyond a perfectly bifurcating tree) have been variously interpreted by other investigators to reflect admixture and/or incomplete lineage sorting phenomena (61, 62).

**S9. Principal component analysis**

To infer population structure among all bear samples with a coverage of depth > 8X (DS3; Table S5), we carried out principal component analysis (PCA) using the smartpca program distributed with the Eigensoft package v. 6.0.1 (63, 64). Default settings were used except outlier removal was turned off. PCA analyses were also carried out with the dataset that excludes nucleotide transitions (DS8), in addition to an analysis that excluded close relatives suggested by the $f_3$-statistic (see below). These analyses gave identical results and only results including all DS3 individuals are shown here. We first conducted a PCA analysis on all bears, including both APB and modern bears (Fig. 1C, S6A). The clustering of the two bear species along eigenvector 1 was predominant in this analysis, in addition to four major clusters among brown bears: European brown bears (EBB), Alaskan mainland brown bears (BB), a cluster of the ABC brown bears from Admiralty Island (ABC-A) and continental brown bears (YB), and a cluster of ABC brown bears from Baranof and Chichagof Islands (ABC-BC). To explore more detailed structure and relationships among the brown bear populations, we then conducted a PCA analysis only on brown bear samples and polar bears, respectively. The brown bear individuals grouped into five major population clusters: including EBB, BB, YB, ABC-A, and ABC-BC (Fig. S6B). The population structure among polar bears exhibited strong differentiation between the ancient and the modern polar bears (Fig. S6C), while the clustering of modern polar bears is associated with their geographic locality (Fig. S6D): WG (West Greenland), EG (East Greenland), AK (Alaska), and SV (Svalbard), with AK and EG polar bears clustering close together.

**S10. Genomic diversity**

We estimated the autosomal heterozygosity from genotype calls with "--het" in VCFtools. The individual observed heterozygosity was calculated as "(N_SITE - O(HOM)) /N_SITE" from the output. To assess the effect of filtering on measures of heterozygosity, we explored multiple datasets with different filters (see Table S5): dataset DS1, DS2 (excludes private alleles), DS3 (includes only individuals with depth of coverage ≥ 8X and no missing data allowed), DS5 (transversion sites only of DS1), DS6 (transversion sites only of DS2), and DS8 (transversion sites only of DS3). Bar plots of the heterozygosity frequencies among all polar bear and brown bear genomes (Fig. S7A,B) and box plots of heterozygosity frequencies across polar and brown bear populations (Fig. 2A, S8) identified from PCA analysis demonstrate overall dramatically lower heterozygosity levels among polar bears compared to brown bears. Interestingly, the ancient polar bear displays an increased level of heterozygosity compared to modern polar bears. In addition, the continental brown bears display the lowest heterozygosity among the brown bear populations. While the high number of private alleles (here, alleles unique to the ancient polar bear compared to modern polar bears; Fig. S7C) caused a dramatic increase in heterozygosity for the ancient polar bear, heterozygosity levels were still higher in the ancient polar bear when considering all other autosomal SNPs (Fig. S8).

We measured nucleotide diversity within each polar and brown bear population identified from PCA analysis in a window size of 10 kb using VCFtools with option "--window-pi 10000". The results (Fig. 2B) show that brown bear populations have overall higher levels of nucleotide diversity than polar bear populations, and that among polar bear populations, APB exhibits a higher level of nucleotide diversity than modern polar bears. This is consistent with the patterns observed in heterozygosity across polar and brown bear populations (Fig. 2A, S7, S8).

We also calculated the pairwise fixation index $F_{ST}$ between populations to measure population differentiation using the VCFtools option "--weir-fst-pop" and "--fst-window-size 10000". The populations were identified from clusters in the PCA analysis. As expected, the results (Fig. S9) demonstrate low population differentiation among different modern polar bear populations (mean weighted $F_{ST}$ = 0.031–0.054), compared to among different brown bear populations (mean weighted $F_{ST}$ = 0.127–0.276). Interestingly, the differentiation between APB and the modern polar bears ($F_{ST}$ = 0.245–0.266) is within the range of brown bear population differentiation. Similarly, the differentiation between APB and brown bears is generally lower ($F_{ST}$ = 0.407–0.591) than between the modern polar bear populations and brown bear populations ($F_{ST}$ = 0.57–0.796).

**S11. Ancestral population size analysis using PSMC**

We used pairwise sequentially Markovian coalescent (PSMC) analysis (65) to infer demographic history for all bear populations. First, we used SAMtools to generate the diploid consensus sequence for each sample with option "-C 50", which will reduce the reads with multiple mismatches, and "-d minimum coverage -D maximum coverage", which only considers positions with coverage greater than one third of the average depth of coverage but less than twice. We also performed a total of 100 bootstrap replicates with the option "-b" in PSMC for APB (Fig. S10). We have observed previously that depth of coverage can affect PSMC results (2). For a diploid genome with low coverage, randomly lost heterozygotes due to lack of coverage on both alleles will lead to an underestimated population size and more recent timeline. This has the same effect as a smaller mutation rate and can be corrected by applying the false negative rate (FNR) in PSMC. Because shifts in time scaling could be caused by differences in generation times between the two species, we plotted the PSMC results from high-coverage genomes representing our defined populations using different generation time (g) values. Hence, we rescaled the axes using (i) an average generation time for polar bear (g = 11.5 years) and brown bear (g = 10 years) following comprehensive assessments of the generation length in the two species (66, 67) and (ii) an average generation time "-g 10" as previously reported (2), as well as a per generation mutation rate "-u 1e-08". We applied the FNR value empirically to generate the best overlap of the curves for related samples with different depth of coverage.

We also picked one bear individual from each population and downsampled bam files from modern samples to approximately 10X (the depth of coverage of APB) using the DownsampleSam tool of the Picard toolkit. We plotted modern samples with APB using the previously reported generation time of g = 10 for all bears (Fig. S10D), as well as g = 11.5 and g = 10 for polar bears and brown bears respectively (Fig. 2C). We also plotted the downsampled high-coverage genomes using different g value estimates based on aligning the oldest peak for the brown bear PSMC curves to the polar bear (Fig. S10E).

As expected, the demographic historic trends stay the same, although the curves are shifted to slightly older time scales at higher generation time estimates. Aligning the oldest peak for the two species implied generation times that are inconsistent with timings calculated from generation length estimates, suggesting that a shift of the polar bear demographic curves towards modern times may at least in part be caused by higher levels of inbreeding among polar bears. In coalescent modeling, highly heterozygous regions coalesce further back in time, and therefore this shift could have resulted from events that reduce genome-level heterozygosity, such as high levels of inbreeding, as has been demonstrated previously in plant systems (68).

**S12. Estimating population size histories and split times using SMC++**

SMC++ is a statistical tool that can jointly infer effective population size histories and split times in diverged populations (69). It combines the computational efficiency of the site frequency spectrum and the advantage of utilizing LD information in coalescent HMMs. It employs a novel spline regularization scheme that greatly reduces estimation error, and phased genomes are not required. Here, we performed SMC++ on datasets DS3 (all SNPs from genomes > 8X coverage, excluding private alleles) using the default parameters and a per generation mutation rate of "1e-8". In brief, the DS3 VCF file was first converted to the SMC++ input with "vcf2smc". Then, demographic history was inferred by using the command "cv". This command uses cross-validation to obtain sensible model parameters for use during estimation and is the recommended way to run SMC++ as of version 1.15. The split time between pairs of populations was inferred by using "estimate" and "split". The populations were identified from clusters identified in the PCA analysis, although only two modern polar bear populations were used here. The ancient polar bear was excluded from the split estimate, as SMC++ is not currently able to incorporate an age from an ancient sample (https://github.com/popgenmethods/smcpp/issues/114). It is also important to note that the current version of the implementation assumes a "clean split" model, in which no gene flow occurs after the populations split (69). Hence, if past gene flow has taken place, split times may be underestimated and instead reflect cessation of gene flow between the diverged population. Demographic inferences are shown in Fig. S11. The split times between populations are shown in Fig. S12 and summarized in Table S6.

**S13. Divergence time estimation using a molecular clock**

We followed the strategy applied in (70) to estimate the average coalescence time between polar bears and brown bears. Each of the populations BLK, EBB (as European brown bears are regarded as free of polar bear admixture that would otherwise bias the approximation) and APB is represented by random alleles of a single individual. We used the dataset DS2 and restricted our analysis to C/G and A/T polymorphisms, because they are least affected by DNA degradation (71). Denote by $N_X$ the number of SNPs where the allele from population X differs from the two others, for instance, on a single locus allele C on APB and G on BLK and EBB would increase $N_{APB}$ by one. Assuming mutations occur at similar rate per year on each lineage, we get $N_{EBB} = N_{APB} + A*\rho$, where A is the age of the ancient sample, 115,000–130,000 years. From the data $N_{APB} = 368,285$, $N_{EBB} = 402,509$ we solve the genetic divergence time between brown bears and polar bears $N_{EBB}/\rho$ as 1.3–1.6 ma. As a genetic divergence time, this is an upper bound for the population divergence time. We note this split estimate is consistent with the coalescence time estimated by PSMC analysis that implies the population divergence between brown bears and polar bears happened over one million years ago (Fig. S10D–F).

We note, however, that there might be some enrichment of mutations on the EBB lineage compared to polar bears: using PB instead of APB (so that A = 0) gives $N_{PB} = 397{,}643 < 401{,}031 = N_{EBB}$. Accounting for the enrichment could bring the genetic divergence time estimate a little closer, but the difference is miniscule. Another potential issue is the use of DS2 that excludes private alleles: if some of the many private alleles of APB (see Fig. S7C) were genuine instead of sequencing errors and DNA degradation, our divergence time estimate is not old enough.

**S14. ADMIXTURE analysis**

ADMIXTURE uses a maximum likelihood-based approach for estimating individual ancestries from multilocus SNP genotype datasets (72). To infer the genetic structure of bear populations, we employed ADMIXTURE to datasets DS3 and DS8 for different K values ranging from 2 to 7 in an unsupervised mode, where K is the number of hypothetical ancestral source populations (Fig. S13). The convergence criterion for each run was when the log-likelihood increased by less than $10^{-4}$ between iterations. We also employed cross-validation, where the most suitable K value is identified as judged by minimizing an estimated prediction error on a grid of K values (73). Cross-validation indicated K = 4 as the best-fit K value considering four hypothetical ancestral source populations (Fig. S13C), with one each for American black bear and polar bears, and two ancestral populations largely corresponding to ABC and continental (YB) brown bears on the one hand, and Alaskan mainland (BB) and European (EBB) brown bears on the other hand (Fig. S13A, S13B). While the results do not exhibit admixture between modern polar bears and brown bears, slight admixture in APB was observed at K = 1, 2, 3 for DS3 (Fig. S13A). The polar bears sorted into several ancestral groups at K = 5–7, suggesting significant population structure among popular bears.

**S15. The $f_3$-statistics**

We next analyzed population structure of the bears using ADMIXTOOLS and *f*-statistics (57), which uses correlations in allele sharing to measure drift between populations or individuals. As a first test for admixture, we used the $f_3$-statistic, $f_3$(C; A, B). When this statistic is significantly negative, it provides evidence that C is admixed (although a positive value does not necessarily indicate that C is not admixed). The statistics capture drift on overlapping paths from C to A and C to B, and if C is admixed, some of this drift can contribute a negative value to the statistics. This can happen either when A or B is an outgroup, while the other is closely related to one of the source populations of the admixture, or when A and B are related to different source populations. The closer A and B are to the source populations of the admixture, the more negative $f_3$ becomes, so we can search for the most likely source population for an admixed C by looking for the pair (A, B) that gives the most negative $f_3$.

It is important to note that when C and either A or B are from the same population and the other is an outgroup, we are also likely to sometimes see a negative $f_3$ due to recent family structure. To see this, consider an example where A, B and C are single individuals, and A and C come from the same population. To formalize the relationship between A and C, we denote the proportion of the genome where A and C are independent by $\pi_0$, the proportion where they have exactly one chromosome identical by descent (IBD) by $\pi_1$, and the proportion where they have both chromosomes IBD by $\pi_2$. When two chromosomes are not IBD, the alleles can still match by chance (identical by state, IBS), and we model the allelic states using a binomial distribution and the population level frequencies c (for A and C who come from the same population) and b (for B). When an estimator for an $f_3$-statistic is constructed from estimators $\hat{a}$, $\hat{b}$ and $\hat{c}$ for the allele frequencies, a correction is necessary as derived by Reich et al. 2009 (74), written in an equivalent form:

$$\hat{f}_3(C;\, A, B) = (\hat{c} - \hat{a})(\hat{c} - \hat{b}) - \frac{\hat{c}(1 - \hat{c})}{n_c - 1},$$

where $n_c$ is the sample size of C; in our example $n_c = 2$ for the two chromosomes. In the presence of enough loci with chromosomes IBD, this estimator is no longer unbiased and does not match the correct value $f_3(C; A, B) = 0$. We break the analysis of expected behavior of the estimator at a random locus into three cases depending on the number of chromosomes IBD between A and C.
Case 0) All alleles are independent, probability $\pi_0$. By the standard rules for expected value, $E((\hat{c} - \hat{a})(\hat{c} - \hat{b})) = E(\hat{c}^2) - E(\hat{c})E(\hat{a}) - E(\hat{c})E(\hat{b}) + E(\hat{a})E(\hat{b})$. The following table demonstrates the values $E(\hat{c}) = c$ and $E(\hat{c}^2) = 0.5c(c + 1)$:

| **Probability** | **Genotype of C** | **$\hat{c}$** | **$\hat{c}^2$** |
|---|---|---|---|
| $c^2$ | mm | 1 | 1 |
| $2c(1 - c)$ | mM | 0.5 | 0.25 |
| $(1 - c)^2$ | MM | 0 | 0 |
| | | $E(\hat{c}) = c$ | $E(\hat{c}^2) = 0.5c(1 + c)$ |

Similarly, $E(\hat{a}) = c$ and $E(\hat{b}) = b$, so we get $E((\hat{c} - \hat{a})(\hat{c} - \hat{b})) = 0.5c(1 - c)$.
Case 1) One of the two chromosomes are IBD, probability $\pi_1$. This time, $\hat{a}$ and $\hat{c}$ are no longer independent, and we only have $E((\hat{c} - \hat{a})(\hat{c} - \hat{b})) = E(\hat{c}^2) - E(\hat{c}\hat{a}) - E(\hat{c})E(\hat{b}) + E(\hat{a})E(\hat{b})$. The following table and some arithmetics demonstrate the value $E(\hat{c}\hat{a}) = 0.25c + 0.75c^2$:

| **Probability** | **Genotype of C** | **Genotype of A** | **$\hat{c}$** | **$\hat{a}$** | **$\hat{c}\hat{a}$** |
|---|---|---|---|---|---|
| $c^3$ | **m**m | **m**m | 1 | 1 | 1 |
| $c^2(1 - c)$ | **m**m | **m**M | 1 | 0.5 | 0.5 |
| $c^2(1 - c)$ | **m**M | **m**m | 0.5 | 1 | 0.5 |
| $c(1 - c)^2$ | **m**M | **m**M | 0.5 | 0.5 | 0.25 |
| $c^2(1 - c)$ | **M**m | **M**m | 0.5 | 0.5 | 0.25 |
| $c(1 - c)^2$ | **M**m | **M**M | 0.5 | 0 | 0 |
| $c(1 - c)^2$ | **M**M | **M**m | 0 | 0.5 | 0 |
| $(1 - c)^3$ | **M**M | **M**M | 0 | 0 | 0 |
| | | | | | $E(\hat{c}\hat{a}) = 0.25c + 0.75c^2$ |

We used the bold font weight to mark the genotype that was IBD. The rest of the terms are already familiar to us, so we get $E((\hat{c} - \hat{a})(\hat{c} - \hat{b})) = 0.25c(1 - c)$.
Case 2) Both chromosomes are IBD, probability $\pi_2$. Trivially $E((\hat{c} - \hat{a})(\hat{c} - \hat{b})) =0$.
In all three cases, the correction gives $E(-\hat{c}(1 - \hat{c})) = -(E(\hat{c}) - E(\hat{c}^2)) = -0.5c(1 - c)$.
Noting that $2c(1 - c)$ is the heterozygosity, we have now derived the formula:

$$E(\hat{f}_3(C;\, A, B)) = \frac{HET_c}{4}\left(\pi_0 + \tfrac{1}{2}\pi_1 - 1\right).$$

In the case of a mother and a child, such as BB049 and BB059, we have $\pi_0 = \pi_2 = 0$, $\pi_1 = 1$. Then the estimator of the $f_3$-statistic necessarily becomes negative without presence of any admixture at all, as is the case whenever $\pi_0 < 1$.

We calculated the $f_3$-statistics on both an individual and population level for DS2 and DS6, with populations following the clusters identified with PCA. All possible combinations of three individuals (or populations for population level) were tested using default parameters. The resulting $f_3$ *Z*-scores were then corrected for multiple testing by converting *Z*-scores into *p*-values and applying a Benjamini-Hochberg false discovery rate correction, and subsequently converting back into *Z*-scores for plotting (75). The corrected *Z*-scores were visualized as heatmaps using ggplot2 in R (76) for the population (Fig. S14) and individual (Fig. S15 for all substitutions and Fig. S16 for transversion substitutions only) level comparisons.

##### S16. The $f_4$-statistics

The $f_4$-statistic (closely related to the *D*-statistic) is a four-taxon test of admixture that has become an important tool for estimating gene flow in population genomics (57, 70, 77). In a phylogenetic tree (A, (B, (C, D))) without admixture we expect $f_4$(A, B; C, D) = 0, but admixture between populations B and D makes the $f_4$-statistic positive and admixture between B and C makes it negative.

We used DS2 (Table S5) and a python script available at https://github.com/KalleLeppala/f_stats to calculate $f_4$-statistics between all populations and selected individuals. Populations were defined as clusters identified by the principal component analysis, with the simplification that all modern polar bears from Svalbard and Greenland were combined as PB. The uncertainty of the statistics was measured using a block jackknife of 10 SNP windows for variances, ignoring the covariances.

##### S17. Searching for actual segments

The $f_2$-statistic measures the amount of drift separating two populations (57, 74).

We set out to search for actual introgressed DNA fragments by calculating $f_2$-statistics on 50 kb windows using the python library available at https://github.com/KalleLeppala/f_stats and the data set DS2 (see Table S5). The library did not account for the upwards deviation inflicted by using the same sample to estimate the allele frequency twice, as described by Reich et al. 2009 (74). Therefore, each distance is overestimated by the heterozygosity rates of the two populations, divided by sample size. Each population was represented by a single diploid genome. We defined the distance between ABC-BC and PB as

$$d(\text{ABC} - \text{BC}, \text{PB}) = \frac{f_2(\text{ABC}-\text{BC},\text{PB})}{f_2(\text{ABC}-\text{BC},\text{PB}) + f_2(\text{ABC}-\text{BC},\text{EBB})}$$

and between ABC-BC and APB as

$$d(\text{ABC} - \text{BC}, \text{APB}) = \frac{f_2(\text{ABC}-\text{BC},\text{APB})}{f_2(\text{ABC}-\text{BC},\text{APB}) + f_2(\text{ABC}-\text{BC},\text{EBB})}.$$

The population EBB was used for calibration, because the distances are bound between zero and one. We computed these two distances as well as their difference on 50 kb windows (Fig. S17). We highlighted unusual, potentially introgressed regions, where any of the three variables (two distances and their difference) differed from its sample mean by more than 2.437 times its sample standard deviation on at least five consecutive windows (Fig. S18).

It is important to note that these potentially introgressed segments are less than 1 Mb in length, most of them only 250 kb. This is consistent with Liu et al. (43), who state that the admixture event did not occur during the last hundreds of generations (thousands of years). We also point out that recent gene flow from the ABC bears into modern polar bears would leave fragments with small *d*(ABC-BC, PB) and big *d*(ABC-BC, APB), of which we found only one potential example (the longest highlighted segment in Fig. S18).

**S18. Analysis on $f_4$-ratio estimation**

The fraction $\hat{f}$ was developed to estimate the proportion of Neanderthal admixture in populations with European ancestry (70). The $f_4$-ratio estimator $\hat{f}$ corresponds to the proportion of DNA a hybrid population H inherits from an introgressing population I after diverging from their sister populations H' and I', assuming the phylogeny (O, ((H', H), (I, I'))), where O is an outgroup, H is a potential hybrid population, H' is its unadmixed sister population, I is a potential introgressing population and I' is its sister population not involved in the admixture (notation our own). Allowing gene flow I→H, the proportion of DNA inherited from the I lineage in H is α. The proportion α can be estimated by the fraction

$$\hat{f} = \frac{S(H,\ H',I',O)}{S(I,H',I',O)}\ ,$$

where S( ) is the numerator of the *D*-statistic, i.e., the number of observed BABA-events minus the number of observed ABBA-events. We stress that $\hat{f}$ only equals α when the assumed phylogeny is correct and I→H is the only type of gene flow happening.

Since the expectation of S(W, X, Y, Z) is just the $f_4$-statistic $f_4$(W, X; Y, Z) = E((w − x)(y − z)), where the expectation is taken over all SNPs and the lowercase letters are the minor allele frequencies of the corresponding populations, we have

$$\hat{f} = \frac{f_4(H,\ H';\ I',O)}{f_4(I,H';\ I',O)}\ .$$

We can thus use the interpretation of $f_4$-statistics as overlapping paths (57) to gain insight on the $f_4$-ratio estimator $\hat{f}$. For example, Fig. S19 shows how under the assumptions made above $\hat{f}$ really equals α.

Cahill et al. (78) calculate $f_4$-ratios for unidirectional gene flow from polar bears into brown bears and vice versa, which we shall call $\hat{f}_B$ and $\hat{f}_P$. Both statistics use black bear (BLK) as the outgroup O. For $\hat{f}_B$, H and H' are some potential hybrid brown bear (HBB) and a Swedish (presumably unadmixed) brown bear (SWE), and I and I' are two polar bear samples (PB1 and PB2). Conversely, for $\hat{f}_P$ the two brown bears are swapped with the two polar bears. For all choices of representatives, they then observe $\hat{f}_B \gg 0$ (7.2–8.3% when HBB is from Baranof or

Chichagof islands) and $\hat{f}_P \approx 0$. However, the conclusion of only recent gene flow into ABC bears is not the only possible explanation for the observation, as the fractions $\hat{f}_B$ and $\hat{f}_P$ represent admixture proportions under assumptions that are not necessarily met. As demonstrated in Fig. S20, ancient gene flow from polar bears into ABC bears (Case II) or ancient gene flow from ABC bears into polar bears (Case III) with suitable branch lengths explains the observations just as well (but no other type of unidirectional gene flow event would do that alone), as would several more complicated scenarios with multiple admixture edges present.

Our closest match among the populations with (78) is keeping BLK, using ABC-BC as HBB, EBB as SWE and, say, PB and AK as PB1 and PB2. With these choices, using the $f_4$-statistics calculated earlier (see section S16 above), the results replicate $\hat{f}_B$ = 8.5% using O = BLK, H = ABC-BC, H' = EBB, I = PB and I' = AK, and $\hat{f}_P$ = 0.0% using O = BLK, H = PB, H' = AK, I = ABC-BC and I' = EBB. We also have access to the ancient sample APB, which is, while also admixed, more suitable to play the role of an unadmixed sister population of PB than AK is. Indeed, with APB the fractions increase $\hat{f}_B$ = 8.9% and $\hat{f}_P$ = 2.4%. Using APB as PB2 therefore rules out the three scenarios in Fig. S20 to explain the data using a single gene flow type. In particular it shows:

1. Some type of admixture happened before PB–AK -split and that using AK (or any modern polar bear) as the unadmixed polar bear sister population is therefore problematic.
2. A single unidirectional gene flow event, including the current consensus scenario, is not enough to explain the data and $f_4$-ratio analysis is therefore inadequate.

##### S19. The graph fitting method applied selectively

The interpretation of $f_4$-statistics as weighted overlaps between paths on a graph, derived by modelling genetic drift of SNPs as independent Wiener processes of the minor allele frequency in a population, presents a way to compare how well various candidate graphs fit SNP data. This idea was implemented in the qpGraph (57) and admixturegraph (79) software. Using a python library available at https://github.com/KalleLeppala/f_stats, we computed $f_2$-statistics (which are a special case of $f_4$-statistics) between the populations within each 500 kb window of DS2 (see Table 5). The library did not account for the upwards deviation inflicted by using the same sample to estimate the allele frequency twice, as described by Reich et al. 2009 (74). Therefore, each distance is overestimated by the heterozygosity rates of the two populations, divided by sample size. The uncertainty of the statistics was measured using a block jackknife of 10 SNP windows for variances, ignoring the covariances.

To distinguish the direction of gene flow, we typically need to use more than four populations. With the aim of making an additional point about timing, we studied the six populations BLK, EBB, BB, ABC-BC, APB and PB, and the 105 ways to arrange them into a tree. We used admixturegraph (79) to fit each tree to the 4319 $f_2$-datasets on the 500 kb windows. The cost function value C reported by admixturegraph is for the edge lengths $\theta_{best}$ that fit best, and can be scaled to an actual likelihood by

$$p(\text{observed } f_2 \mid \text{graph}) = \frac{\exp(-0.5\,C)}{\sqrt{(2\,\pi)^k\,|\Sigma|}},$$

where k is the number of statistics used (here k = 6(6−1)/2 = 15) and |Σ| is the product of the variance estimators of those statistics (since we ignore the covariance). We further approximate

$$p(\text{observed } f_2 \mid \text{graph}) \propto p(\text{observed } f_2 \mid \text{graph}, \theta_{\text{best}})$$

essentially replacing integrating over the parameters with taking the maximum over the parameters, then scale, getting the likelihoods of the 105 trees given the observed $f_2$-statistic data within the 500 kb window. The results are depicted as mosaic plots in Fig. 4A and Fig. S21.

Interpreting Fig. 4A and Fig. S21 is somewhat convoluted because of the possibility of incomplete lineage sorting and other (e.g., infraspecific) admixture events, and because there is no reason to think graph fitting is equally sensitive to all violations of the species tree. Therefore, we cannot simply compare the average likelihoods of two arbitrary trees. Still, we can say something using symmetry arguments.

Most regions are most compatible with the correct species history A (average likelihood 0.513). However, due to incomplete lineage sorting and admixture, different genomic regions will fit well with different trees. Many regions fit best with trees B or C (average likelihoods 0.285 and 0.102, respectively), which give alternative orders within the brown bear clade. This could be partially attributed to ILS, but since tree B has much higher average likelihood than tree C, admixture of EBB and BB with one another or with a ghost population could also explain these patterns. These potential complications are shown with grey branches in Fig. 4B. Indeed, if W = BLK, X = EBB, Y = BB and Z = ABC-BC, then ABBA-events and BABA-events are not balanced, yielding $D$ = -0.063 ($Z$-score 22.9).

Trees D and E (Fig. S21) model the direction that drift within brown bear phylogeny, versus drift within polar bear phylogeny, contributes alleles to each respective species in a putative admixture event. In D, drift from within the polar bear clade contributes alleles to brown bear phylogeny only via entry into the ABC-BC edge. In contrast, in E, brown bear phylogenetic drift, only from ABC-BC, contributes alleles to polar bear at the edge subtending APB+PB. In short, trees D and E (Fig. 4B) represent gene flow from ancient polar bears (predating the APB-PB split) into ABC-BC brown bears, and the inverse, respectively.

Tree D, the cyan tree in main text Fig. 4B, has the fourth best in fit (average likelihood 0.057). As visualized, gene flow from the ancestors of all polar bears into ABC bears (but not the other way) should make some genomic regions locally look like the cyan tree D. Let us rule out alternative explanations for the high average likelihood. Its success cannot be explained with ILS alone, because by symmetry ILS should favor the tree L (average likelihood 0.001, pictured in Fig. 4B) equally often. Neither can it be explained with ILS together with admixture between EBB and BB, or between both of them and a brown bear ghost population, because when gene flow pulls the EBB and BB lineages together before meeting the ABC-BC population, again by symmetry, ILS should equally often favor the tree G (Fig. S21) but it has a lower average likelihood of 0.005. Gene flow from a ghost population ancestral to both brown bears and polar bears (such as cave bears, see (80)) into both EBB and BB remains a possible explanation for the cyan tree D. However, such ghost admixture would make the comparatively bad fits of trees F (Fig. S21, average likelihood 0.007) and N = (BLK, (BB, ((EBB, ABC-BC), (APB, PB)))) (average likelihood 0.001), corresponding to ghost admixture into only one of the populations EBB and BB, somewhat surprising.

The fifth best-fitting tree is E (average likelihood 0.013), the brown tree from Fig. 4B. Again, our preferred explanation for the high likelihood of the brown tree E is gene flow from ABC bears into the ancestors of all polar bears. By symmetry, ILS alone should not favor it over tree J (Fig. 4B, average likelihood 0.003). Admixture between EBB and BB, or between both of them and a brown bear ghost population, would in fact act against regions locally looking like the brown tree E. Gene flow from two separate ghost populations ancestral to both brown bears and polar bears into both EBB and BB is hardly a possible explanation. Since the trees F (Fig. S21, average likelihood 0.007) and N = (BLK, (BB, ((EBB, ABC-BC), (APB, PB)))) (average likelihood 0.001) fit poorly in comparison, the two separate hypothetical gene flow events would have to concern the same genomic regions, which is unlikely.

The final, and sixth, best-fit tree discussed here, tree F (Fig. S21, average likelihood 0.007), is ambiguous: it could indicate ancient gene flow in either direction.

We conclude that due to lack of other plausible explanations for the occasional good fit of the cyan tree D and the brown tree E from Fig. 4B, there must have been substantial gene flow between ancestors of all polar bears and ancestors of ABC bears in both directions. Because all the well-fitting trees include the pair (APB, PB), while recent gene flow would result in regions breaking the bond, there is no evidence of modern gene flow occurring after the PB–APB -split.

Note, however, that because of asymmetry, the fit of the cyan tree D (0.057) and the brown tree E (0.013) should not be compared to one another:

1. The amount of gene flow sufficient to make the cyan tree D thrive need not be equal to the amount of gene flow sufficient to make the brown tree E thrive.
2. The ILS within the brown bear clade is still happening on the brown tree E, weakening the signal, but does nothing for the cyan tree D.
3. The tendency of EBB and BB to coalesce more than expected (quantified by $D = -0.063$) again does nothing for the cyan tree D, while making the brown tree E less likely.

This method of assessing the direction of gene flow resembles the idea used to determine the direction of the gene flow between brown bears and cave bears (80). However, it should be noted that gene flow from brown bears into the ancestors of both cave bears (opposite to the concluded direction) also contributes to Class 1 in their Figure 3.

#### S20. The graph fitting method applied with brute force

In a manner close to exhaustive, we searched for admixture graphs that fit well with the full set of $f_2$- or $f_4$-statistics ($f_4$-statistics calculated in section S16, $f_2$-statistics calculated similarly) including the last three populations YB, ABC-A and AK (abandoning the backbone tree). As the number of possible admixture graphs grows very fast with the number of leaves and admixture events, a full brute force search is impractical. Instead, we chose a four-stage approach of repeated brute force searches and expansions of a well-fitting subset from a previous stage into more complex graphs:

1. We fitted all the 10,395 8-leaf trees without using the population AK.

2. We chose a number of best fitting 8-leaf trees, for each added a new leaf for AK anywhere, and fitted all the 9-leaf trees obtained.
3. We chose a number of best fitting 9-leaf trees, for each added an admixture event by detaching any edge and choosing any two edges as its new parents and fitted all the 1-admixture graphs obtained. We removed all the duplicate graphs.
4. We chose six best fitting 1-admixture graphs, for each added another admixture event by detaching any edge and choosing any two edges as its new parents and fitted all the 2-admixture graphs obtained. We removed all the duplicate graphs, as well as any graph where BLK appeared as an admixed population.

The number of graphs to be expanded to more complex graphs after the first three steps was chosen to keep the search space manageable. The results of $f_2$- and $f_4$-analyses are shown in Fig. S22 and Fig. S23, respectively.

The results show a clear preference to an ancient gene flow from the ancestors of ABC-BC bears into the ancestors of all polar bears. In 1-admixture graphs of Fig. S22B, S23B the direction from polar bears into brown bears nevertheless fits well also, represented by the 3rd and 6th best fits, respectively. The outstanding best fit among the 2-admixture graphs in Figs. 4D, S22C, and S23C features bidirectional gene flow, but as the position of EBB is not consistent with the species tree, and there was no evidence of admixture between EBB and polar bears in other analyses or the literature, this is likely a result of ILS and ghost admixture to EBB. Beyond that, explicitly bidirectional gene flow is represented by the 4th graph for the $f_2$-statistics (Fig. S22C) and 5th graph for the $f_4$-statistics (Fig. S23C) best fits only.

**S21. TreeMix analysis**

TreeMix estimates the historical relationships among populations from genome-wide allele frequency data. The program generates a maximum likelihood graph that allows both population splits and migration events, and a residual heat map for visualizing the fit of the model to the data (81). We conducted TreeMix analyses to infer the relationships, divergence, and major mixtures between populations based on dataset DS3 (all SNPs from genomes > 8X coverage, excluding private alleles) and DS8 (DS3 but transversions only) (Table S5). We applied the option "-root BLK", which sets the American black bear as the position of the root, and "-noss", which turns off sample size correction. We allowed up to five migration edges on the tree (m ranging from 0 to 5) and generated the residual heatmap to identify populations that are not well-modeled after adding each migration edge (Fig. S24). The percentages of variation explained by the maximum likelihood trees were also calculated (Table S9).

The results obtained both from DS3 and DS8 with 0 migration events, which captured 99.77% of the variance, recapitulated the phylogenetic relationships among our bear samples resolved by RAxML analysis (section S4 above, Fig. S7). However, 0.23% of the variance could not be explained by the best-fit tree with 0 migration events. For both datasets, the first migration event at m = 1, representing 0.19% of the data, is admixture from ABC-BC bears into the ancestral node of the polar bear lineage. For DS3, results with m = 2 to m = 3 include admixture among brown bears, including EBB to BB and EBB to the ancestor of the ABC brown bears, while m = 4 and m = 5 includes admixture edges from outside the brown bear-polar bear lineage into EBB and APB, respectively. The results obtained from DS8 (Fig. S24B) captures the admixture edge from outside the brown bear-polar bear lineage into EBB at m = 2, while m = 5 includes an admixture edge from a brown ancestor into APB. Otherwise, results show similar admixture among brown bears as with DS3.

We repeated the TreeMix analyses with the same two datasets but excluding the ancient polar bear (APB), to investigate if including APB made a difference regarding admixture among polar bears and brown bears (Fig. S25). When excluding APB, the maximum likelihood trees with 0 migration events explained slightly less of the variation (99.63% and 99.67%, respectively; Table S9) than in the analyses with APB included. However, again most of the residual variation was explained by the first migration edge from ABC-BC and into the polar bear lineage.

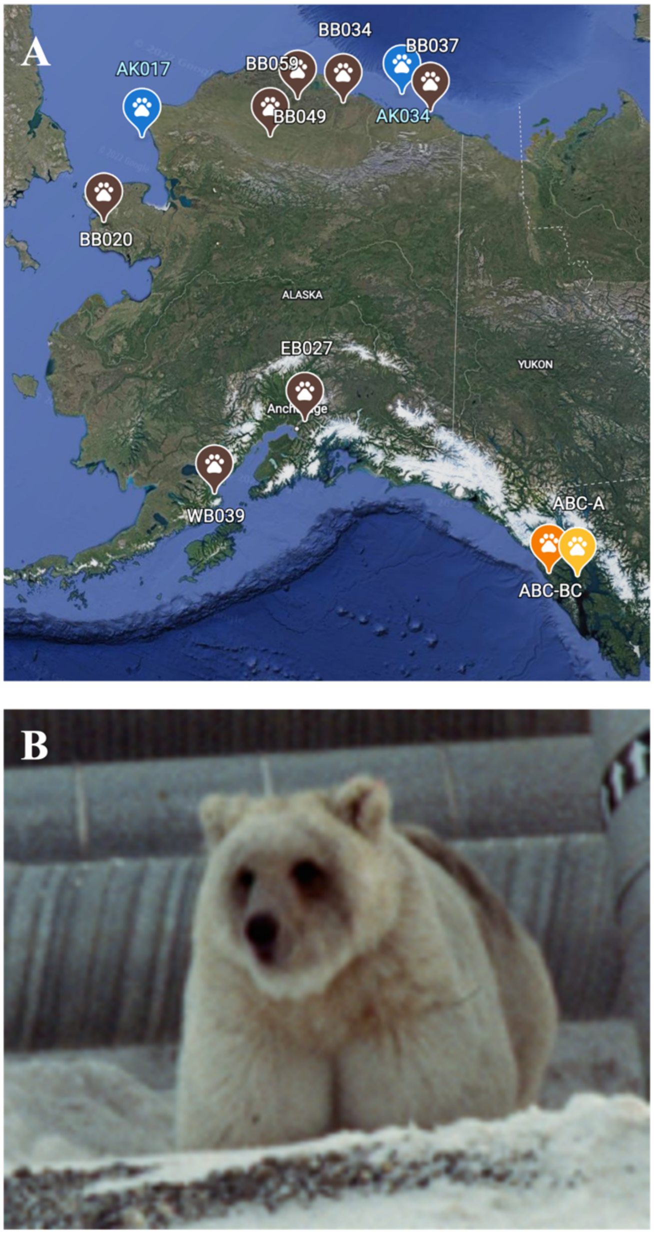


**Fig. S1. A.** Geographic localities of the two polar bears and seven brown bears from Alaska selected for genome sequencing in this study (see also Table S1). **B.** Male brown bear (BB034) from North Slope, Alaska. Photo: D. Shideler.

**A.**

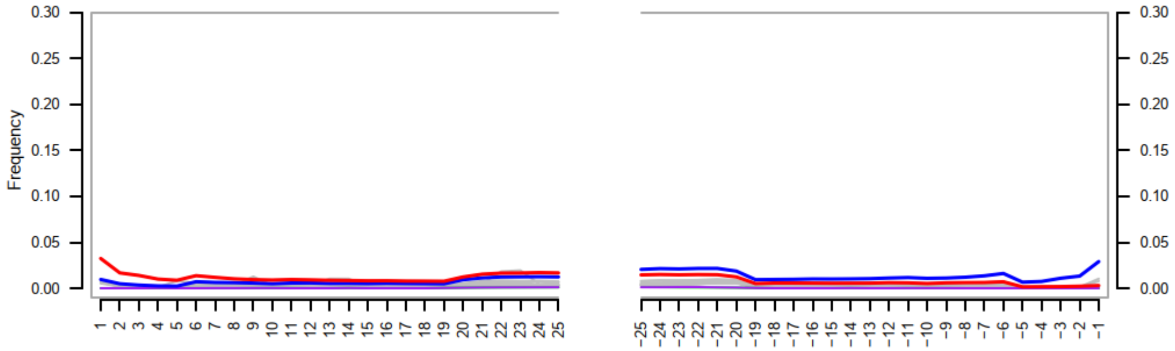
Frequency
0.30
0.25
0.20
0.15
0.10
0.05
0.00
1 2 3 4 5 6 7 8 9 10 11 12 13 14 15 16 17 18 19 20 21 22 23 24 25
−25 −24 −23 −22 −21 −20 −19 −18 −17 −16 −15 −14 −13 −12 −11 −10 −9 −8 −7 −6 −5 −4 −3 −2 −1

**B.**

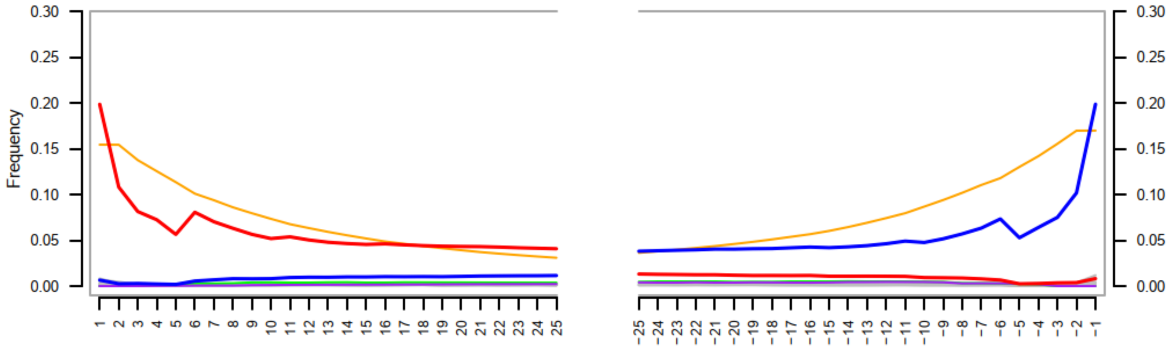
Frequency
0.30
0.25
0.20
0.15
0.10
0.05
0.00
1 2 3 4 5 6 7 8 9 10 11 12 13 14 15 16 17 18 19 20 21 22 23 24 25
−25 −24 −23 −22 −21 −20 −19 −18 −17 −16 −15 −14 −13 −12 −11 −10 −9 −8 −7 −6 −5 −4 −3 −2 −1

**C.**

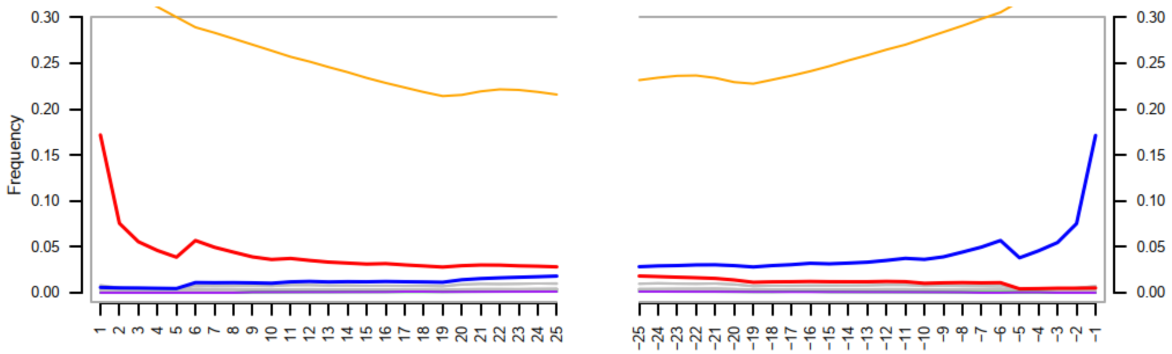
Frequency
0.30
0.25
0.20
0.15
0.10
0.05
0.00
1 2 3 4 5 6 7 8 9 10 11 12 13 14 15 16 17 18 19 20 21 22 23 24 25
−25 −24 −23 −22 −21 −20 −19 −18 −17 −16 −15 −14 −13 −12 −11 −10 −9 −8 −7 −6 −5 −4 −3 −2 −1

**D.**

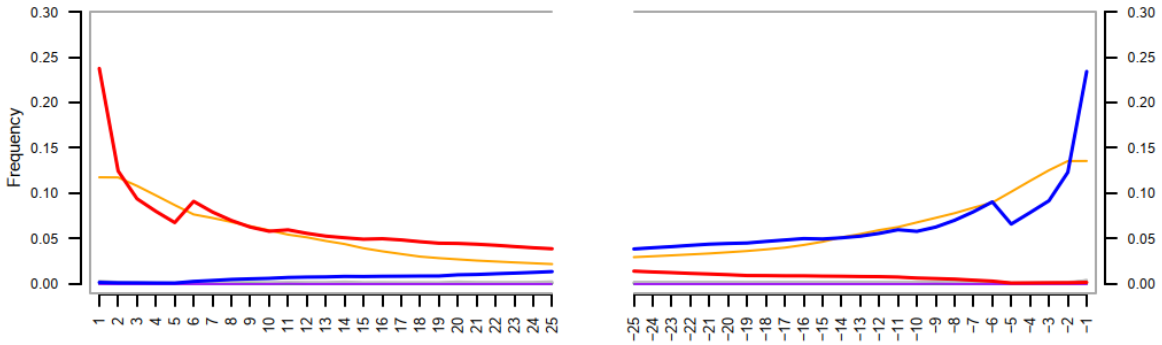
Frequency
0.30
0.25
0.20
0.15
0.10
0.05
0.00
1 2 3 4 5 6 7 8 9 10 11 12 13 14 15 16 17 18 19 20 21 22 23 24 25
−25 −24 −23 −22 −21 −20 −19 −18 −17 −16 −15 −14 −13 −12 −11 −10 −9 −8 −7 −6 −5 −4 −3 −2 −1

**E.**

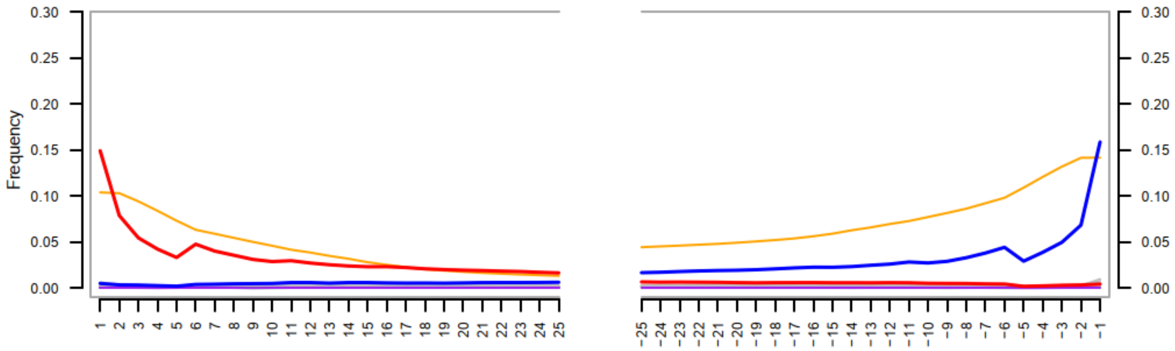

Frequency
0.30
0.25
0.20
0.15
0.10
0.05
0.00


**F.**

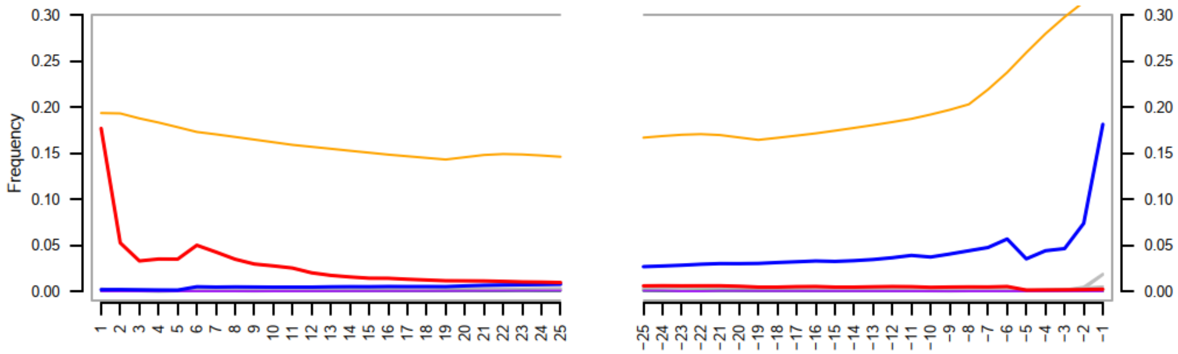

Frequency
0.30
0.25
0.20
0.15
0.10
0.05
0.00


**G.**

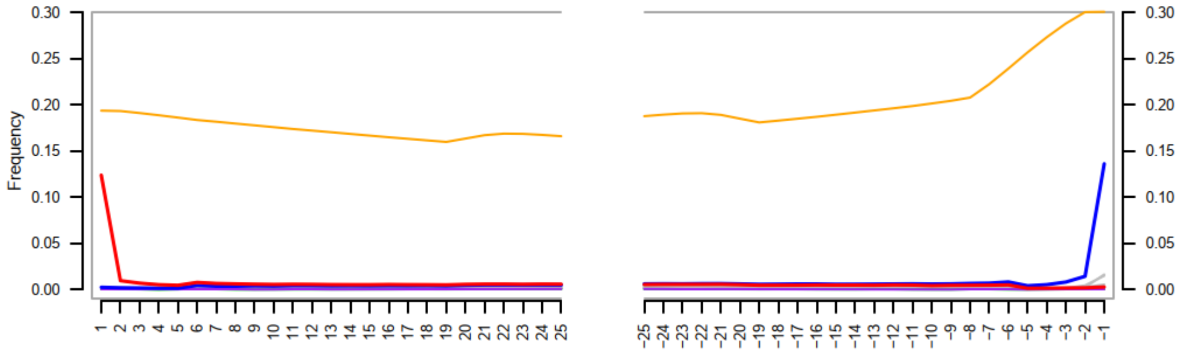

Frequency
0.30
0.25
0.20
0.15
0.10
0.05
0.00

**H.**

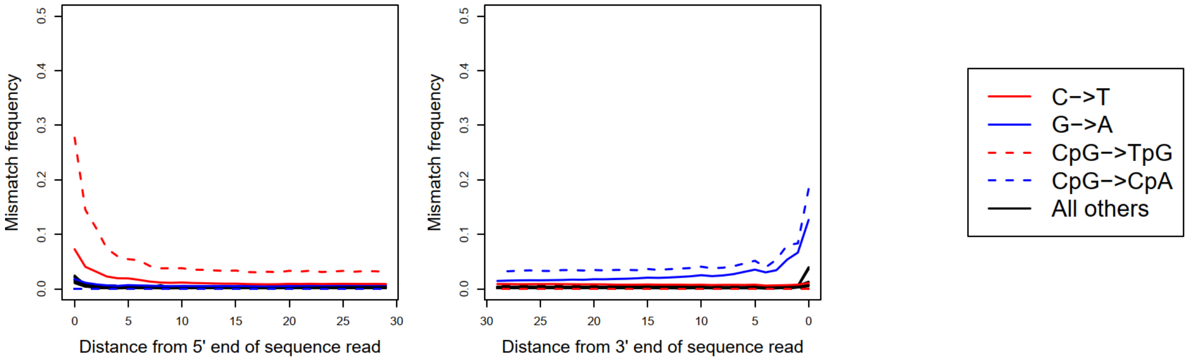
Mismatch frequency
0.0
0.1
0.2
0.3
0.4
0.5
0
5
10
15
20
25
30
Distance from 5' end of sequence read
Mismatch frequency
30
25
20
15
10
5
0
Distance from 3' end of sequence read
C−>T
G−>A
CpG−>TpG
CpG−>CpA
All others

**I.**

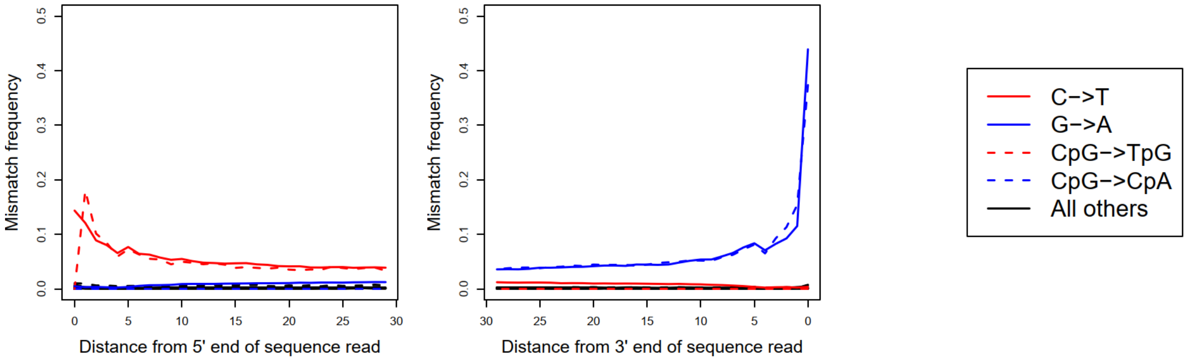
Mismatch frequency
0.0
0.1
0.2
0.3
0.4
0.5
0
5
10
15
20
25
30
Distance from 5' end of sequence read
Mismatch frequency
30
25
20
15
10
5
0
Distance from 3' end of sequence read
C−>T
G−>A
CpG−>TpG
CpG−>CpA
All others

**J.**

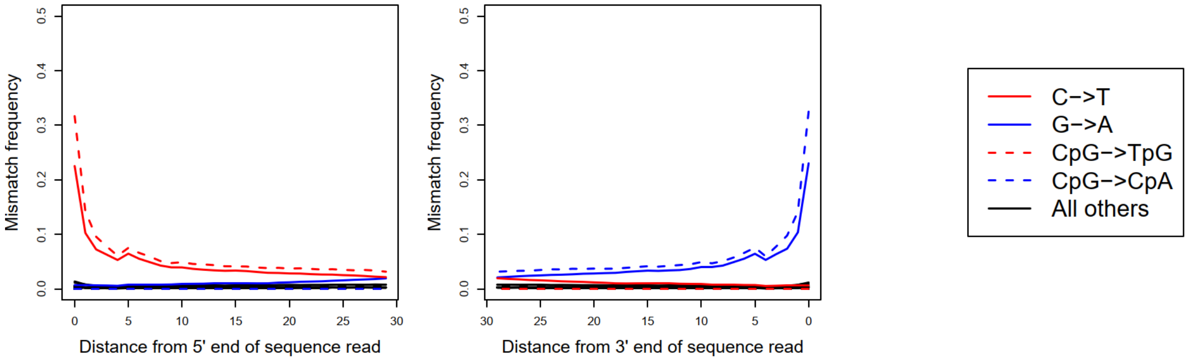
Mismatch frequency
0.0
0.1
0.2
0.3
0.4
0.5
0
5
10
15
20
25
30
Distance from 5' end of sequence read
Mismatch frequency
30
25
20
15
10
5
0
Distance from 3' end of sequence read
C−>T
G−>A
CpG−>TpG
CpG−>CpA
All others

**K.**

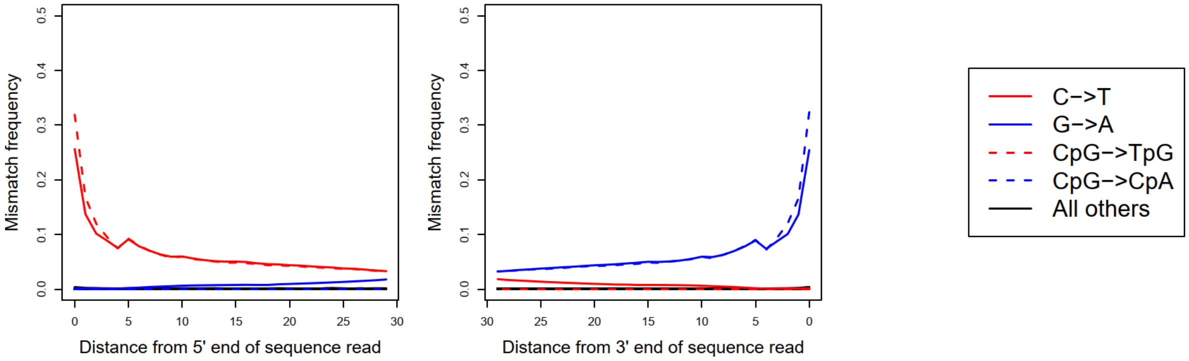
Mismatch frequency
Distance from 5' end of sequence read
Distance from 3' end of sequence read
C−>T
G−>A
CpG−>TpG
CpG−>CpA
All others

**L.**

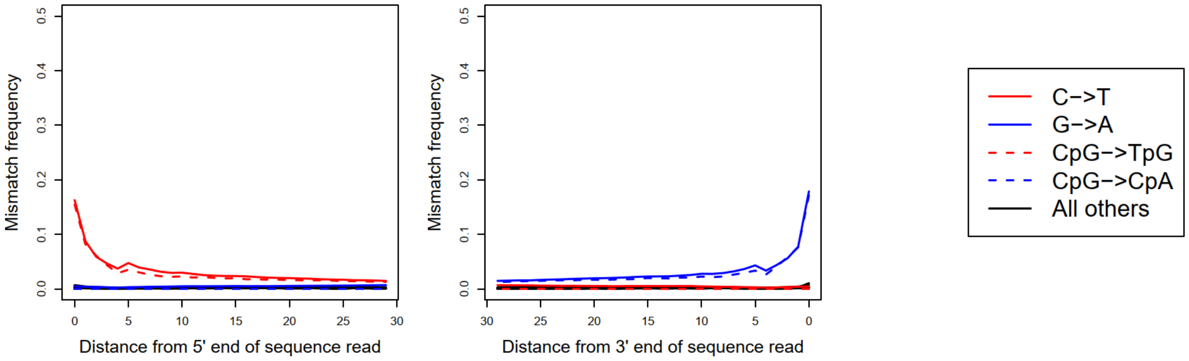
Mismatch frequency
Distance from 5' end of sequence read
Distance from 3' end of sequence read
C−>T
G−>A
CpG−>TpG
CpG−>CpA
All others

**M.**

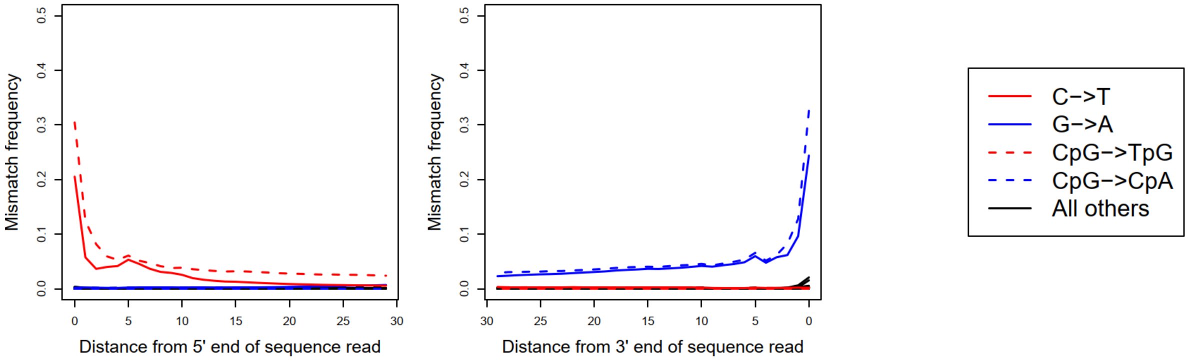
Mismatch frequency
Distance from 5' end of sequence read
Distance from 3' end of sequence read
C−>T
G−>A
CpG−>TpG
CpG−>CpA
All others

**N.**

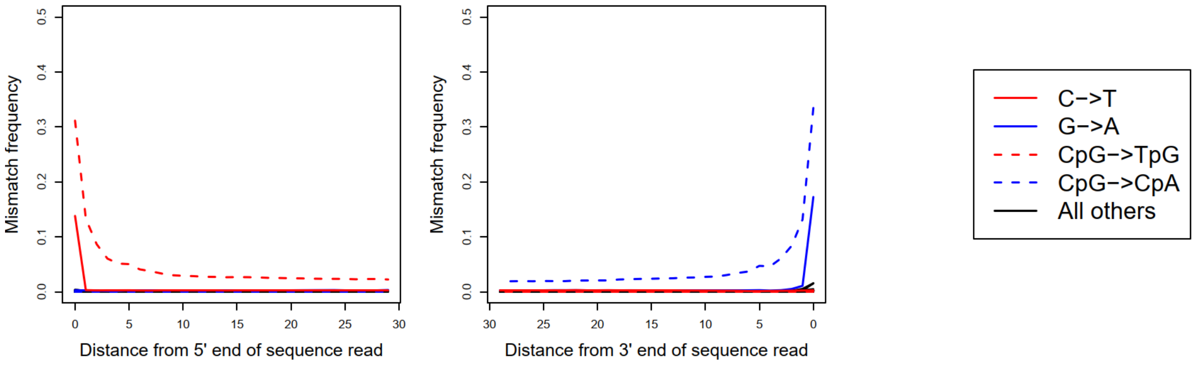


**Fig. S2.** Postmortem DNA damage profiles displaying misincorporation patterns (frequencies of C→T at 5'-end and G→A at 3'-end) for each set of sequence libraries (no end trimming) of the ancient polar bear (see Table S2): APBv2, WGE, EL09, EL10, EL11, EL14, EL15. The plots show the proportions of mapped sequence reads that possess a T compared to a C in the polar bear reference genome (red line), or an A compared to a G in the reference (blue line) at each nucleotide position for the first 25 nucleotides, with 5' end shown to the left and 3' end to the right. The damage profiles were generated using mapDamage2 (**A.** APBv2, **B.** WGE, **C.** EL09, **D.** EL10, **E.** EL11, **F.** EL14, **G.** EL15) and PMDtools (**H.** APBv2, **I.** WGE, **J.** EL09, **K.** EL10, **L.** EL11, **M.** EL14, **N.** EL15).

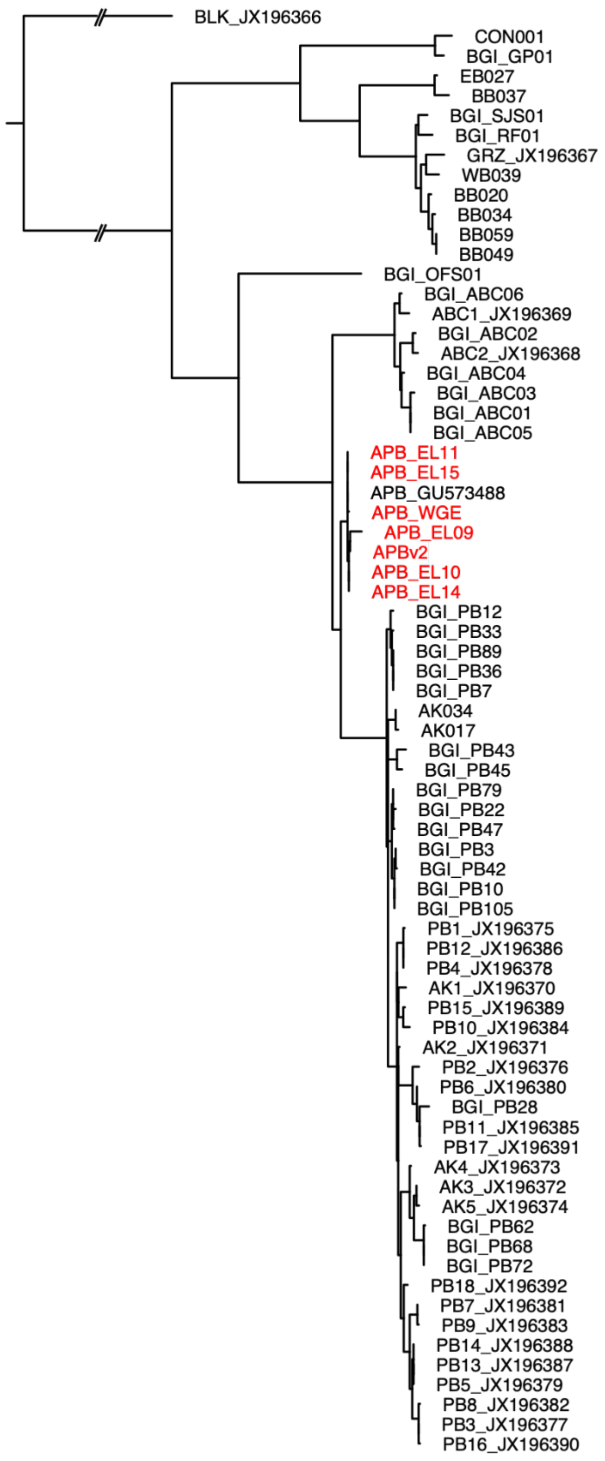


**Fig. S3.** Authentication of APB sequencing libraries. RAxML phylogenetic tree of mitogenome data based on assemblies from each of the DNA libraries constructed for the ancient polar bear combined with the full mitogenome dataset of brown and polar bears included in this study. Assembled mitogenome data from each library group together in a single, well-supported lineage, sister to the modern polar bears, and together with the published mitogenome of the ancient polar bear (GU573488).

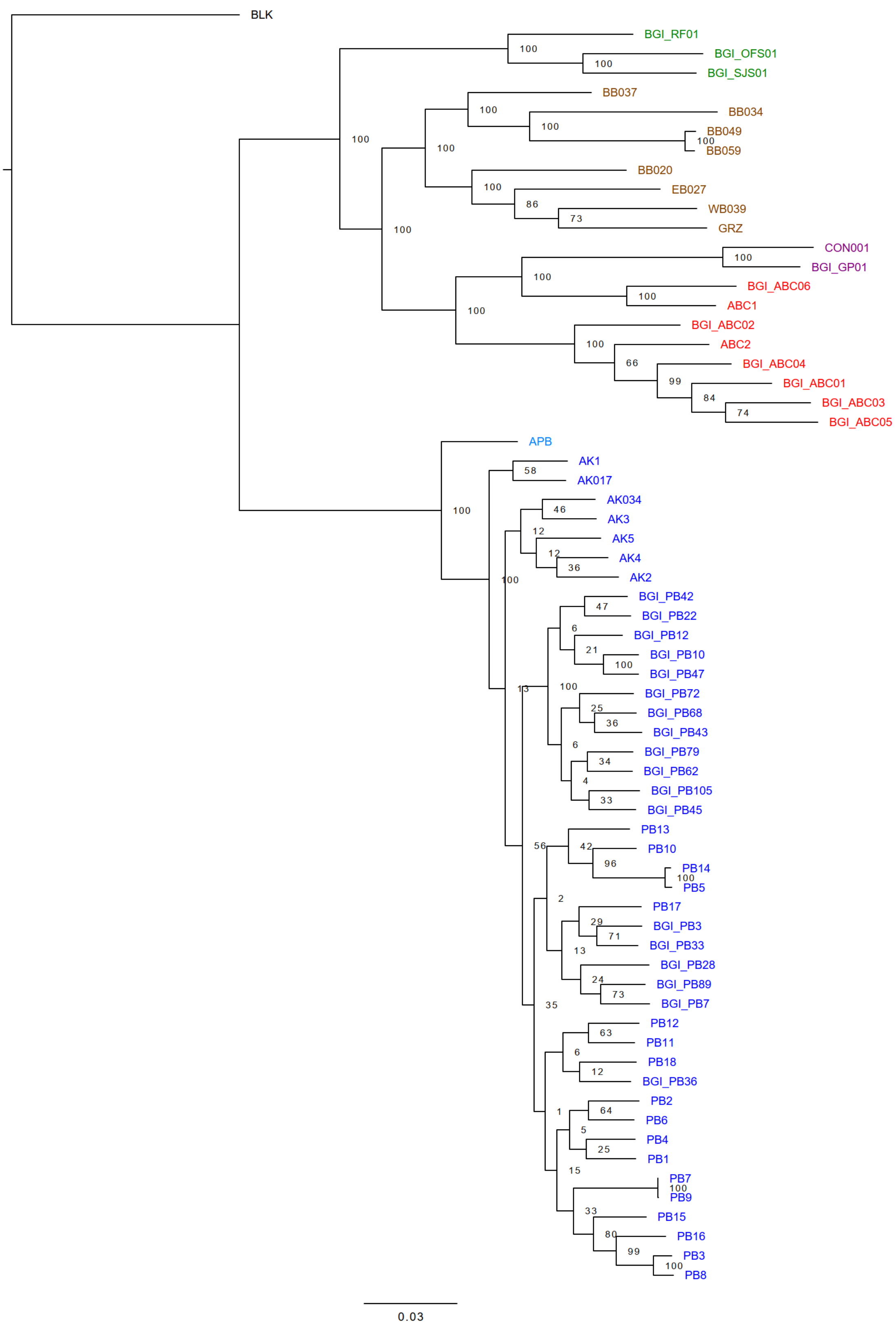


**Fig. S4.** Maximum likelihood phylogenetic tree based on autosomal SNPs (DS7).

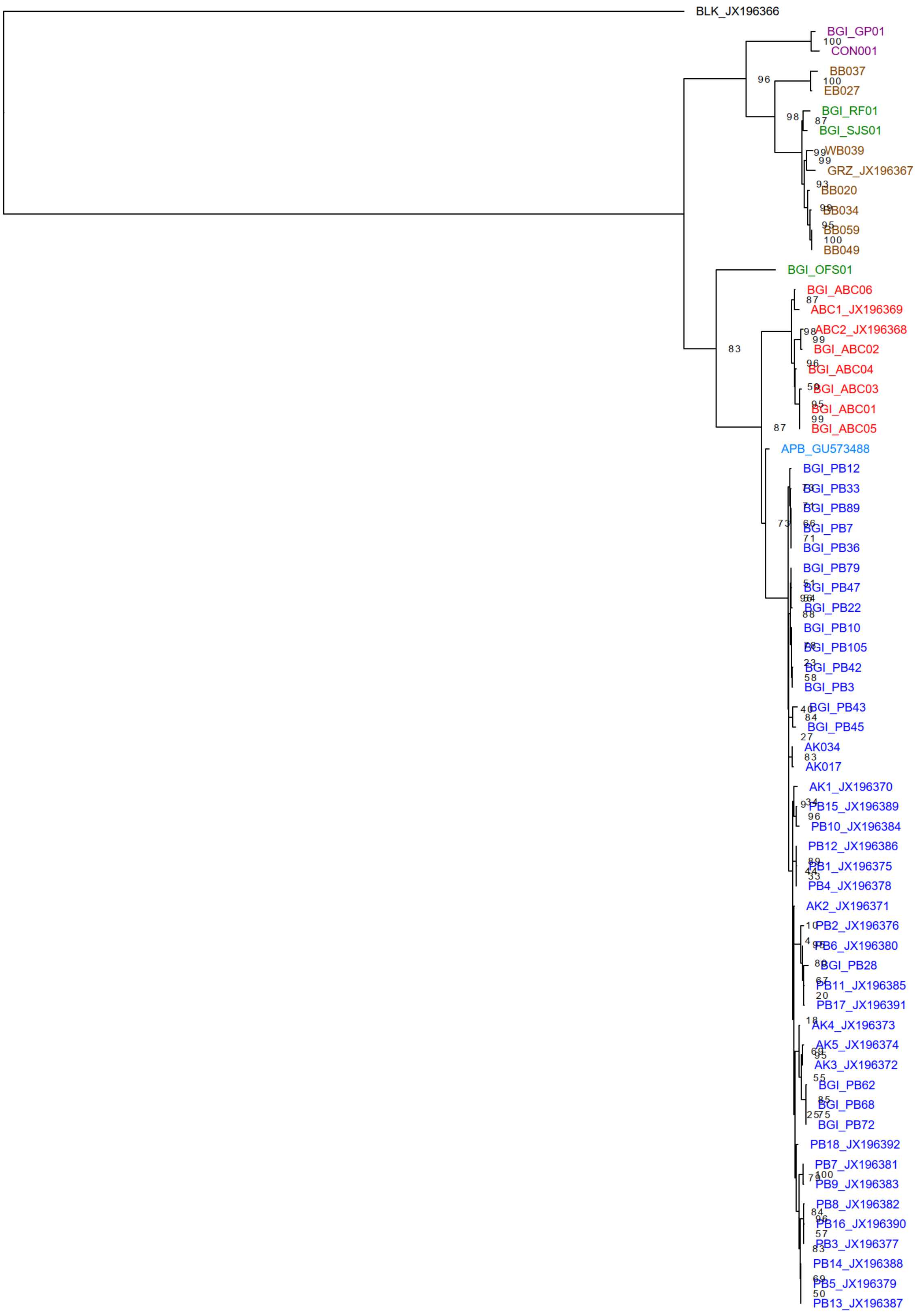


**Fig. S5.** Maximum likelihood phylogenetic tree based on mitogenomes.

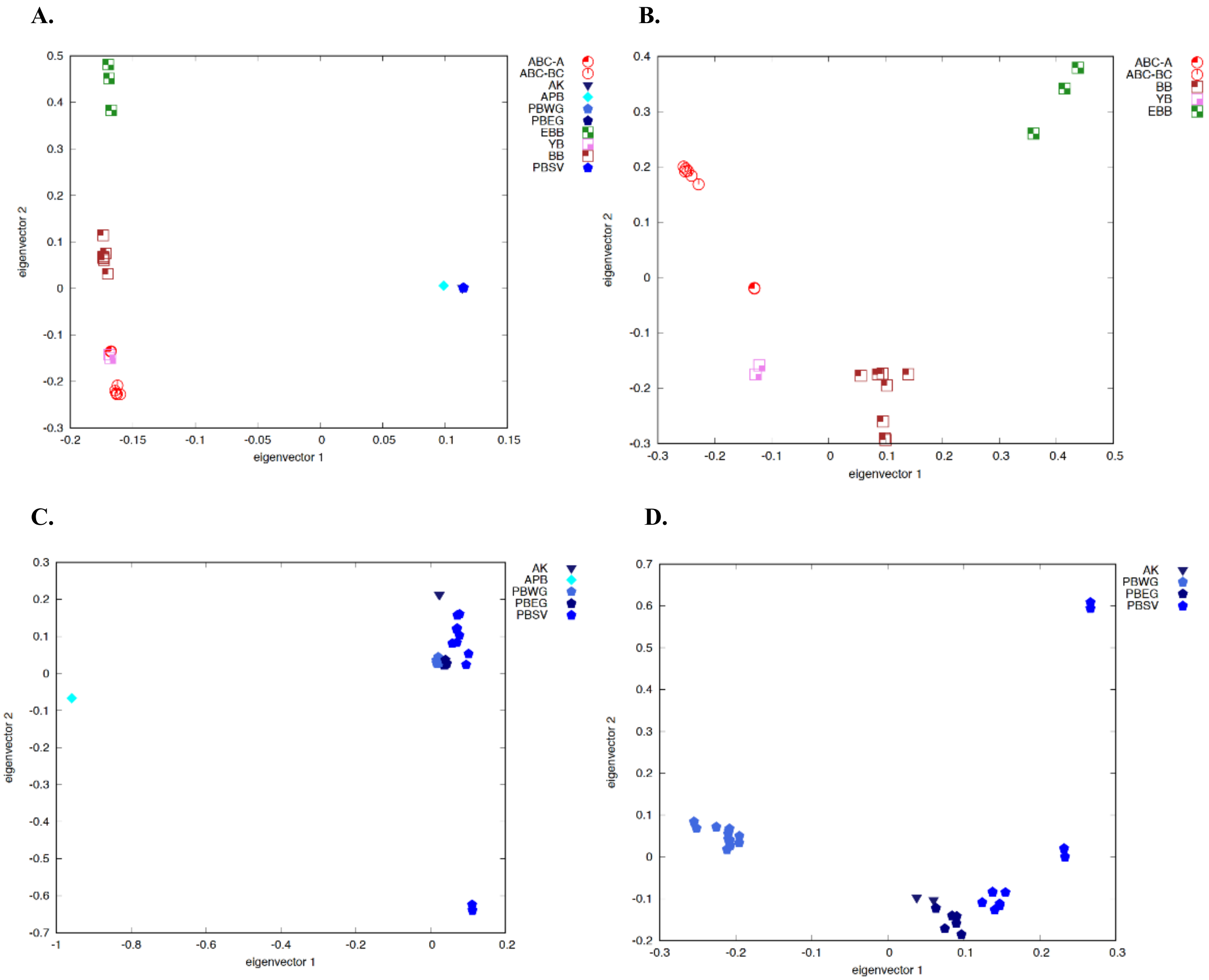


**Fig. S6.** Principal component analyses of the brown and polar bear genomes with a coverage > 8x (DS3). **A.** All brown and polar bears. **B.** Brown bears only. **C.** Polar bears only. **D.** Modern polar bears only.

**A.**

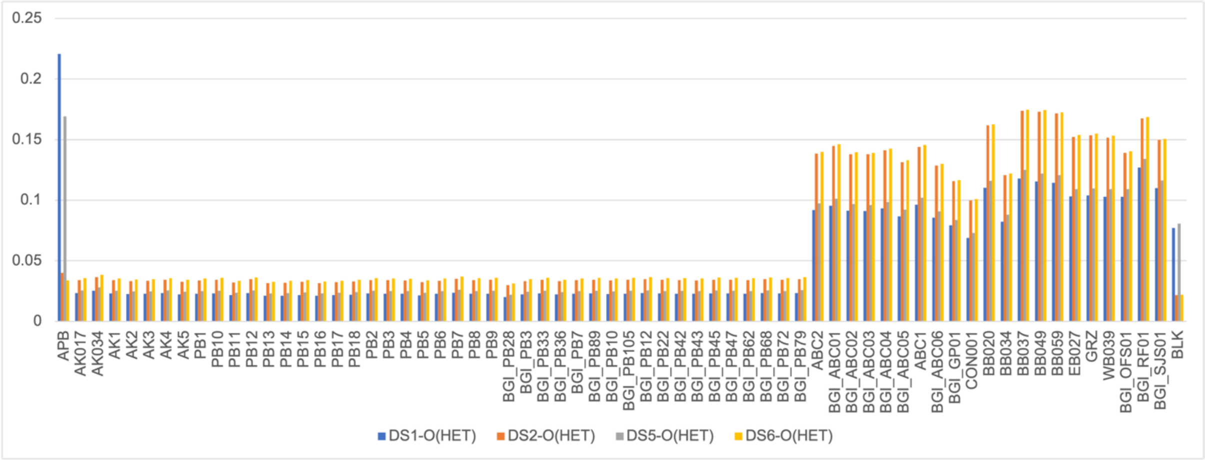


**B.**

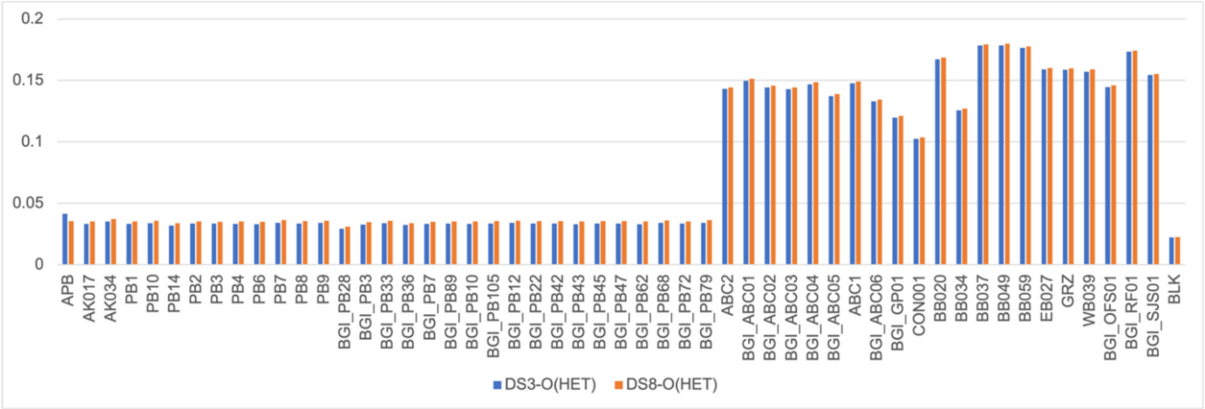


**C.**

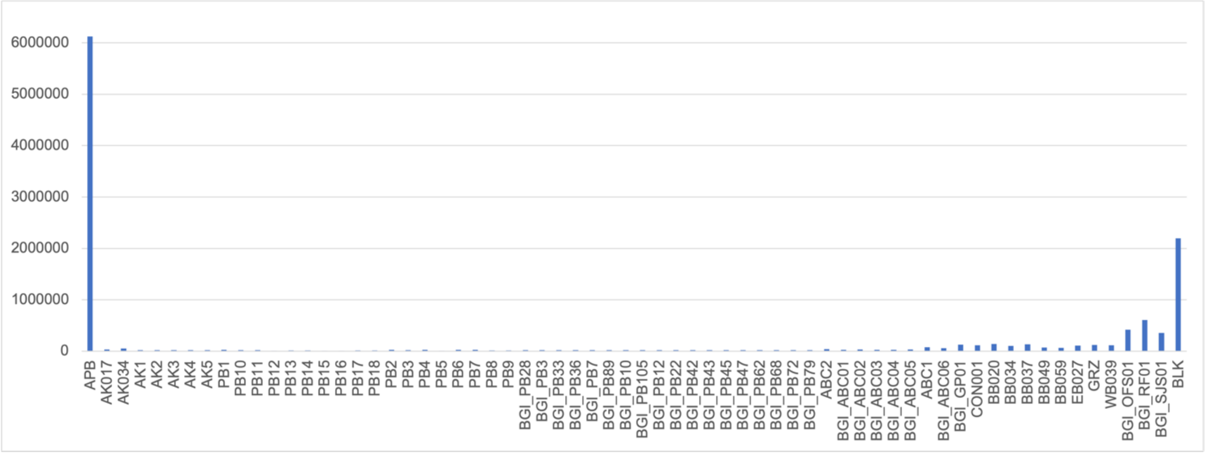


**Fig. S7. A.** Autosomal heterozygosity frequencies for polar and brown bear genomes calculated for the four datasets DS1, DS2, DS5, and DS6 and **B.** datasets DS3 and DS8 (see Table S5). **C.** Number of private alleles (here, unique alleles only found in the ancient polar bear compared to the modern polar bears).

**A.**

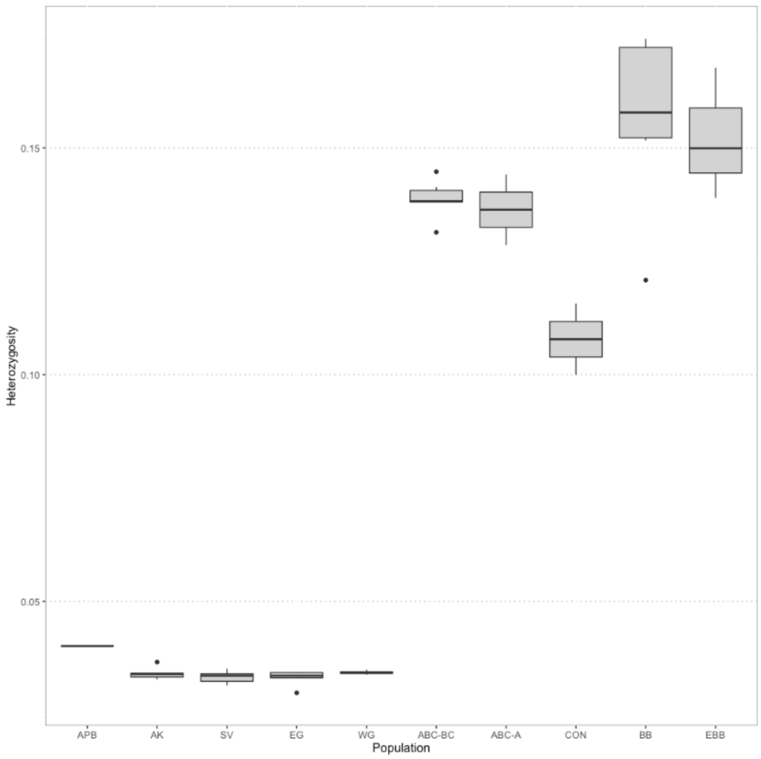


**B.**

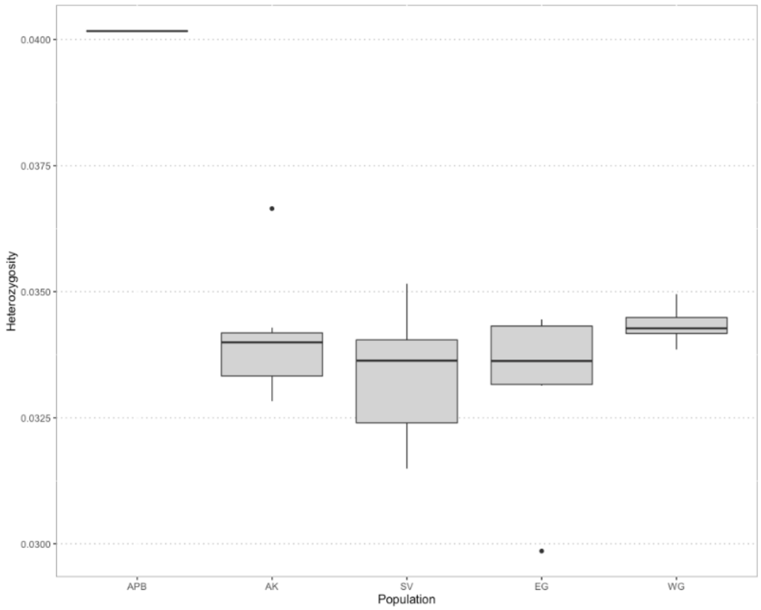


**Fig. S8.** Autosomal heterozygosity frequencies across polar and brown bear populations calculated for dataset DS2: **A.** All populations. **B.** Polar bear populations only.

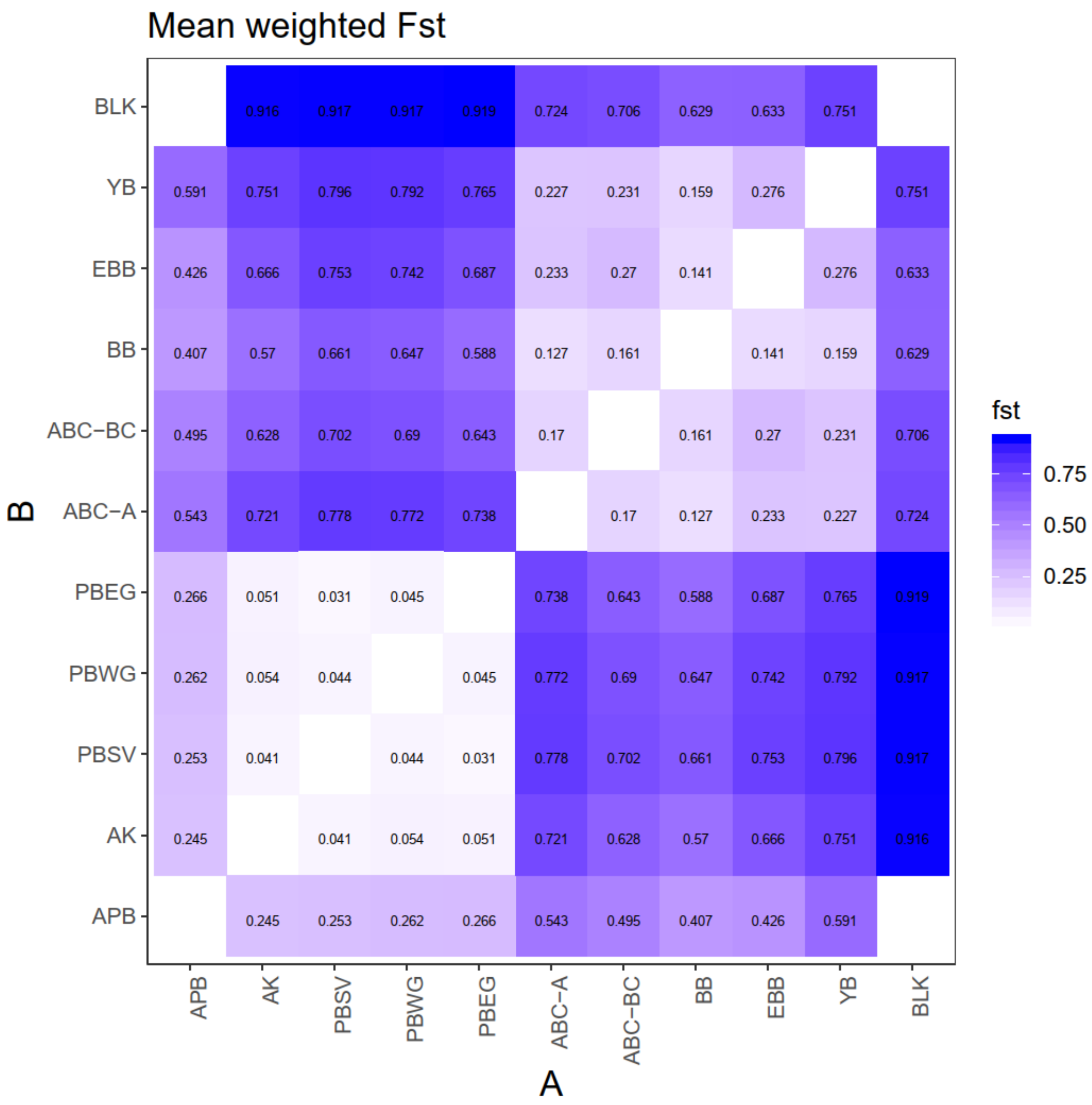


**Fig. S9.** Fixation indices $F_{ST}$ among polar bear and brown bear populations.

A.

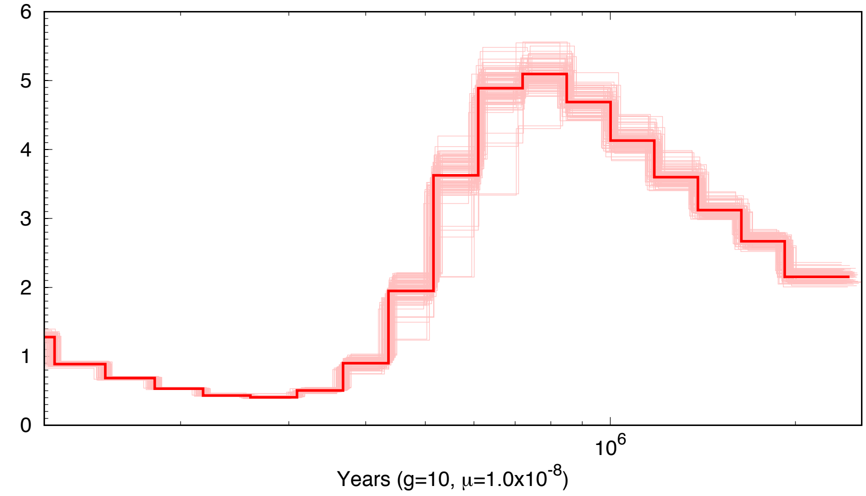

Effective population size (x10^4)
Years (g=10, μ=1.0x10^-8)


B.

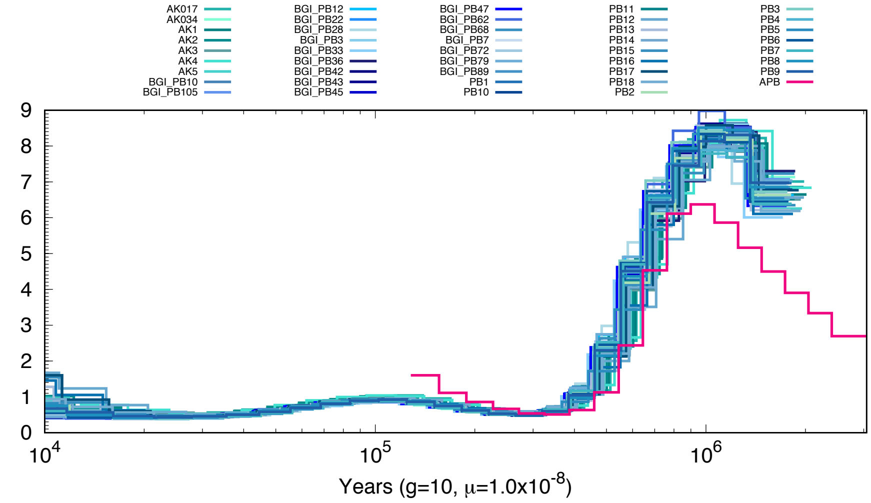

AK017
AK034
AK1
AK2
AK3
AK4
AK5
BGI_PB10
BGI_PB105
BGI_PB12
BGI_PB22
BGI_PB28
BGI_PB3
BGI_PB33
BGI_PB36
BGI_PB42
BGI_PB43
BGI_PB45
BGI_PB47
BGI_PB62
BGI_PB68
BGI_PB7
BGI_PB72
BGI_PB79
BGI_PB89
PB1
PB10
PB11
PB12
PB13
PB14
PB15
PB16
PB17
PB18
PB2
PB3
PB4
PB5
PB6
PB7
PB8
PB9
APB
Effective population size (x10^4)
Years (g=10, μ=1.0x10^-8)


C.

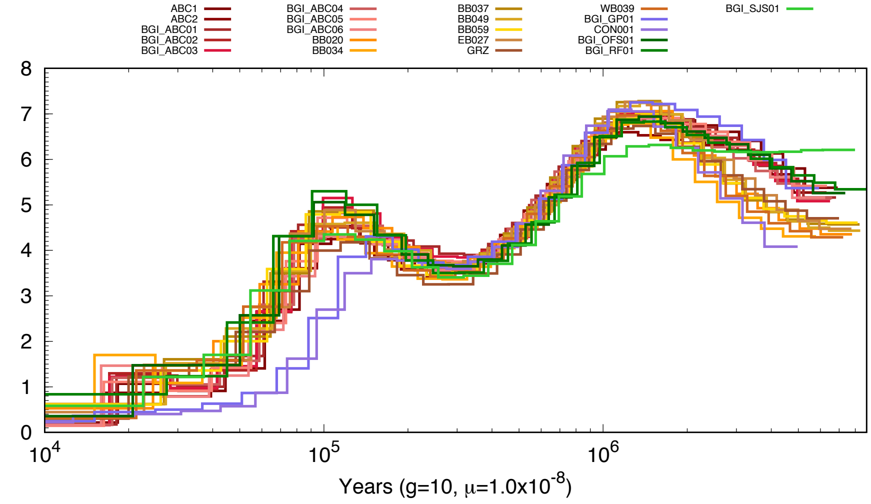

ABC1
ABC2
BGI_ABC01
BGI_ABC02
BGI_ABC03
BGI_ABC04
BGI_ABC05
BGI_ABC06
BB020
BB034
BB037
BB049
BB059
EB027
GRZ
WB039
BGI_GP01
CON001
BGI_OFS01
BGI_RF01
BGI_SJS01
Effective population size (x10^4)
Years (g=10, μ=1.0x10^-8)

**D.**

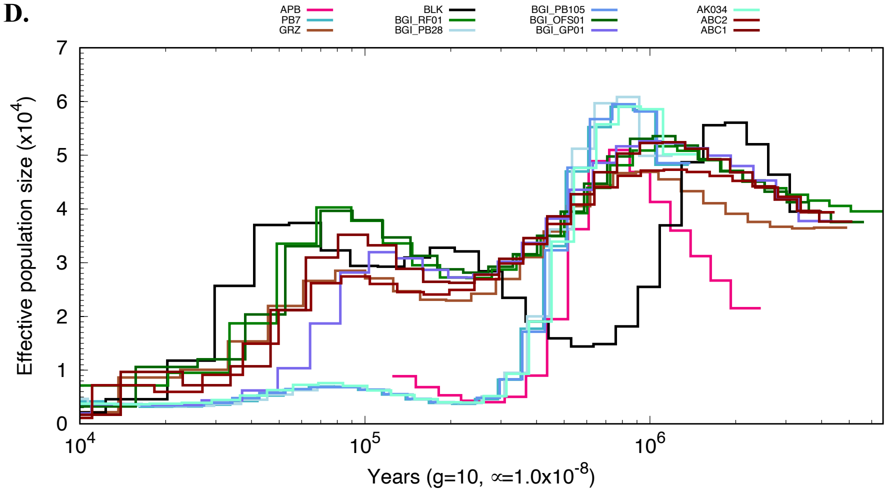


**E.**

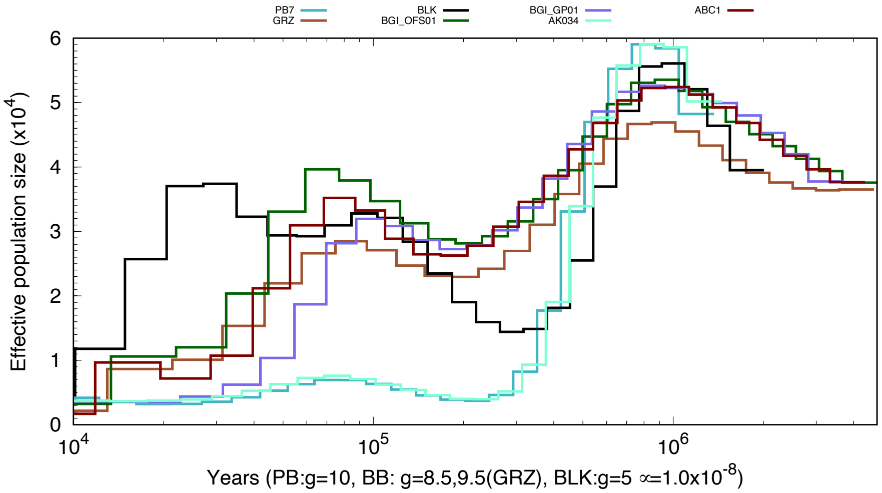


**Fig. S10.** Demographic history inferred using PSMC. Estimates of effective population size over time are shown for the following populations plotted using g = 10: **A.** The ancient polar bear bootstrap replicates. **B.** Ancient and modern polar bears. **C.** Brown bears. **D–E.** One individual from each of the populations with modern bears downsampled to ~10X using different generation times reflecting differences among bears.

**A.**

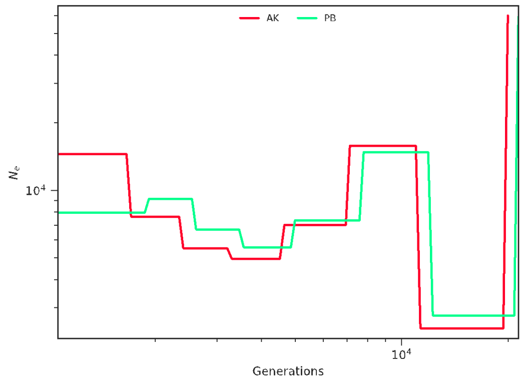


**B.**

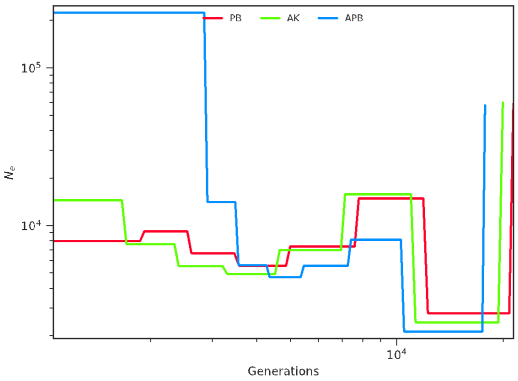


**C.**

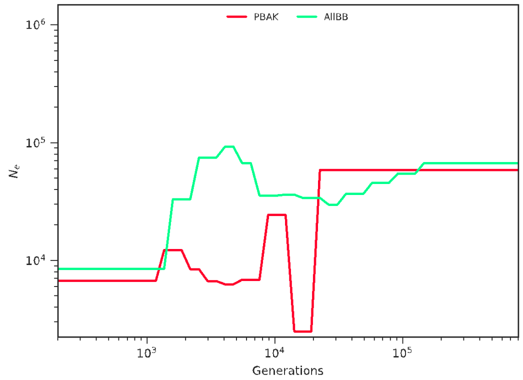


**D.**

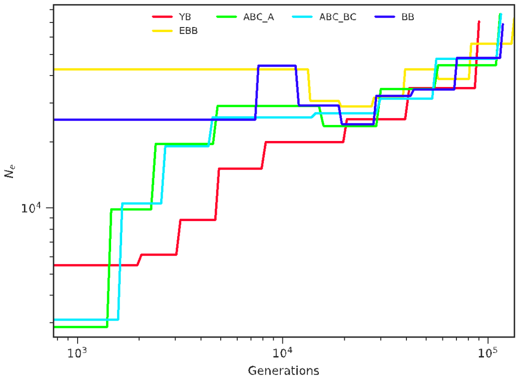


**E.**

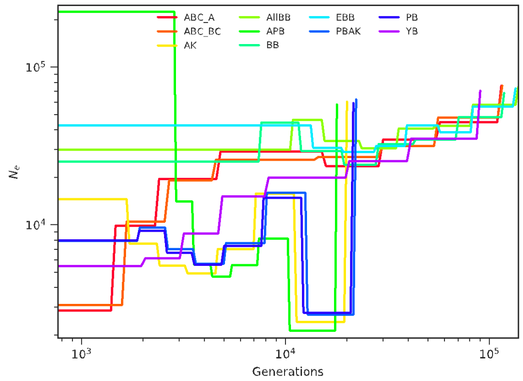


**Fig. S11.** Demographic history inferred using SMC++. Estimates of effective population size over generations are shown for the following populations: **A.** Modern polar bears (AK denotes polar bears from Alaska, whereas PB denotes polar bears from Svalbard and Greenland). **B.** Ancient (APB) and modern polar bears. **C.** All modern polar bears and brown bears. **D.** All populations of brown bear. **E.** All bears plotted together.

**A.**

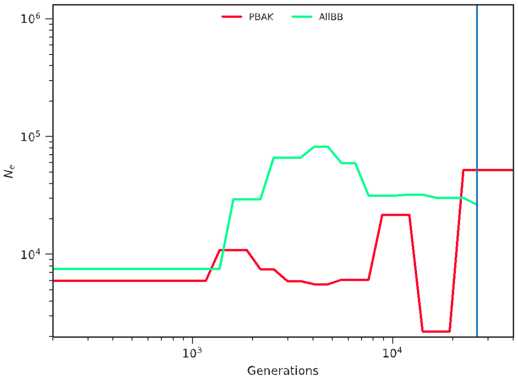

PBAK
AllBB
$10^6$
$10^5$
$10^4$
$N_e$
$10^3$
$10^4$
Generations


**B.**

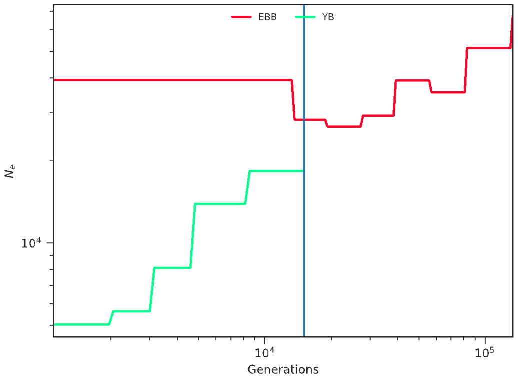

EBB
YB
$10^4$
$N_e$
$10^4$
$10^5$
Generations


**C.**

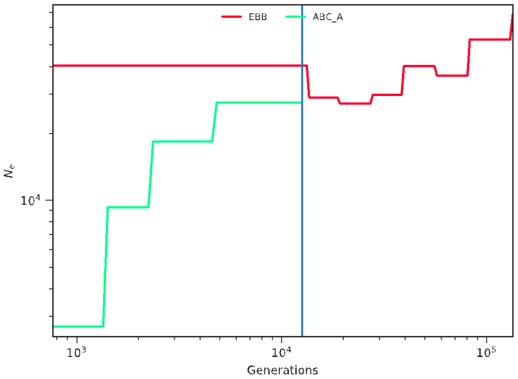

EBB
ABC_A
$10^4$
$N_e$
$10^3$
$10^4$
$10^5$
Generations


**D.**

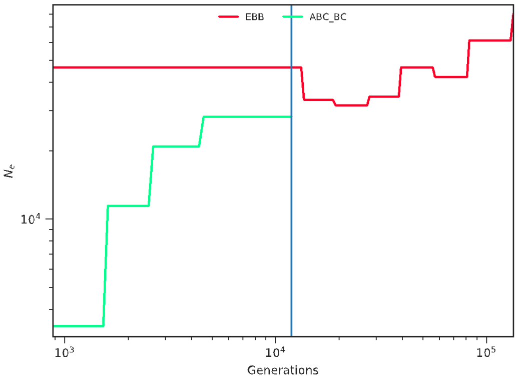

EBB
ABC_BC
$10^4$
$N_e$
$10^3$
$10^4$
$10^5$
Generations


**E.**

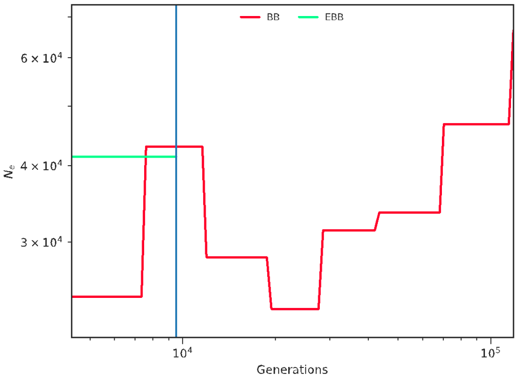

BB
EBB
$6 \times 10^4$
$4 \times 10^4$
$3 \times 10^4$
$N_e$
$10^4$
$10^5$
Generations


**F.**

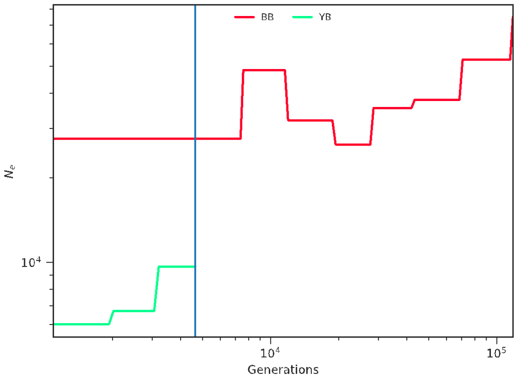

BB
YB
$10^4$
$N_e$
$10^4$
$10^5$
Generations

**G.**

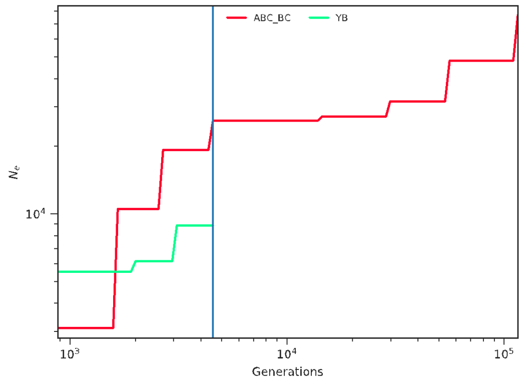


**H.**

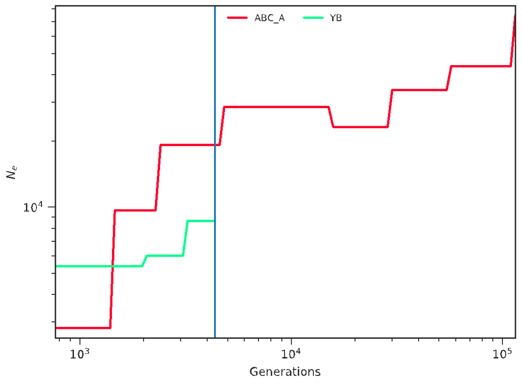


**I.**

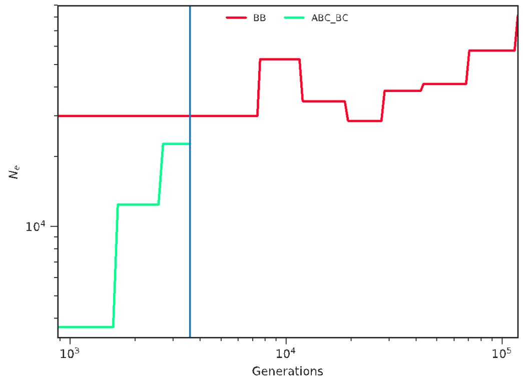


**J.**

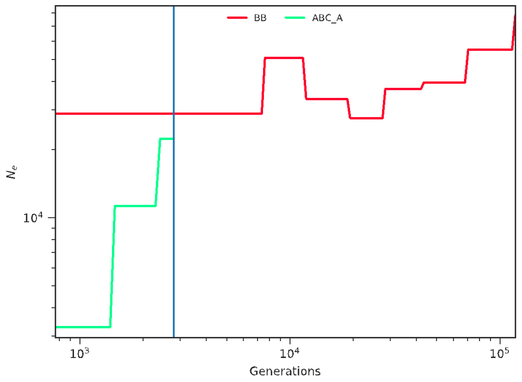


**K.**

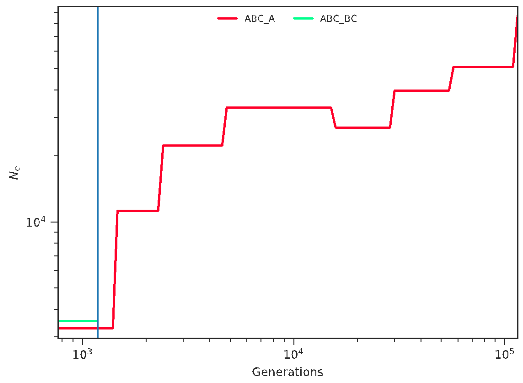


**L.**

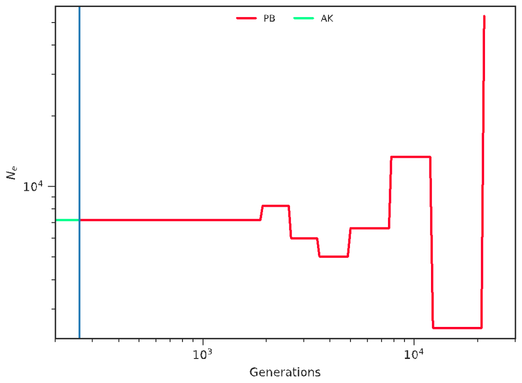


**Fig. S12.** Inference of divergence times of pairs of diverged populations estimated with SMC++. Shown are estimates of split times of the following pairs of populations: **A.** All polar bears vs brown bears. **B.** European (EBB) vs continental NA (YB) brown bears. **C.** European (EBB) vs Admiralty Isl. (ABC-A) brown bears. **D.** European (EBB) vs Baranof/Chichagof (ABC-BC) brown bears. **E.** European (EBB) vs Alaskan mainland (BB) brown bears. **F.** Alaskan mainland (BB) vs continental NA (YB) brown bears. **G.** Baranof/Chichagof (ABC-BC) vs continental NA (YB) brown bears. **H.** Admiralty Isl. (ABC-A) continental NA (YB) brown bears. **I.** Alaskan mainland (BB) vs Baranof/Chichagof (ABC-BC) brown bears. **J.** Alaskan mainland (BB) vs Admiralty Isl. (ABC-A) brown bears. **K.** Baranof/Chichagof (ABC-BC) vs Admiralty Isl. (ABC-A) brown bears. **L.** Alaskan (AK) vs other modern (PB) polar bears.

**A.**

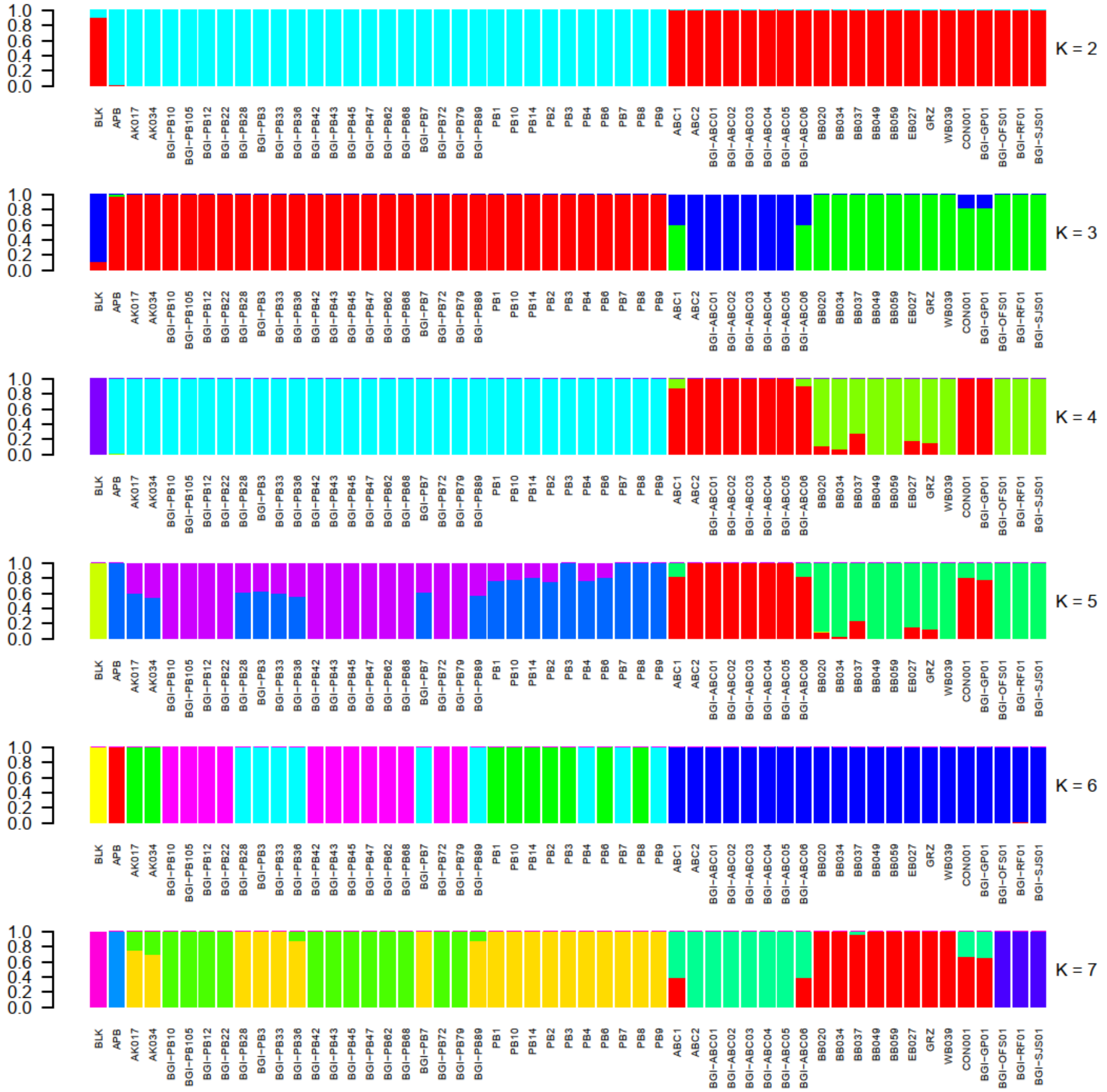
1.0
0.8
0.6
0.4
0.2
0.0
K = 2
K = 3
K = 4
K = 5
K = 6
K = 7
BLK
APB
AK017
AK034
BGI−PB10
BGI−PB105
BGI−PB12
BGI−PB22
BGI−PB28
BGI−PB3
BGI−PB33
BGI−PB36
BGI−PB42
BGI−PB43
BGI−PB45
BGI−PB47
BGI−PB62
BGI−PB68
BGI−PB7
BGI−PB72
BGI−PB79
BGI−PB89
PB1
PB10
PB14
PB2
PB3
PB4
PB6
PB7
PB8
PB9
ABC1
ABC2
BGI−ABC01
BGI−ABC02
BGI−ABC03
BGI−ABC04
BGI−ABC05
BGI−ABC06
BB020
BB034
BB037
BB049
BB059
EB027
GRZ
WB039
CON001
BGI−GP01
BGI−OFS01
BGI−RF01
BGI−SJS01

**B.**

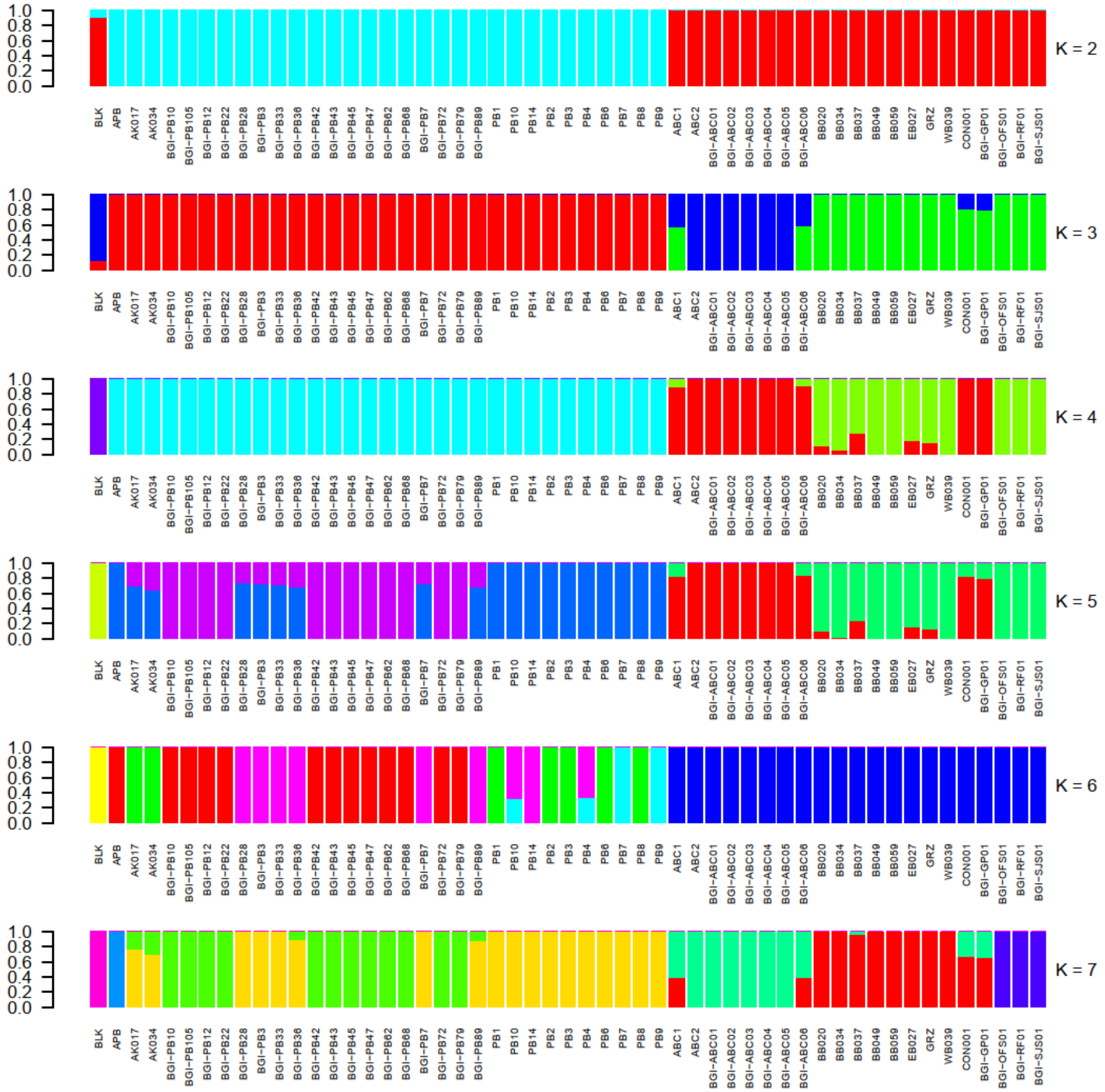

1.0
0.8
0.6
0.4
0.2
0.0
K = 2
K = 3
K = 4
K = 5
K = 6
K = 7
BLK
APB
AK017
AK034
BGI-PB10
BGI-PB105
BGI-PB12
BGI-PB22
BGI-PB28
BGI-PB3
BGI-PB33
BGI-PB36
BGI-PB42
BGI-PB43
BGI-PB45
BGI-PB47
BGI-PB62
BGI-PB68
BGI-PB7
BGI-PB72
BGI-PB79
BGI-PB89
PB1
PB10
PB14
PB2
PB3
PB4
PB6
PB7
PB8
PB9
ABC1
ABC2
BGI-ABC01
BGI-ABC02
BGI-ABC03
BGI-ABC04
BGI-ABC05
BGI-ABC06
BB020
BB034
BB037
BB049
BB059
EB027
GRZ
WB039
CON001
BGI-GP01
BGI-OFS01
BGI-RF01
BGI-SJS01


**C.**

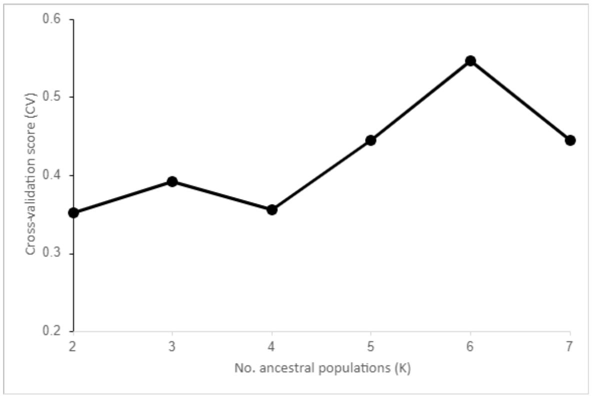

0.6
0.5
0.4
0.3
0.2
Cross-validation score (CV)
2
3
4
5
6
7
No. ancestral populations (K)

**Fig. S13.** ADMIXTURE analysis of brown bear, polar bear and American black bear genomes. Each vertical line represents an individual with the lengths of the colored segments proportional to the contributions of the ancestral components to the genome of the individual. Each panel displays the ancestral proportions with different numbers of hypothetical ancestral populations (K = 2 to K = 7). **A.** Genomes with > 8X coverage. **B.** Genomes with > 8X coverage and transversions only. **C.** Cross validation.

**A.**

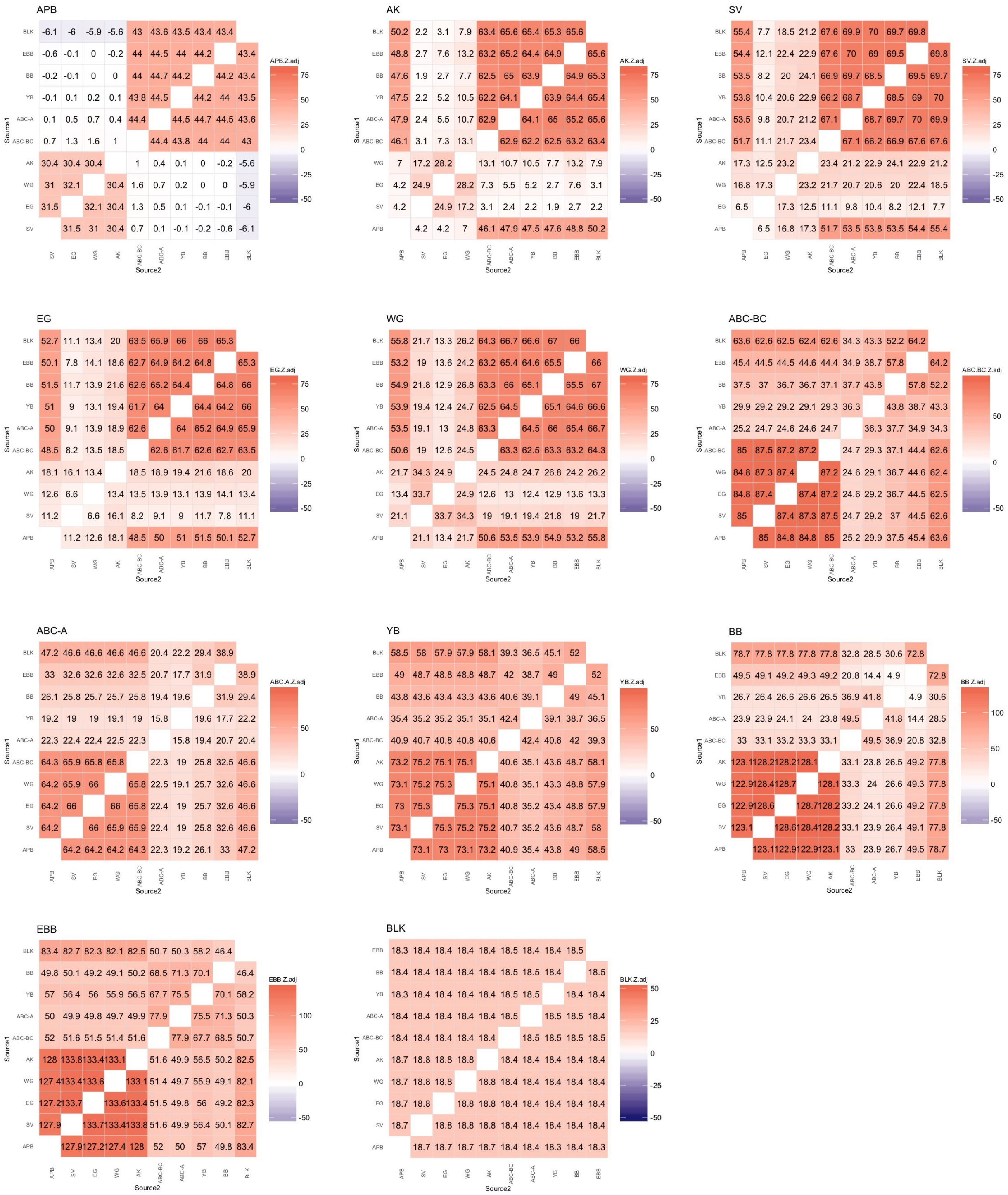
APB
APB.Z.adj
AK
AK.Z.adj
SV
SV.Z.adj
EG
EG.Z.adj
WG
WG.Z.adj
ABC-BC
ABC.BC.Z.adj
ABC-A
ABC.A.Z.adj
YB
YB.Z.adj
BB
BB.Z.adj
EBB
EBB.Z.adj
BLK
BLK.Z.adj
Source1
Source2

**B.**

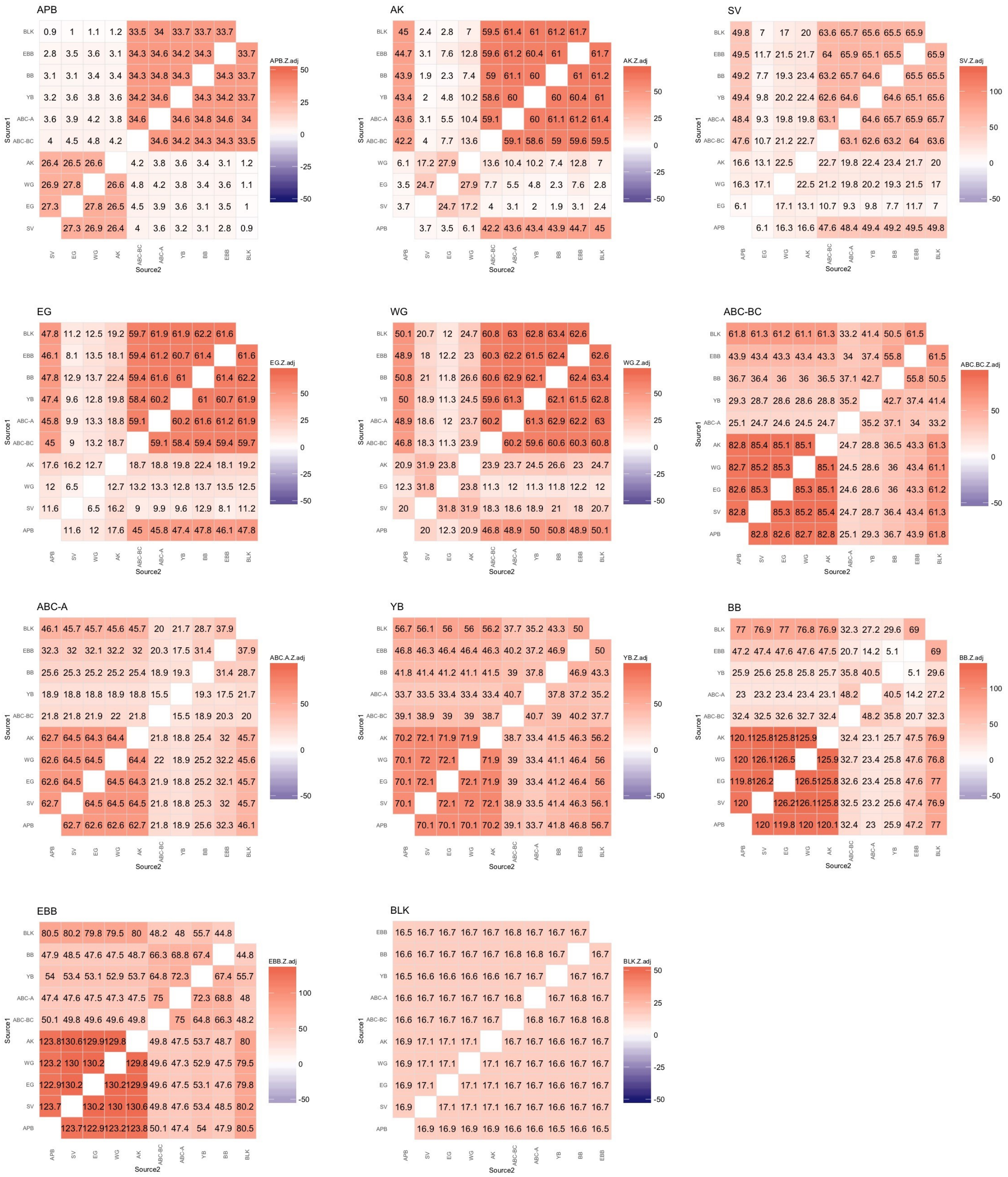

**Fig. S14.** $f_3$-statistics results (adjusted Z scores) based on population level comparisons performed using **A.** dataset DS2 (excluding private alleles) and **B.** DS6 (excluding transition sites from DS2). Population acronyms as follows: EBB = European brown bears, BB = Alaskan mainland brown bears, YB = continental brown bears, ABC-A = ABC brown bears from Admiralty Island, ABC-BC = ABC brown bears from Baranof and Chichagof Islands, WG = West Greenland polar bears, EG = East Greenland polar bears, AK = Alaska polar bears, and SV = Svalbard polar bears.

$f_3$ statistics (adj. Z scores, DS2)

Ancient polar bear (APB)

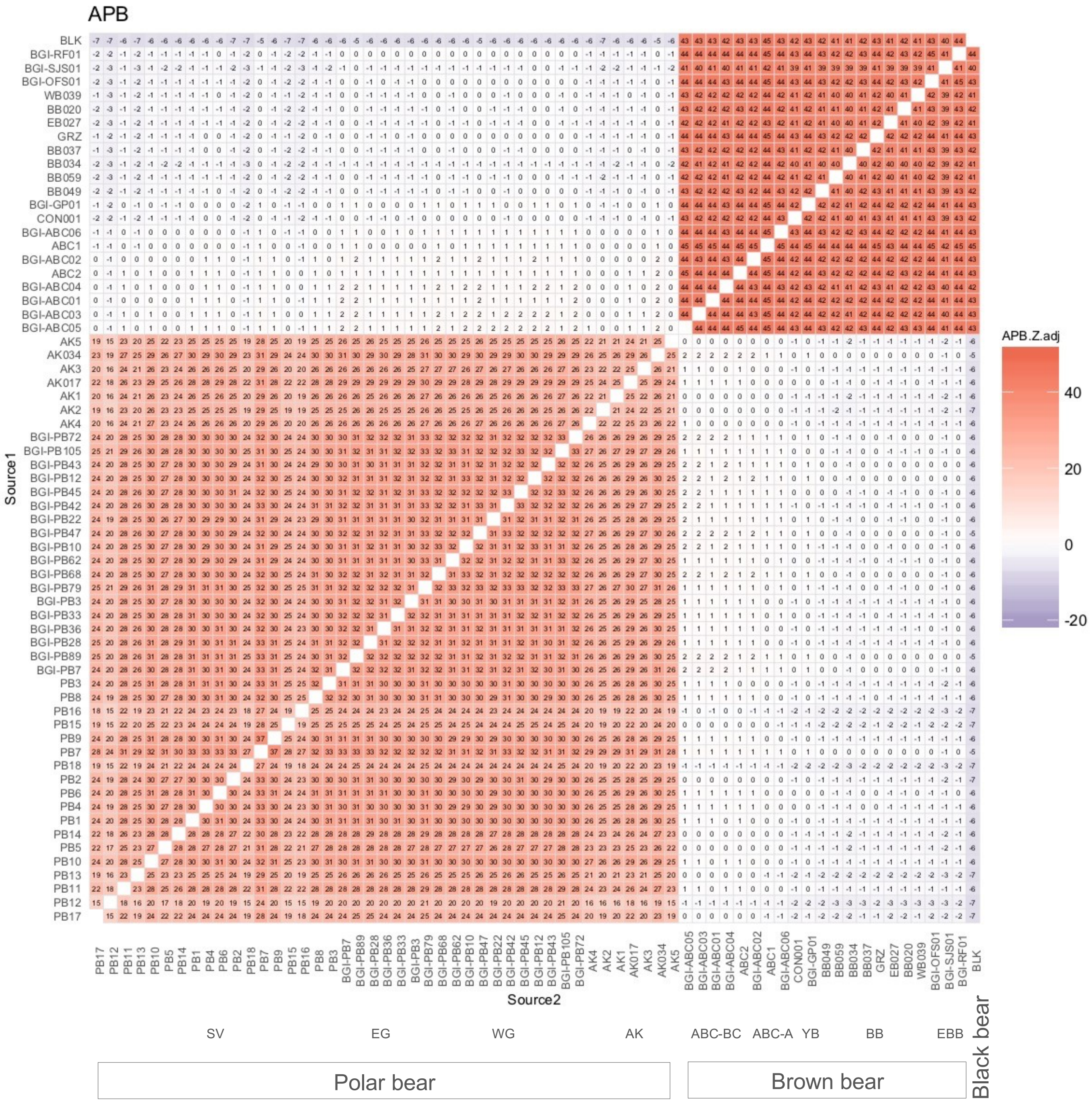

### Modern polar bear (AK)

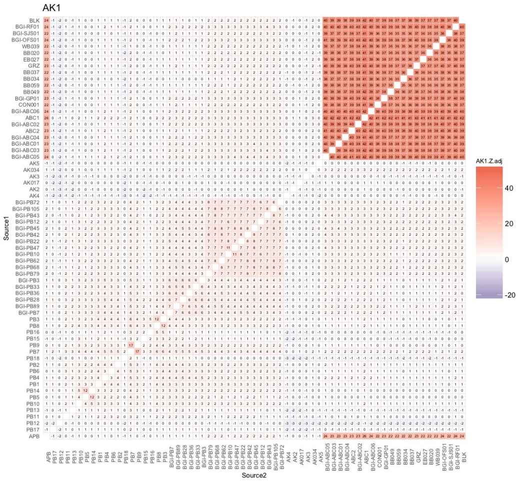


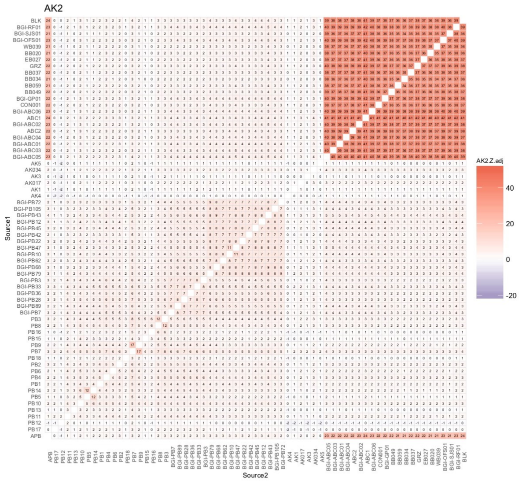


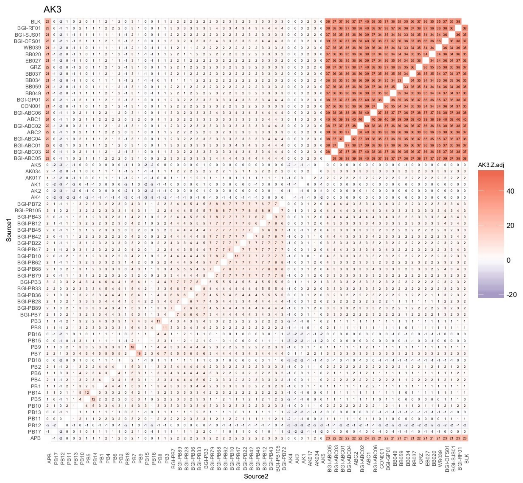


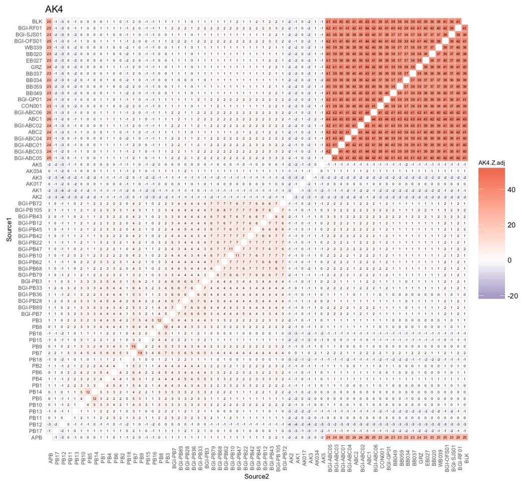


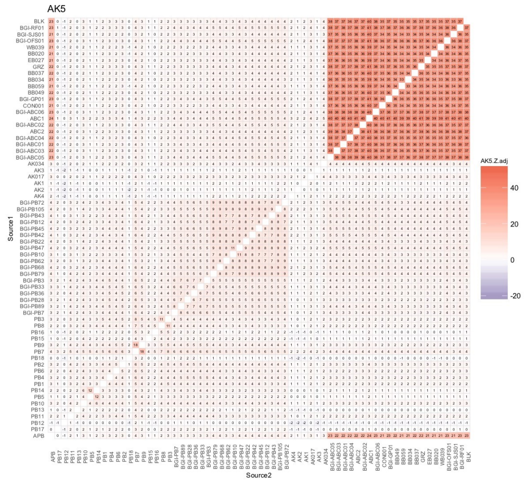


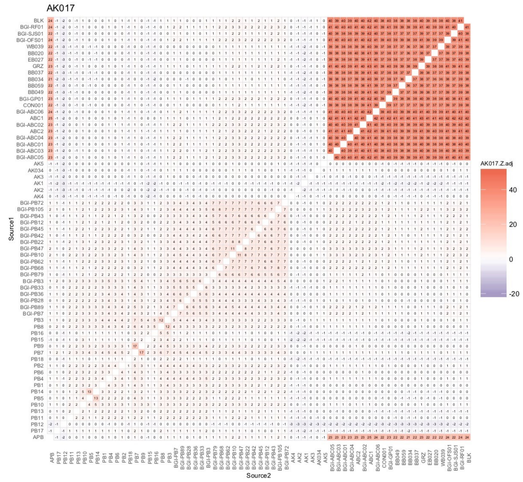

### Modern polar bear (AK, SV)

AK034

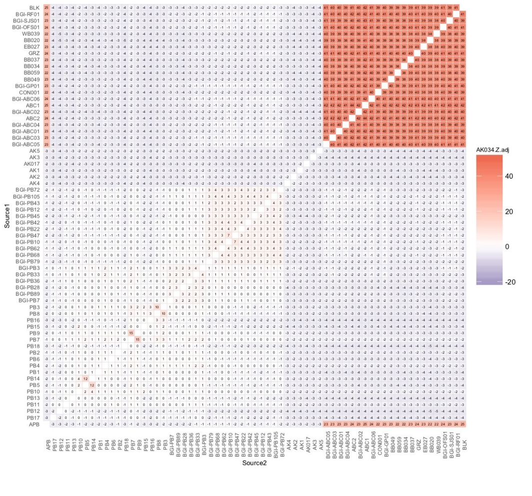

PB1

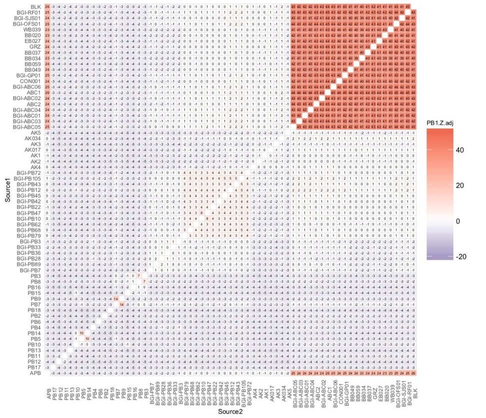

PB2

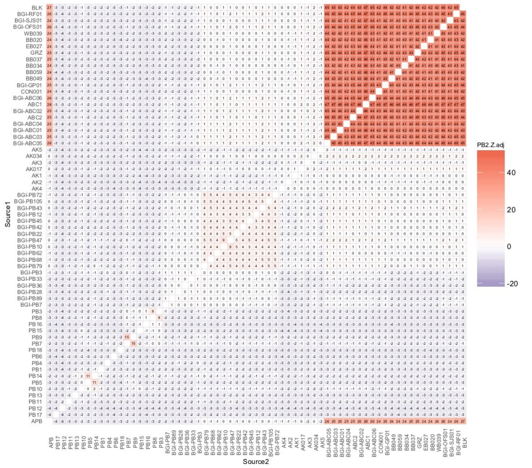

PB3

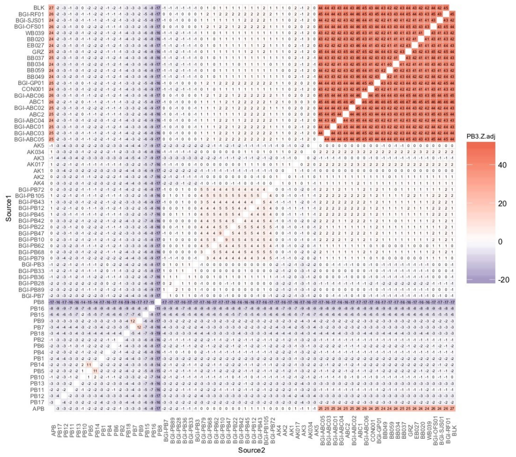

PB4

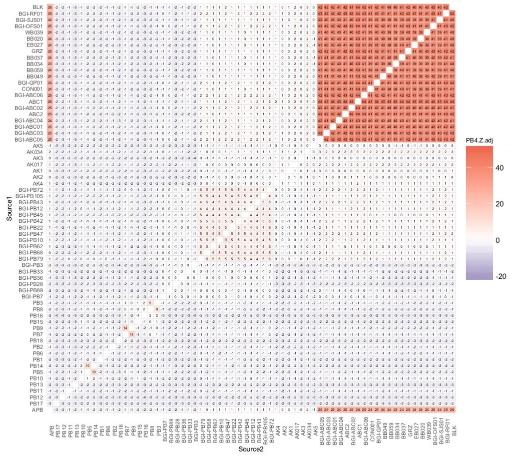

PB5

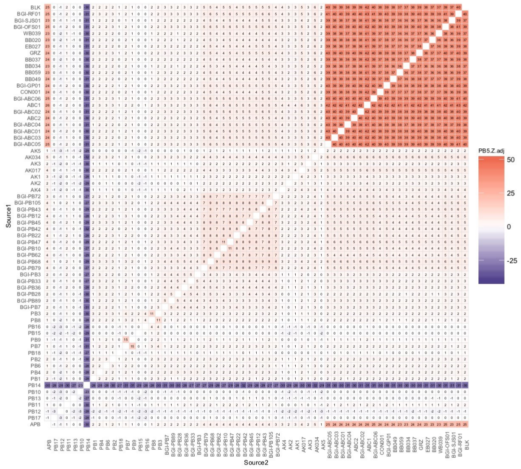

### Modern polar bear (SV)

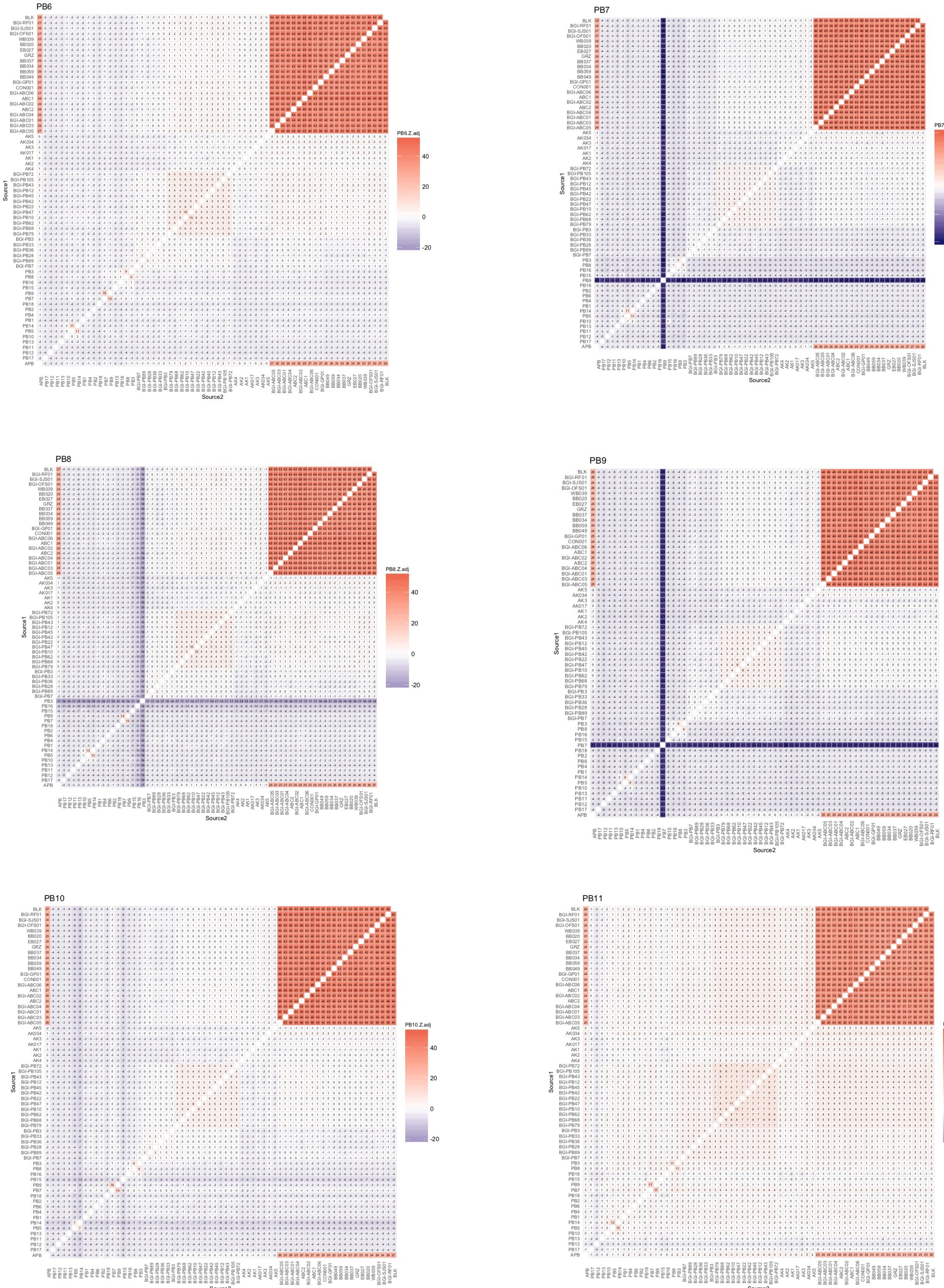

### Modern polar bear (SV)

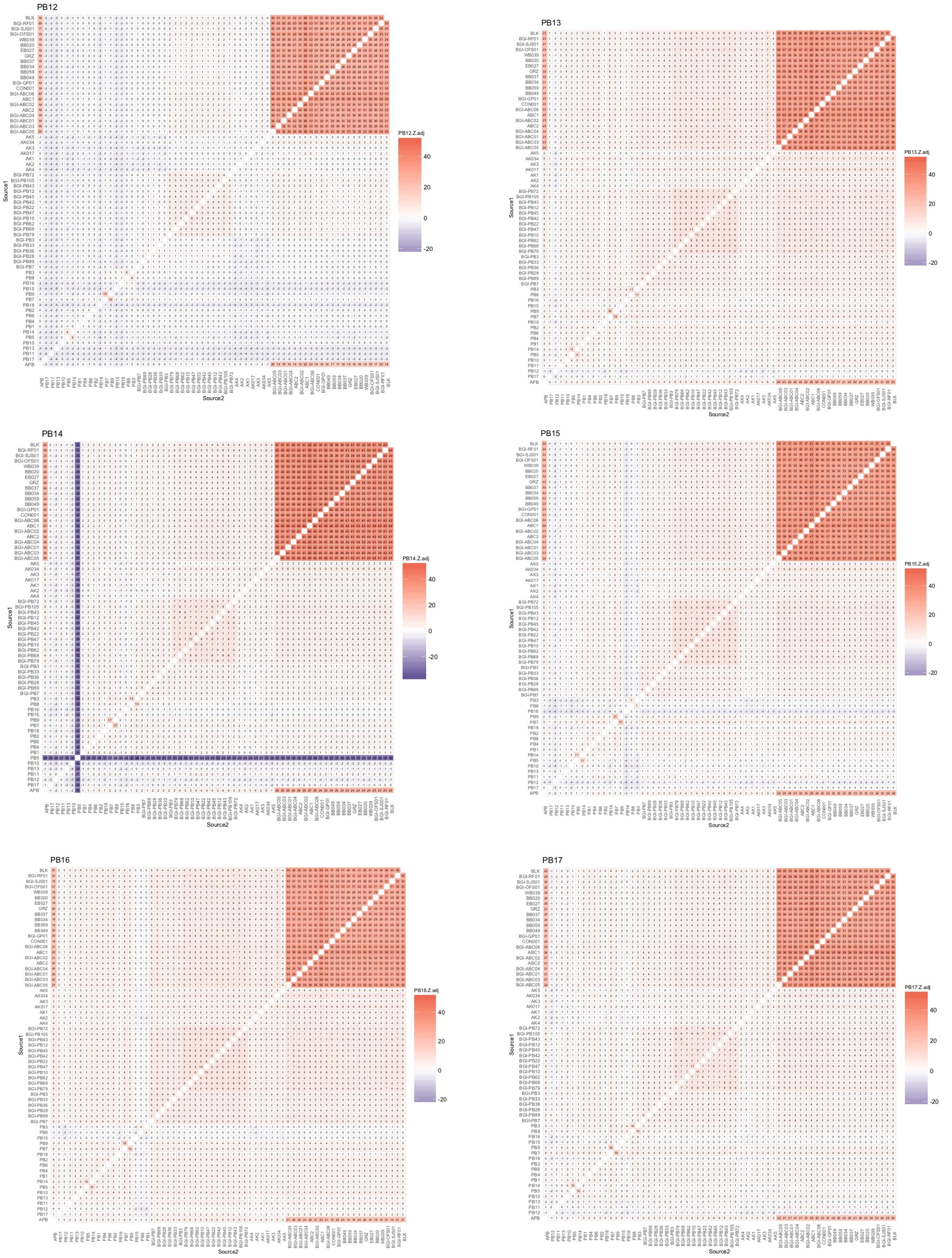

### Modern polar bear (SV, WG)

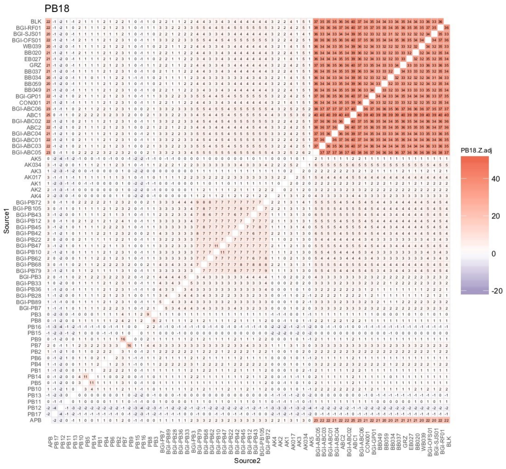


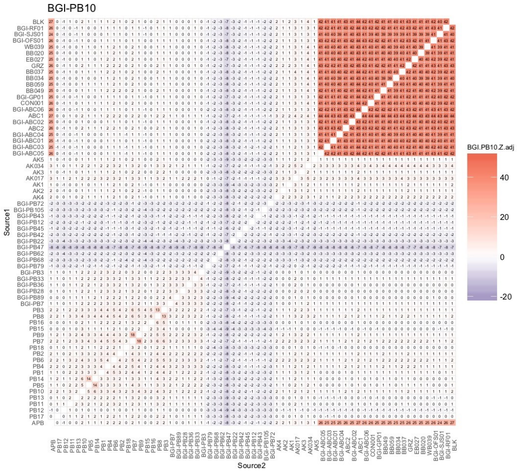


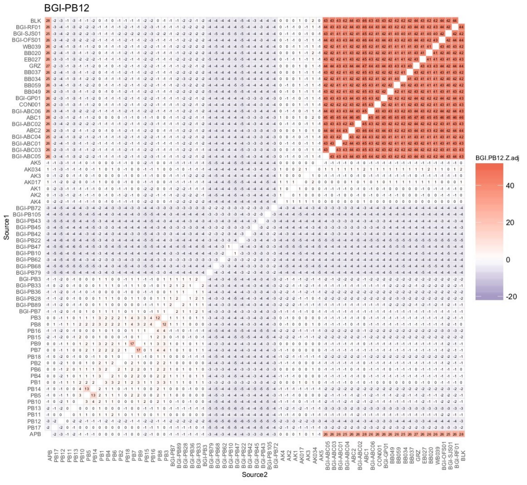


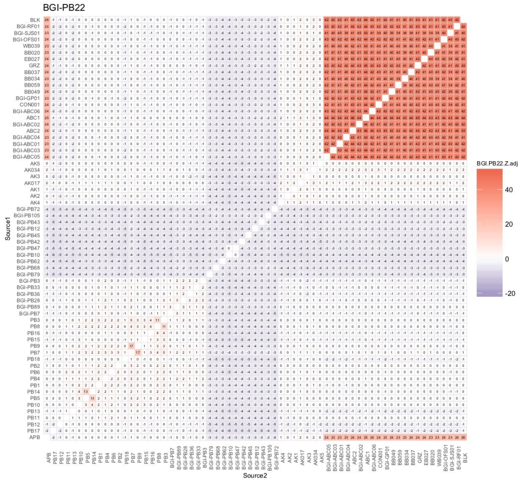


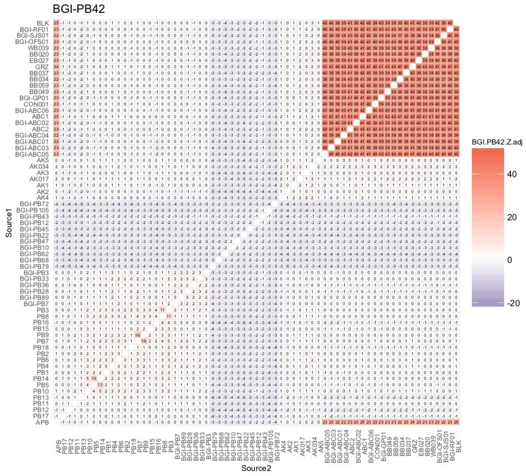


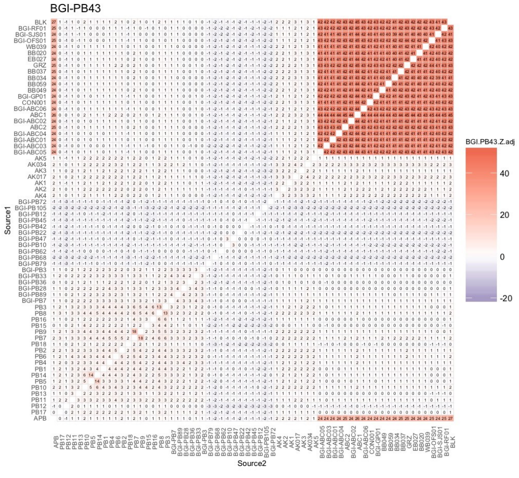

### Modern polar bear (WG)

BGI-PB45

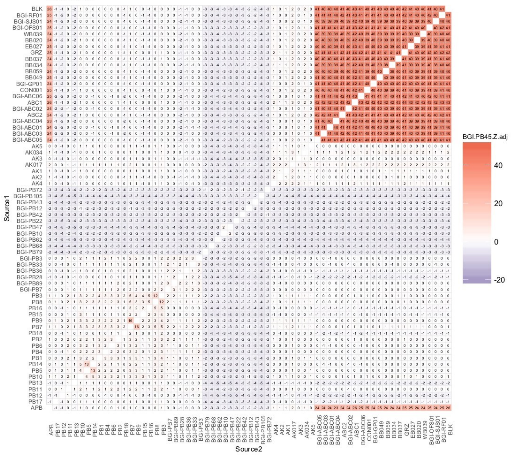


BGI-PB47

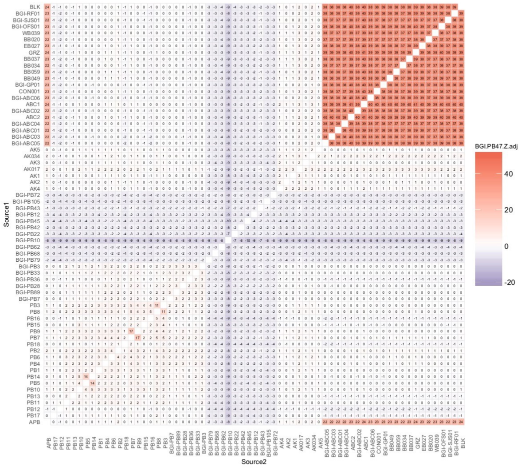


BGI-PB62

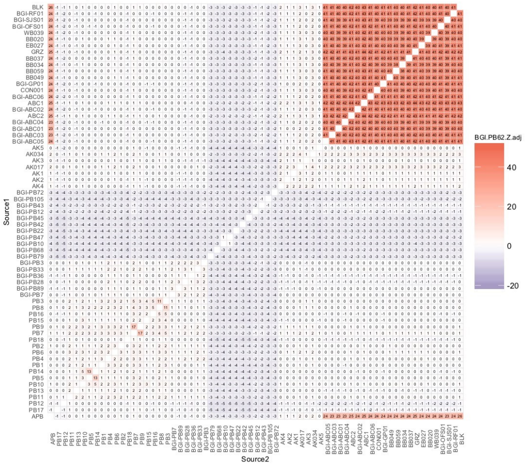


BGI-PB68

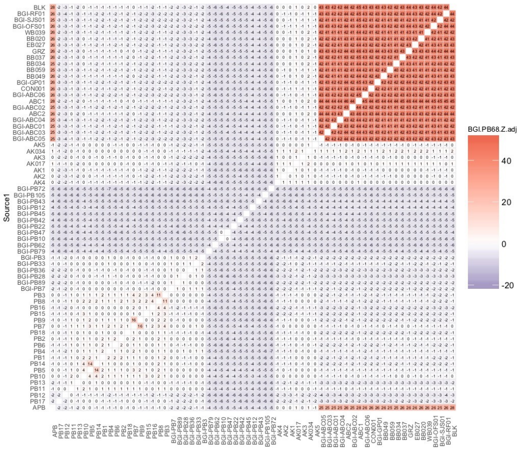


BGI-PB72

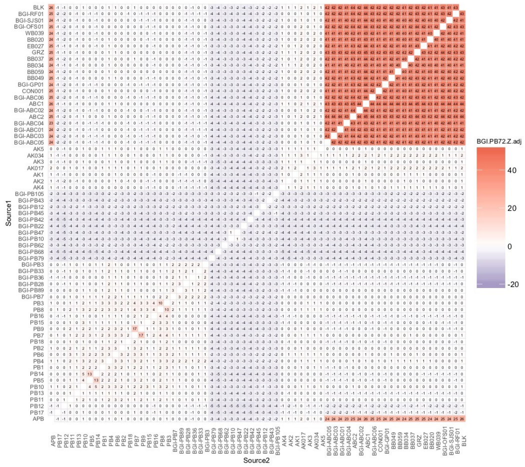


BGI-PB79

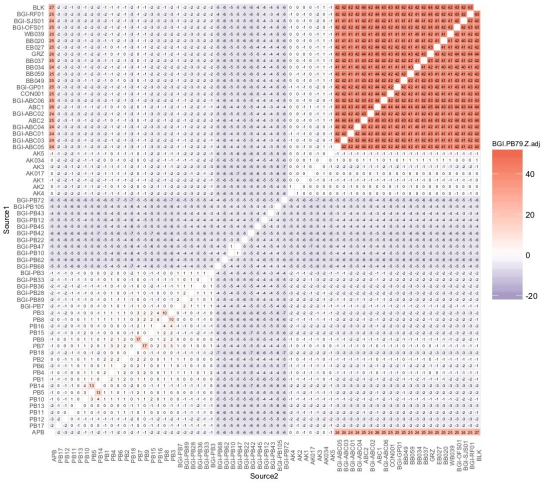

### Modern polar bear (WG, EG)

BGI-PB105

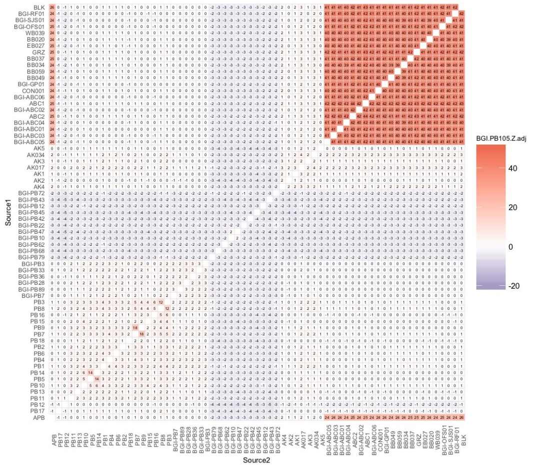


BGI-PB3

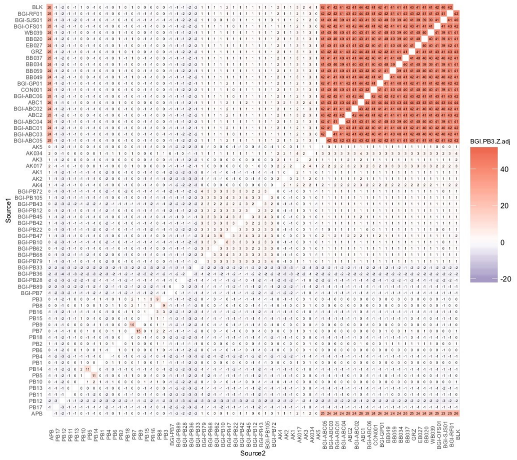


BGI-PB7

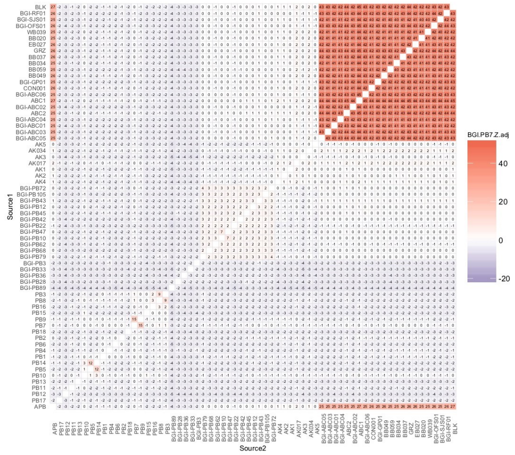


BGI-PB28

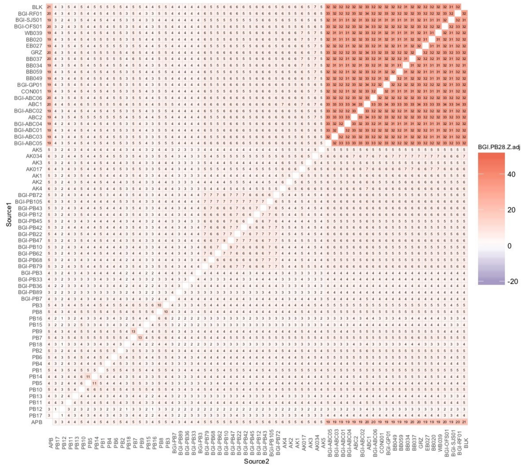


BGI-PB33

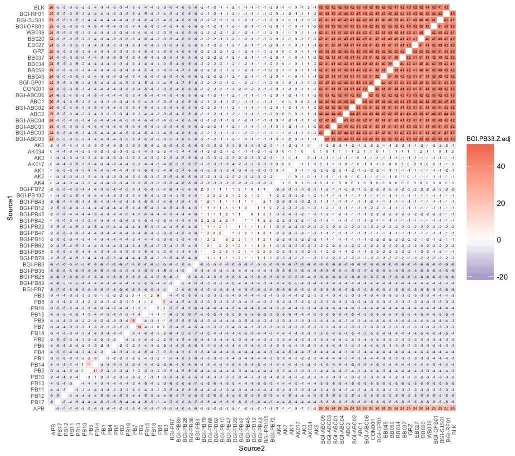


BGI-PB36

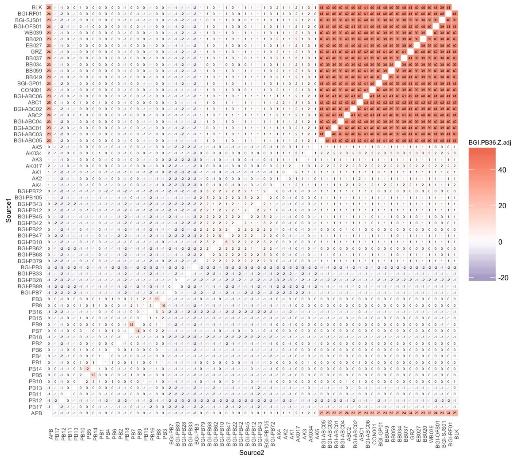

### Modern polar bear (EG), brown bear (ABC-BC)

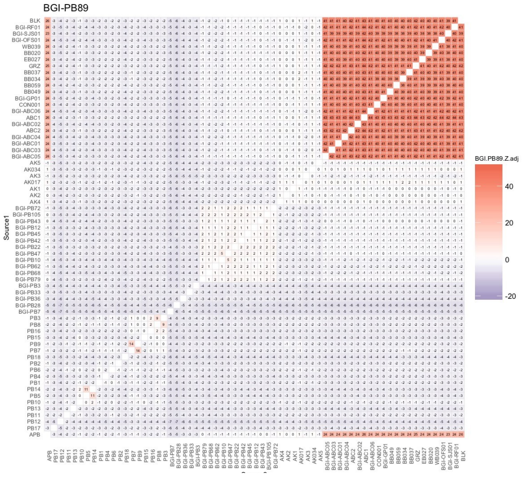


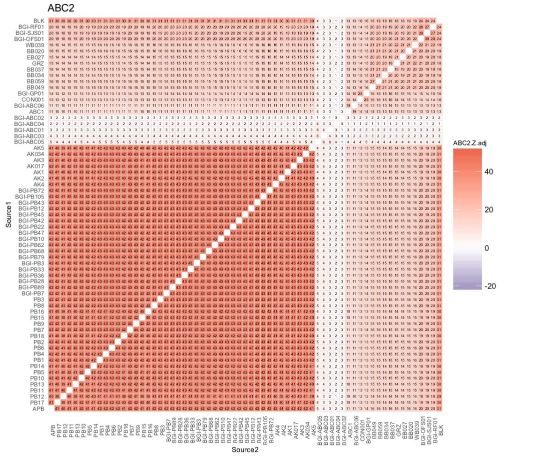


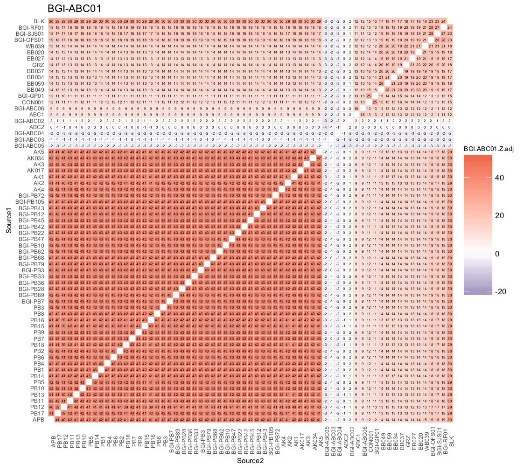


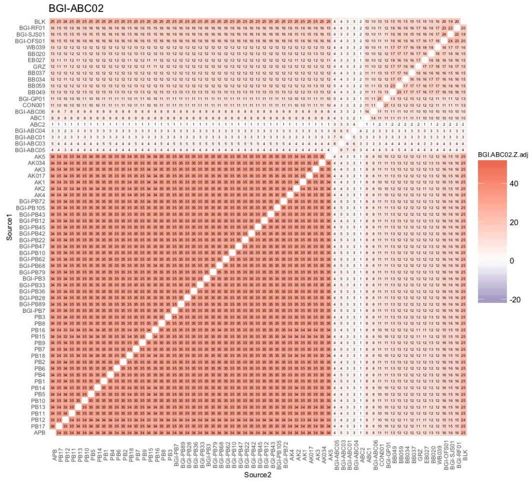


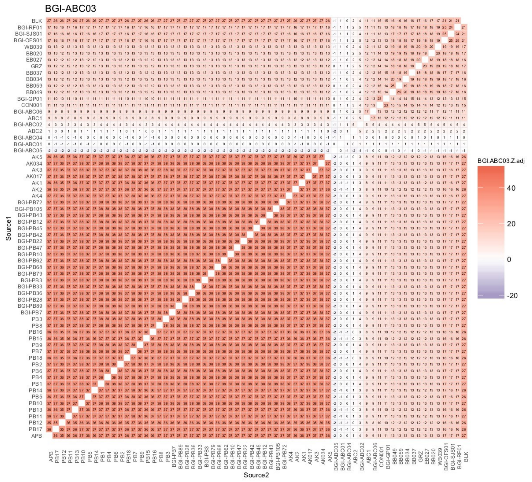


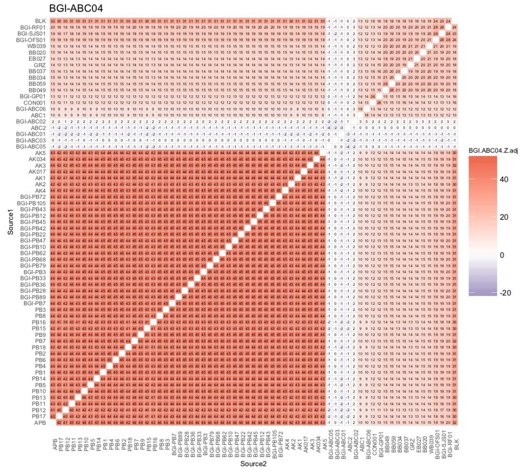

### Brown bear (ABC-BC, ABC-A, CON)

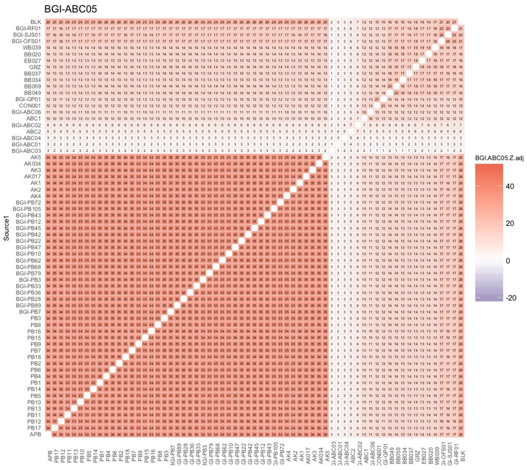


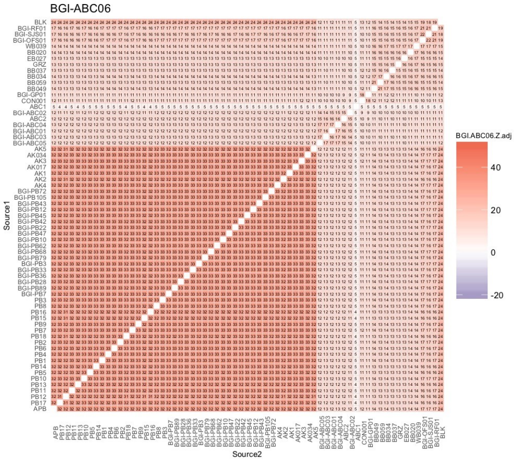


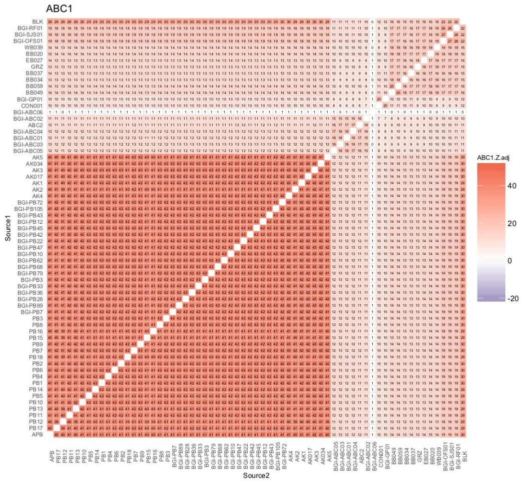


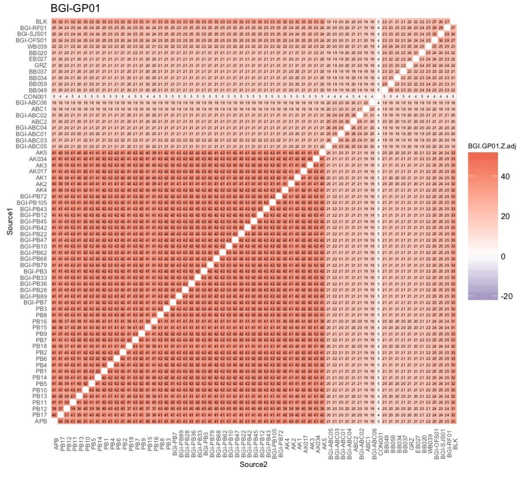


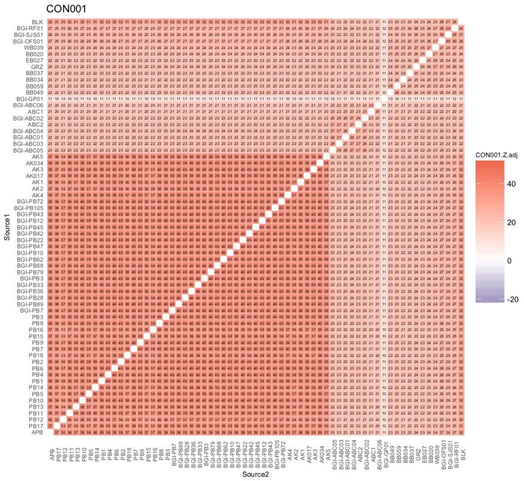

### Brown bear (BB)

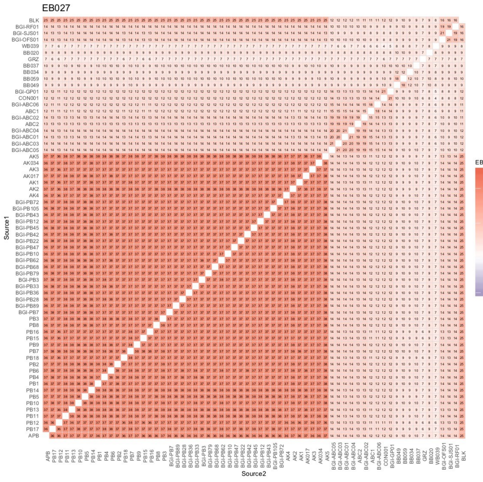


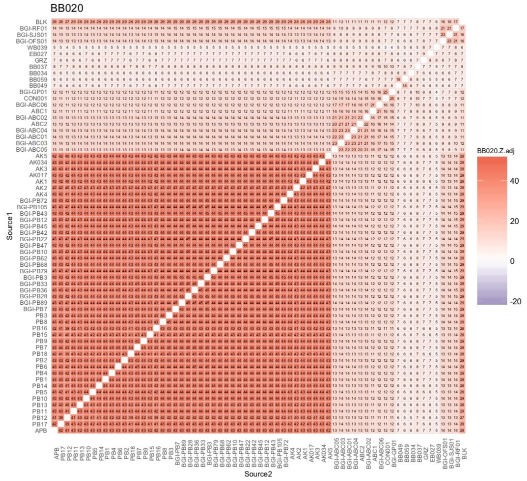


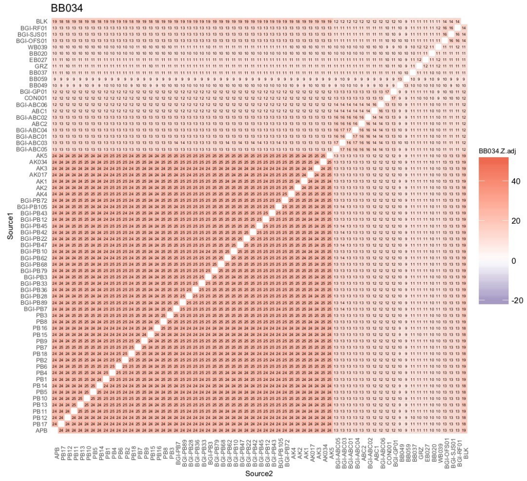


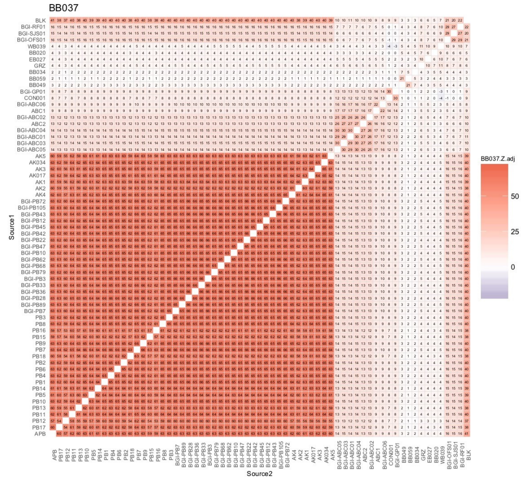


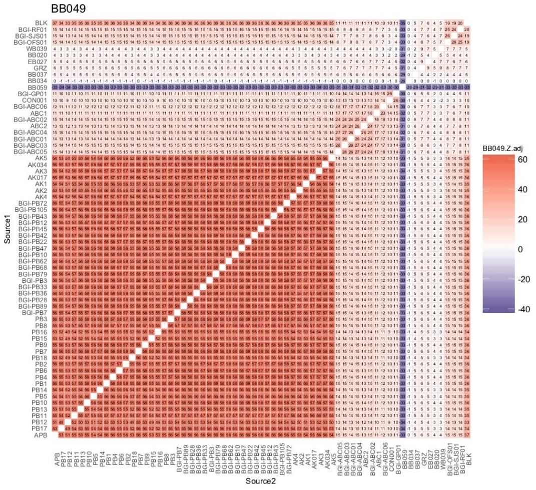


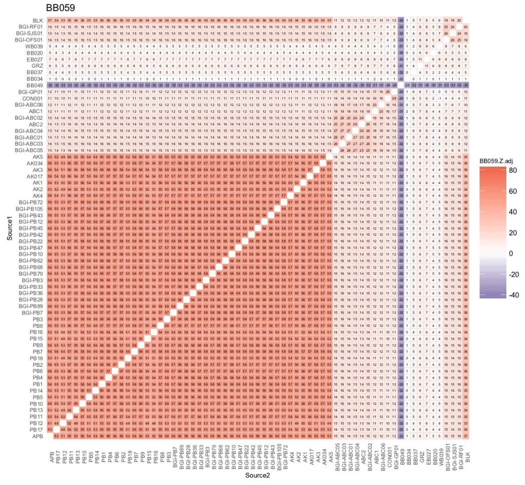

Brown bear (BB, EBB), black bear

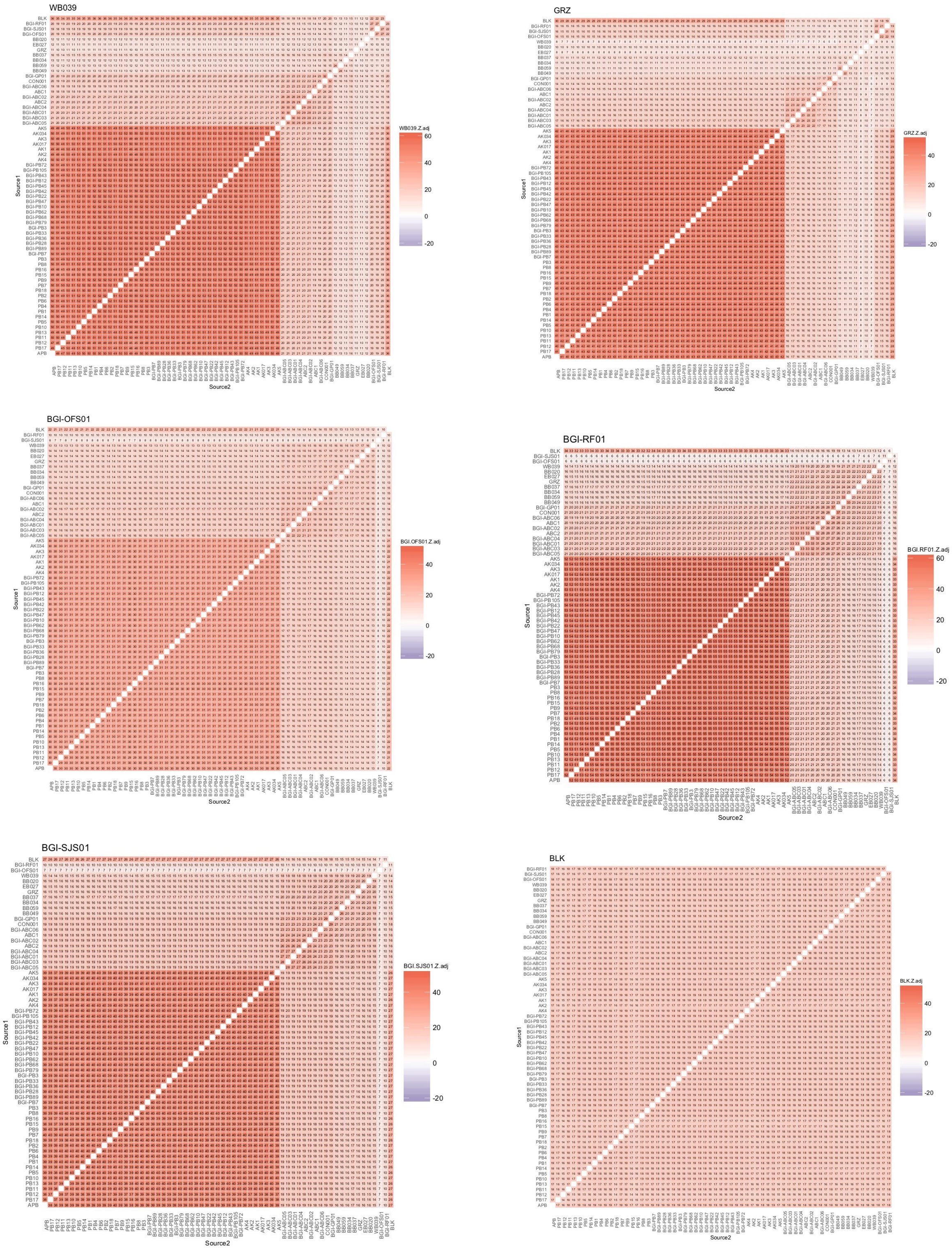


**Fig. S15.** $f_3$-statistics results (adjusted Z scores) based on individual level comparisons performed using dataset DS2 (excluding private alleles). For sample codes, see Table S1.

$f_3$ statistics (adj. Z scores, DS6)
Ancient polar bear (APB)

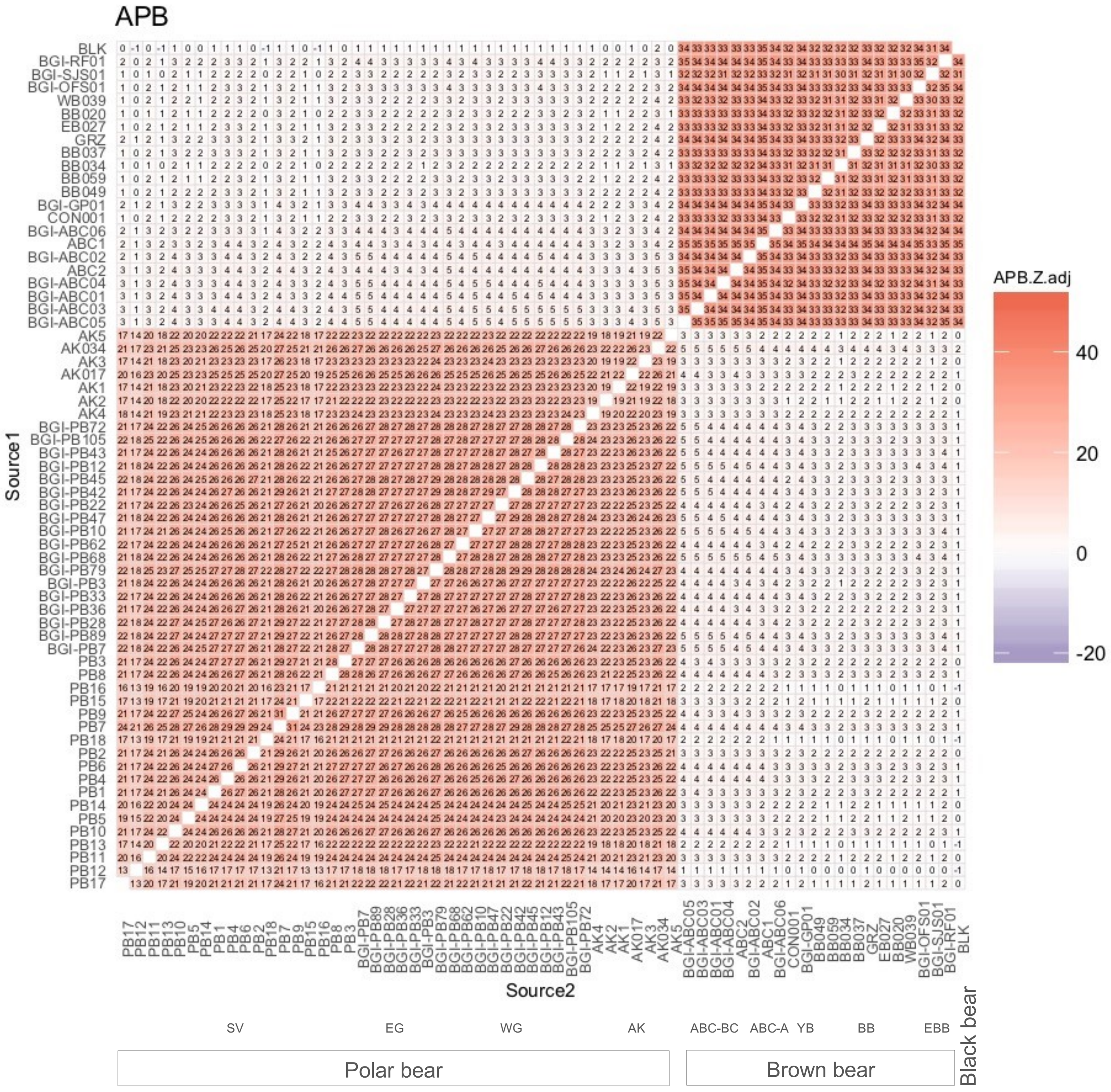


1

### Modern polar bear (AK)

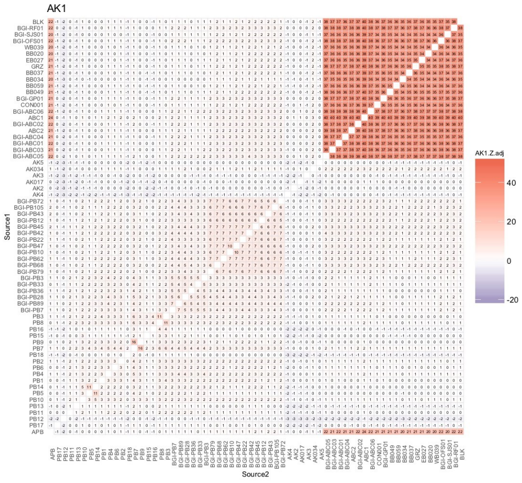


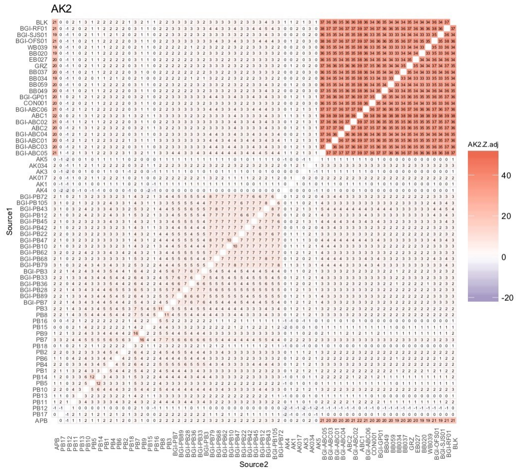


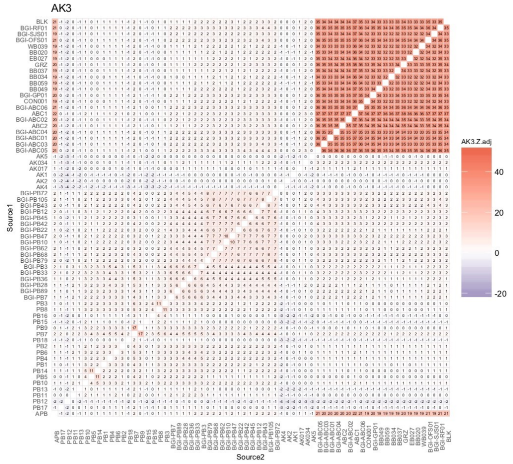


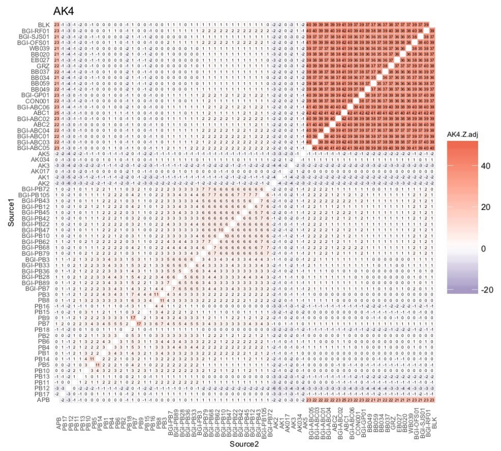


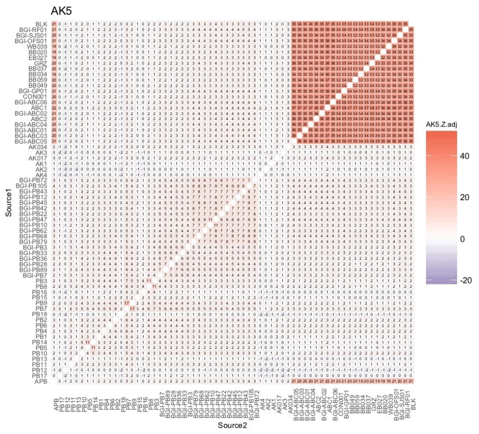


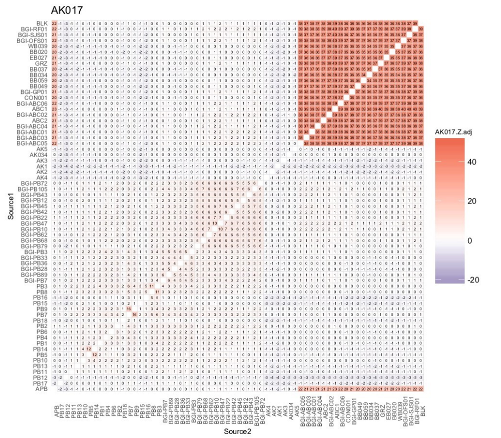

### Modern polar bear (AK, SV)

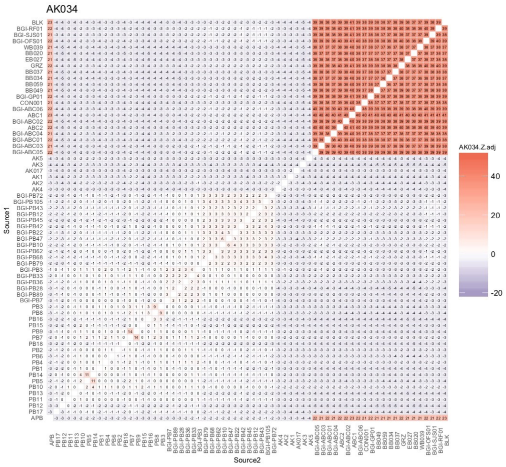


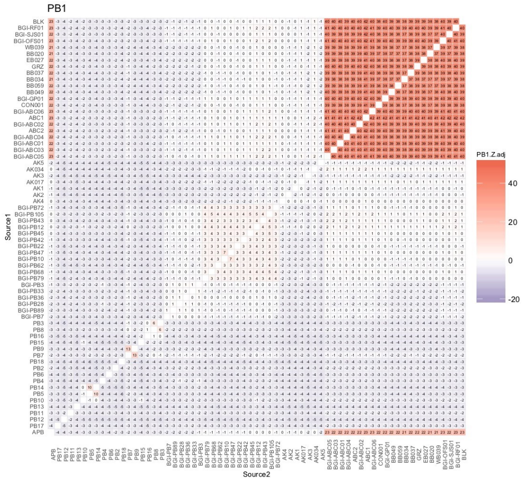


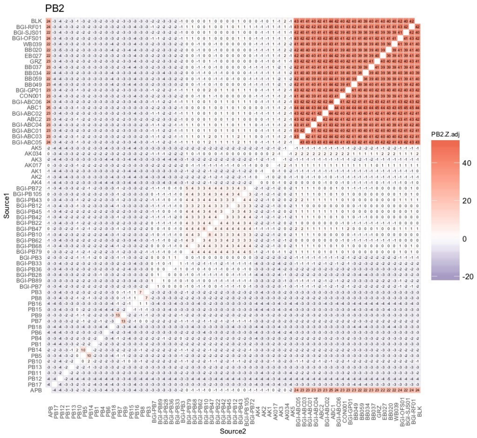


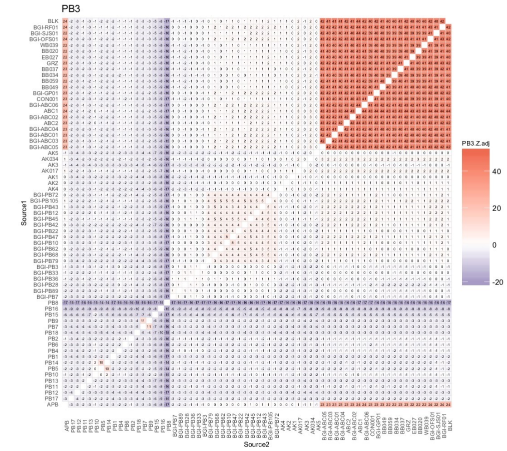


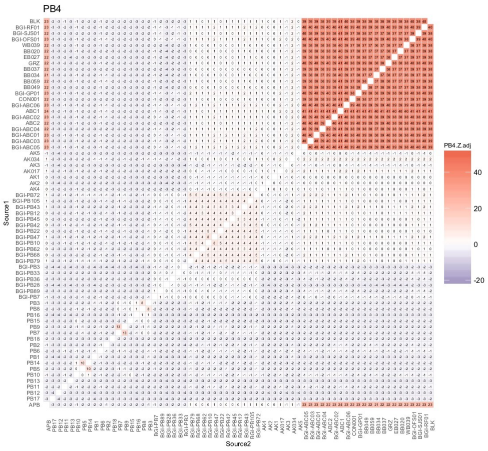


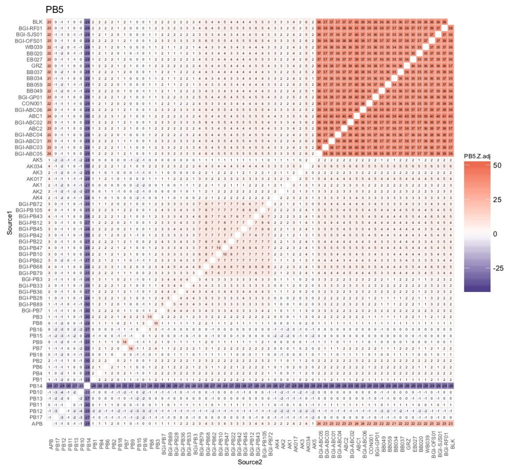

### Modern polar bear (SV)

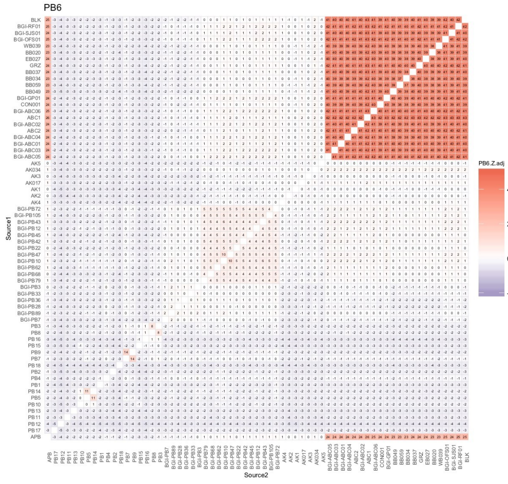


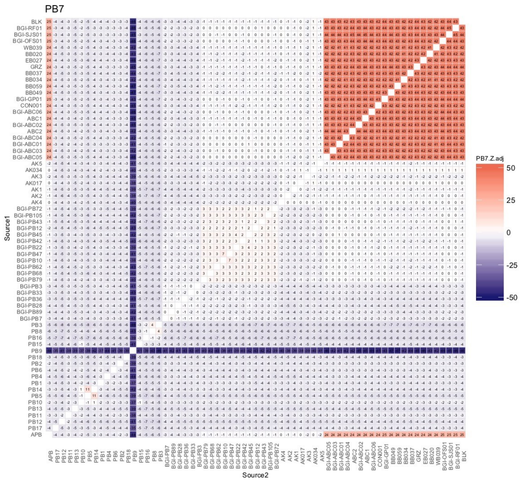


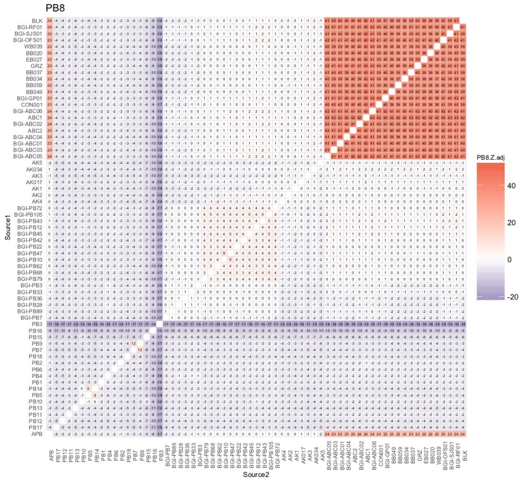


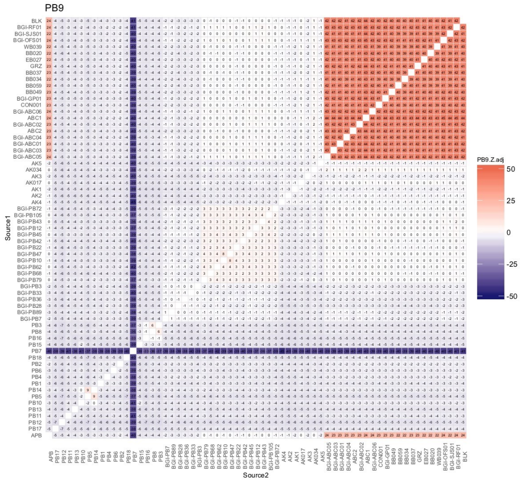


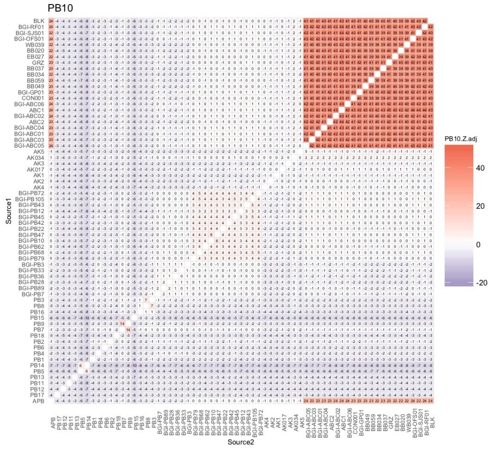


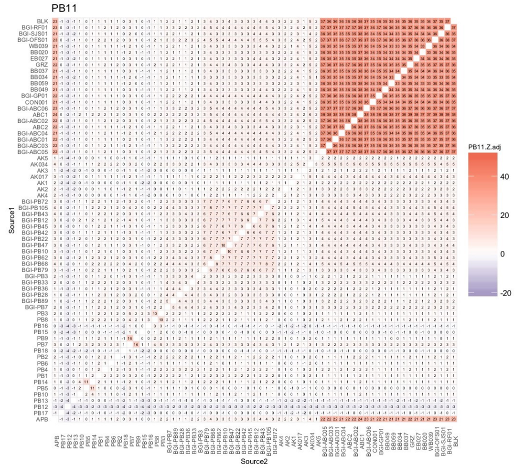

### Modern polar bear (SV)

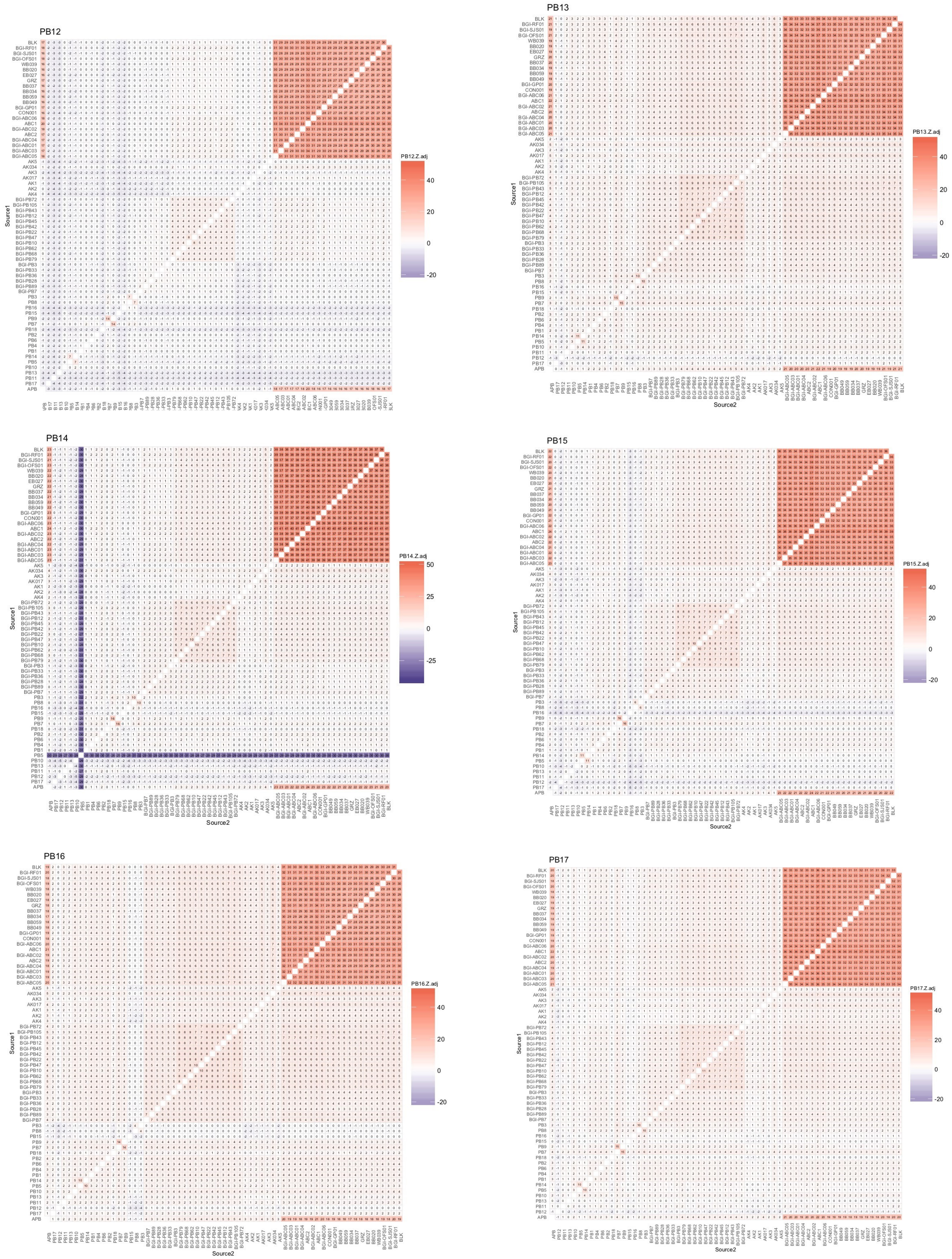

### Modern polar bear (SV, WG)

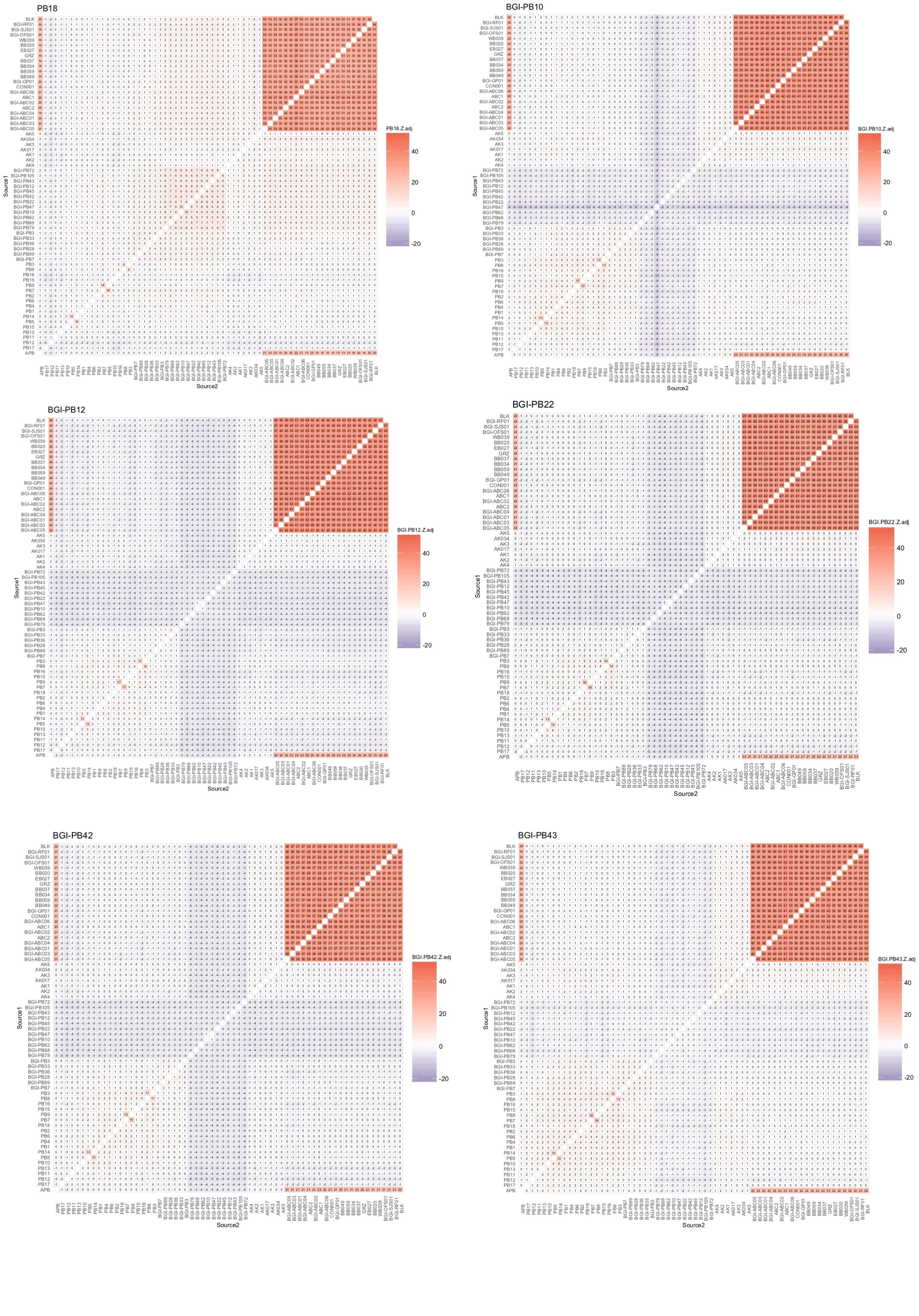

### Modern polar bear (WG)

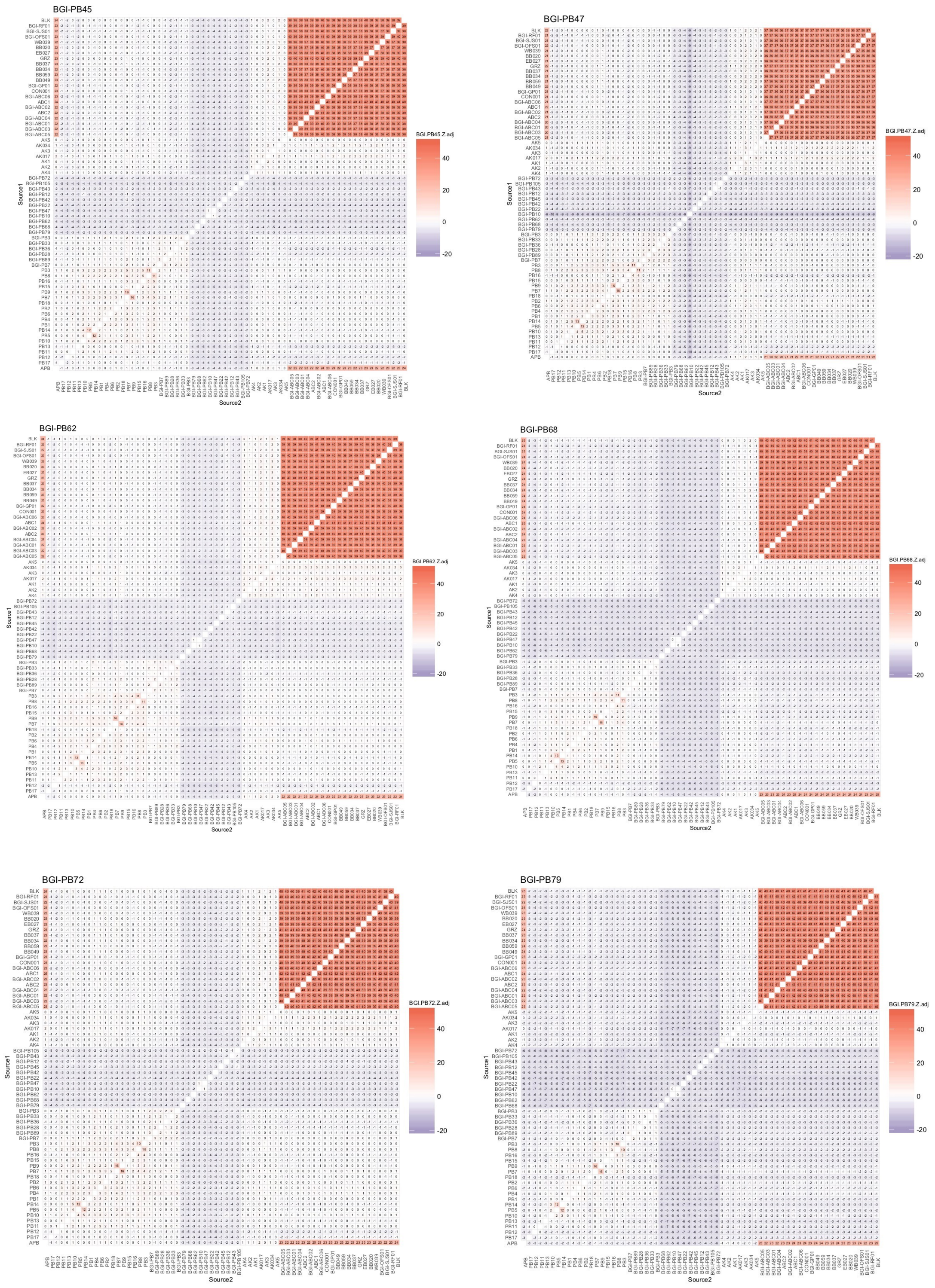

### Modern polar bear (WG, EG)

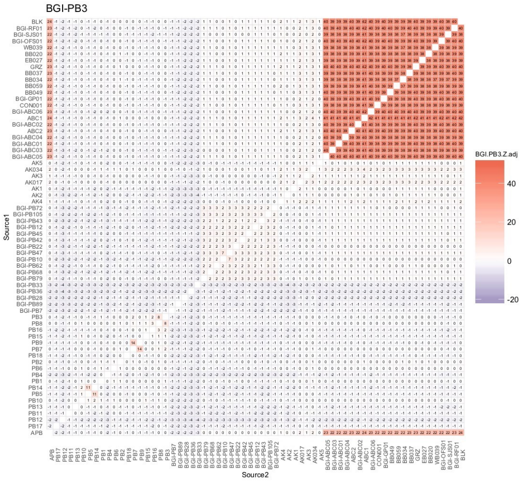


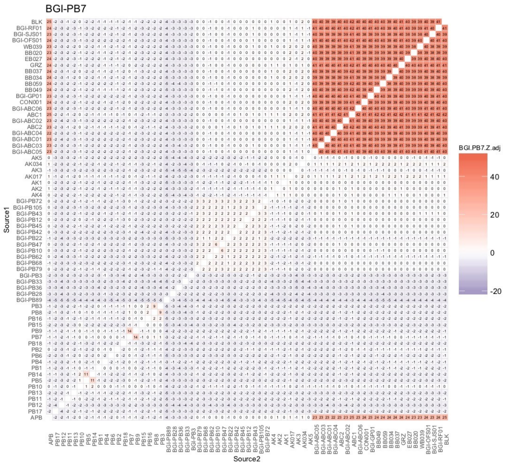


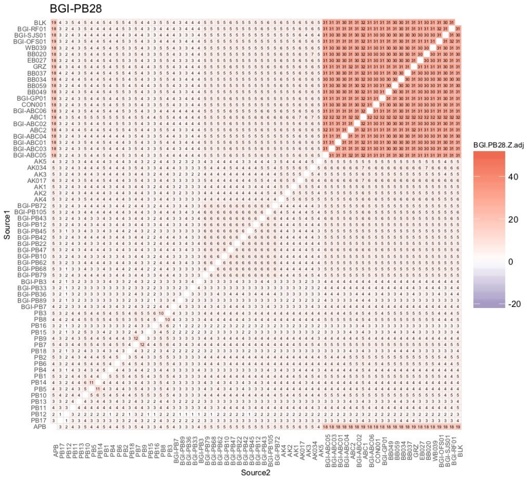


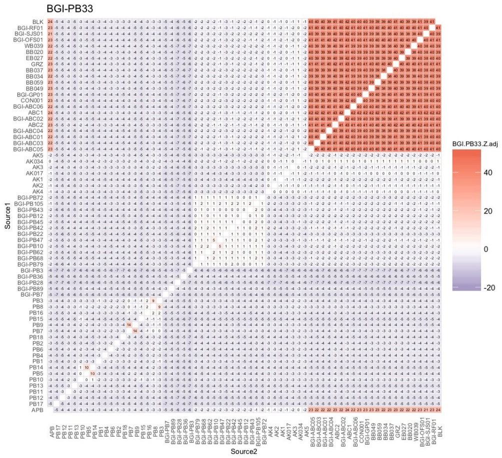


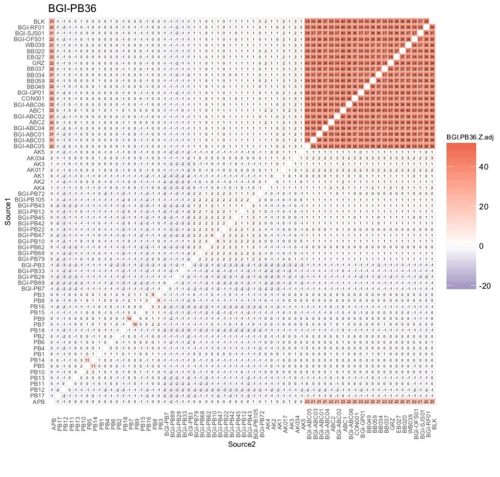


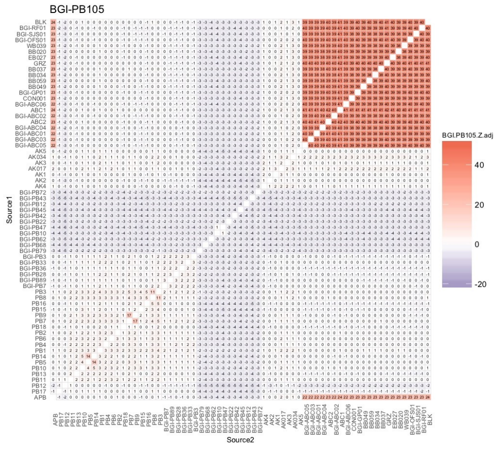

### Modern polar bear (EG), brown bear (ABC-BC)

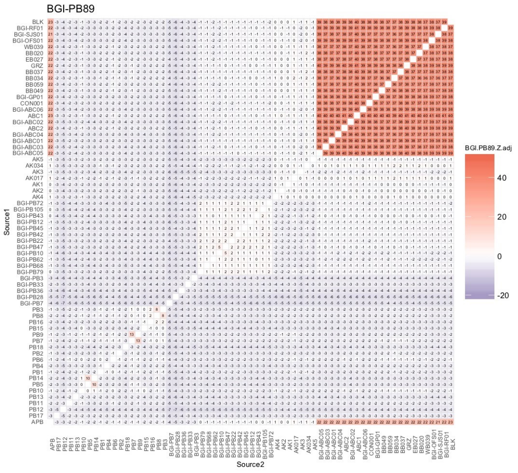


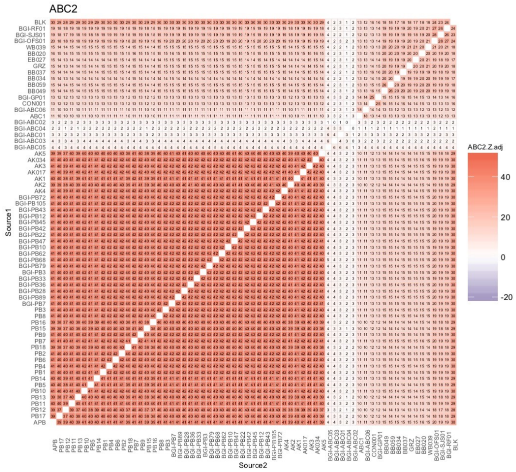


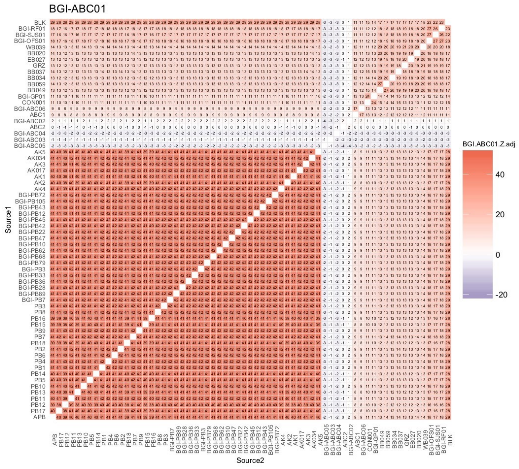


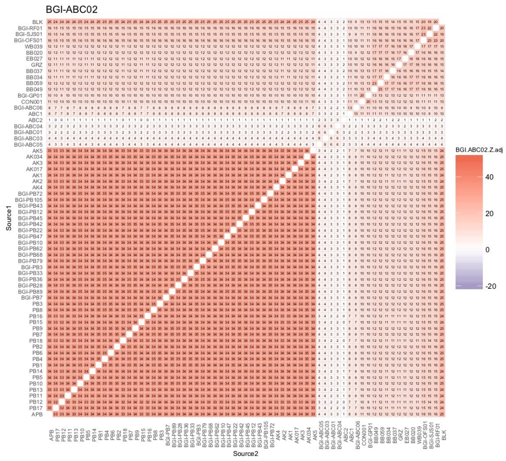


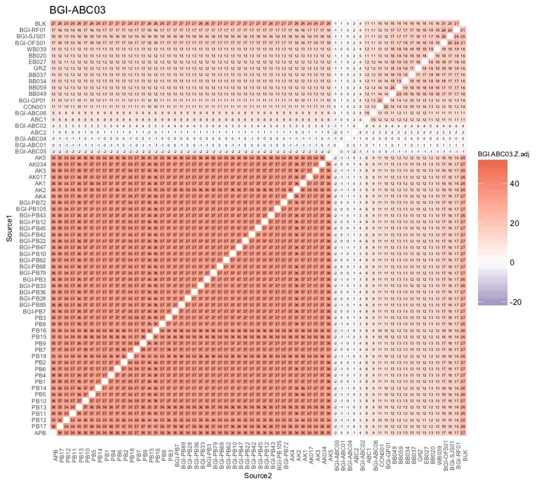


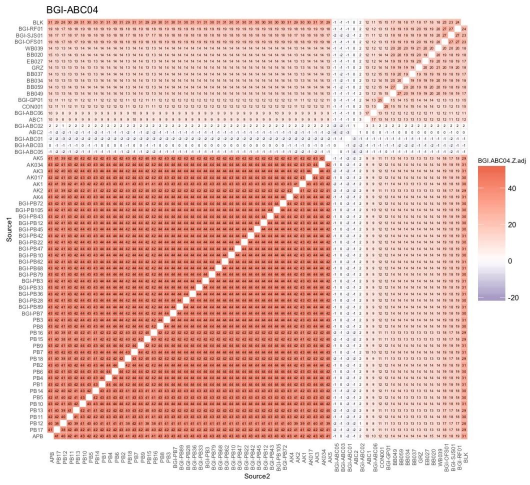

### Brown bear (ABC-BC, ABC-A, CON)

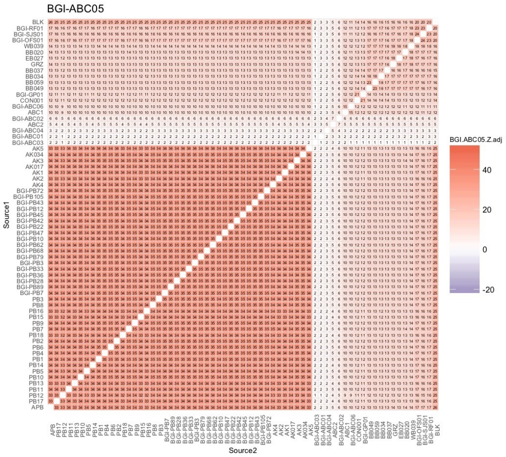


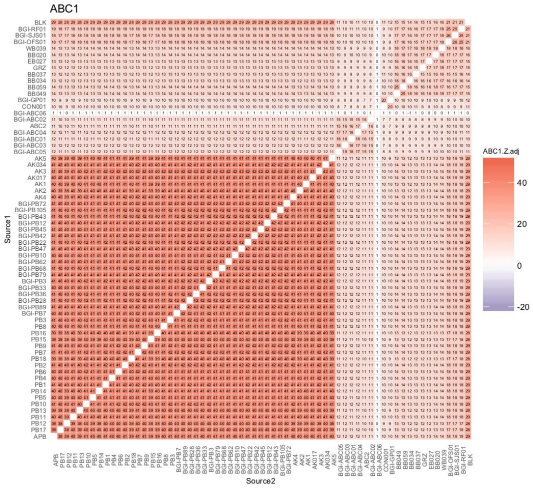


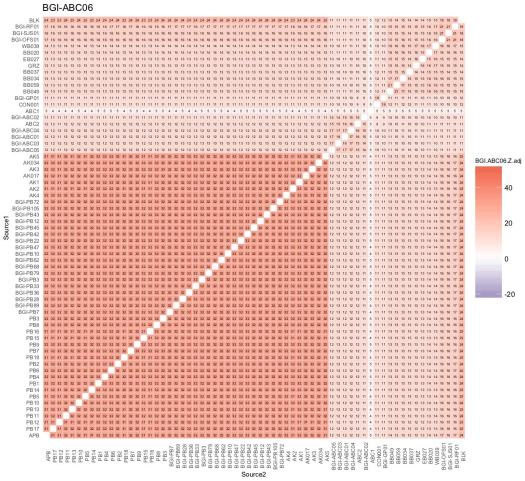


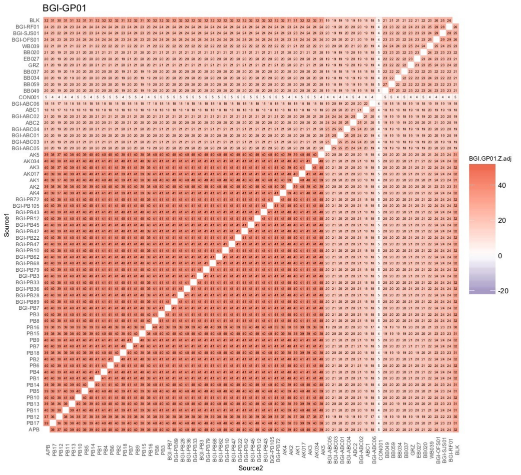


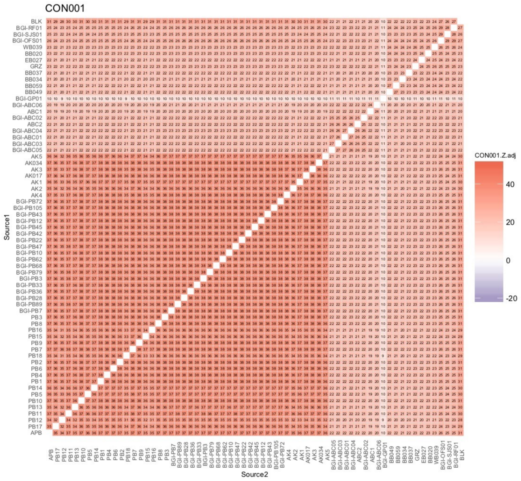

### Brown bear (BB)

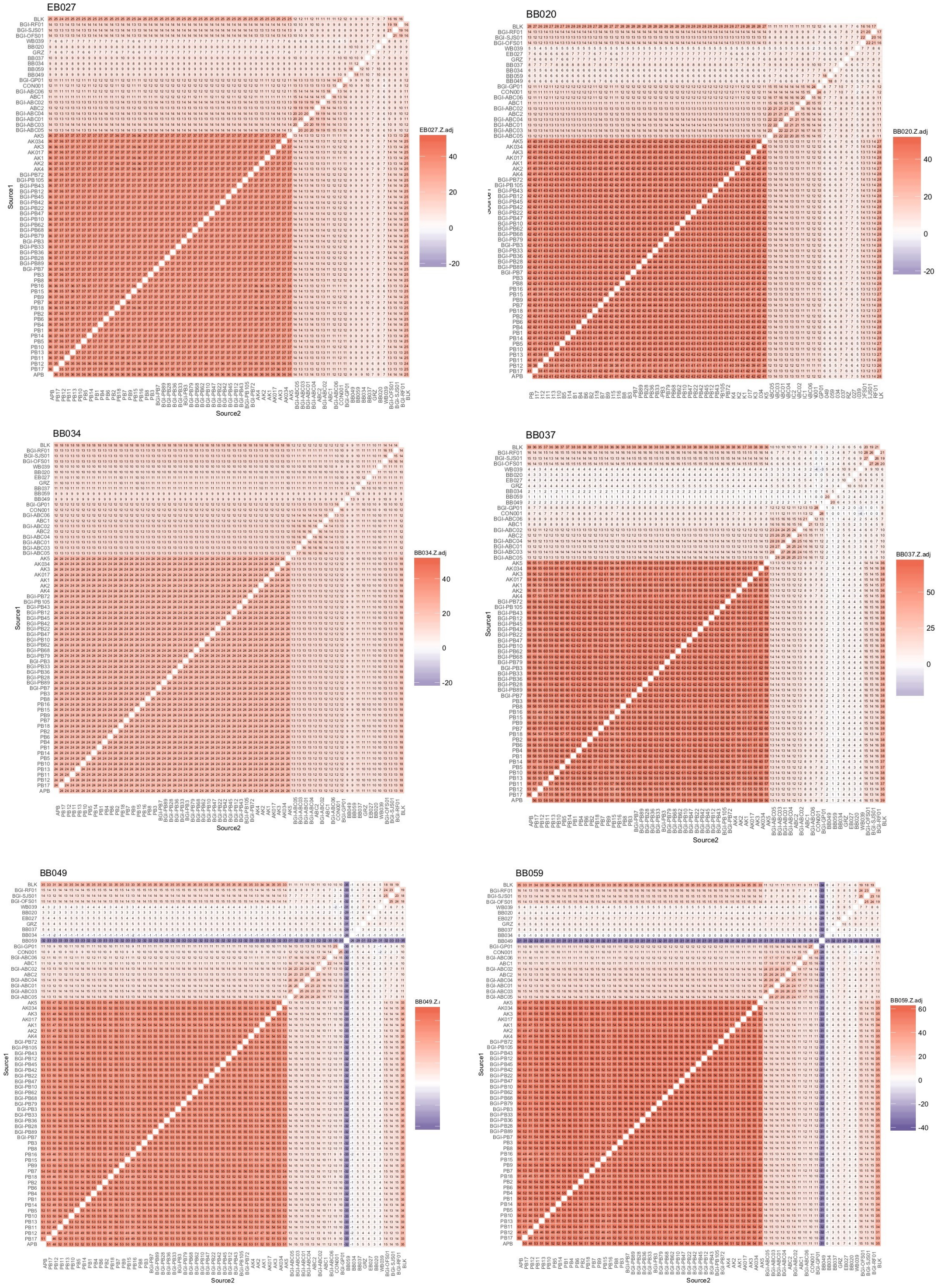

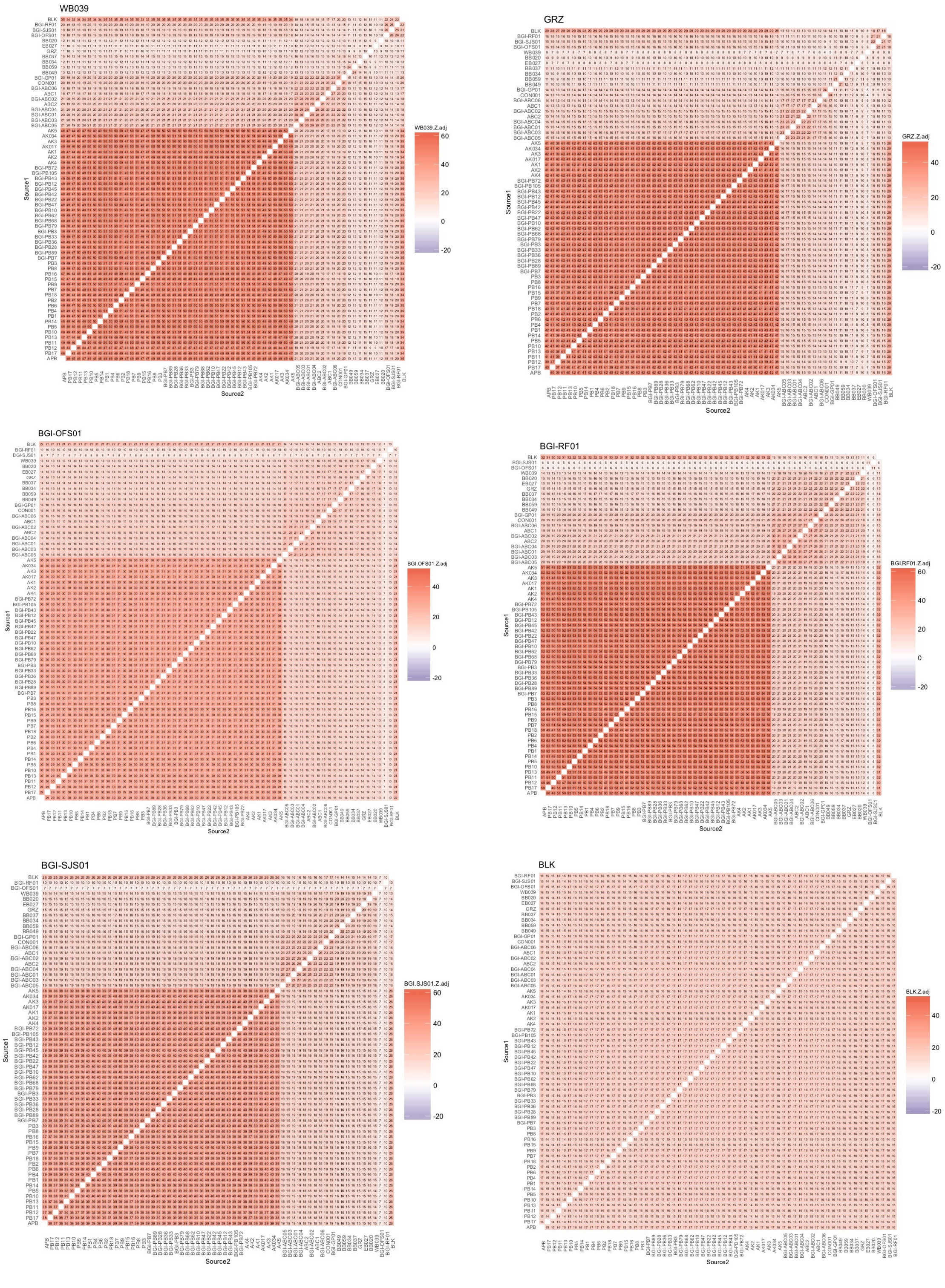

**Fig. S16.** $f_3$-statistics results (adjusted Z scores) based on individual level comparisons performed using dataset DS6 (excluding transition sites from DS2). For sample codes, see Table S1.

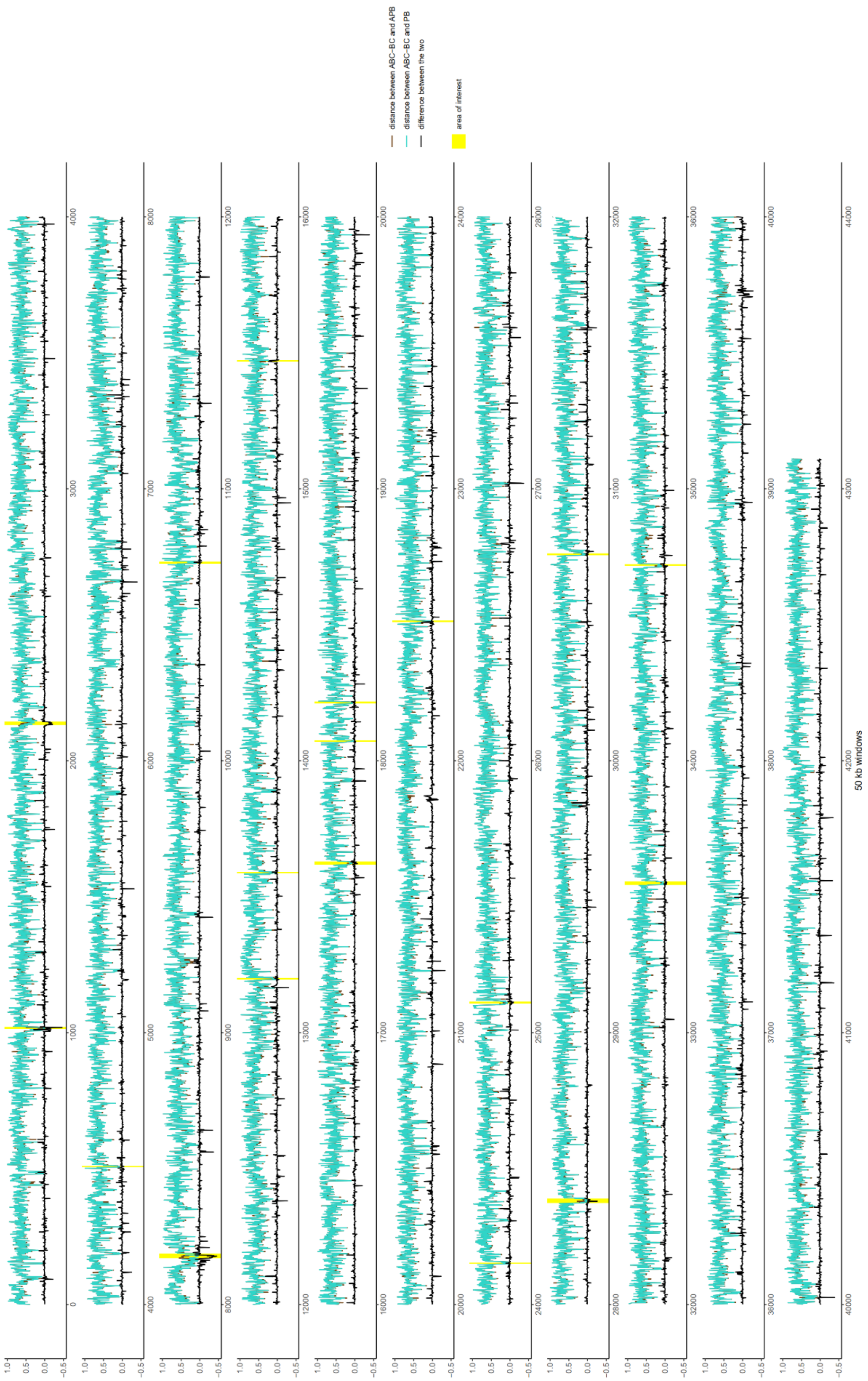


**Fig. S17.** Distance between ABC-BC and APB (brown lines) and ABC-BC and modern PB (cyan lines), and the difference between the two distances within 50 kb windows. Highlighted areas, demonstrating potentially introgressed segments, are shown in Fig. S18.

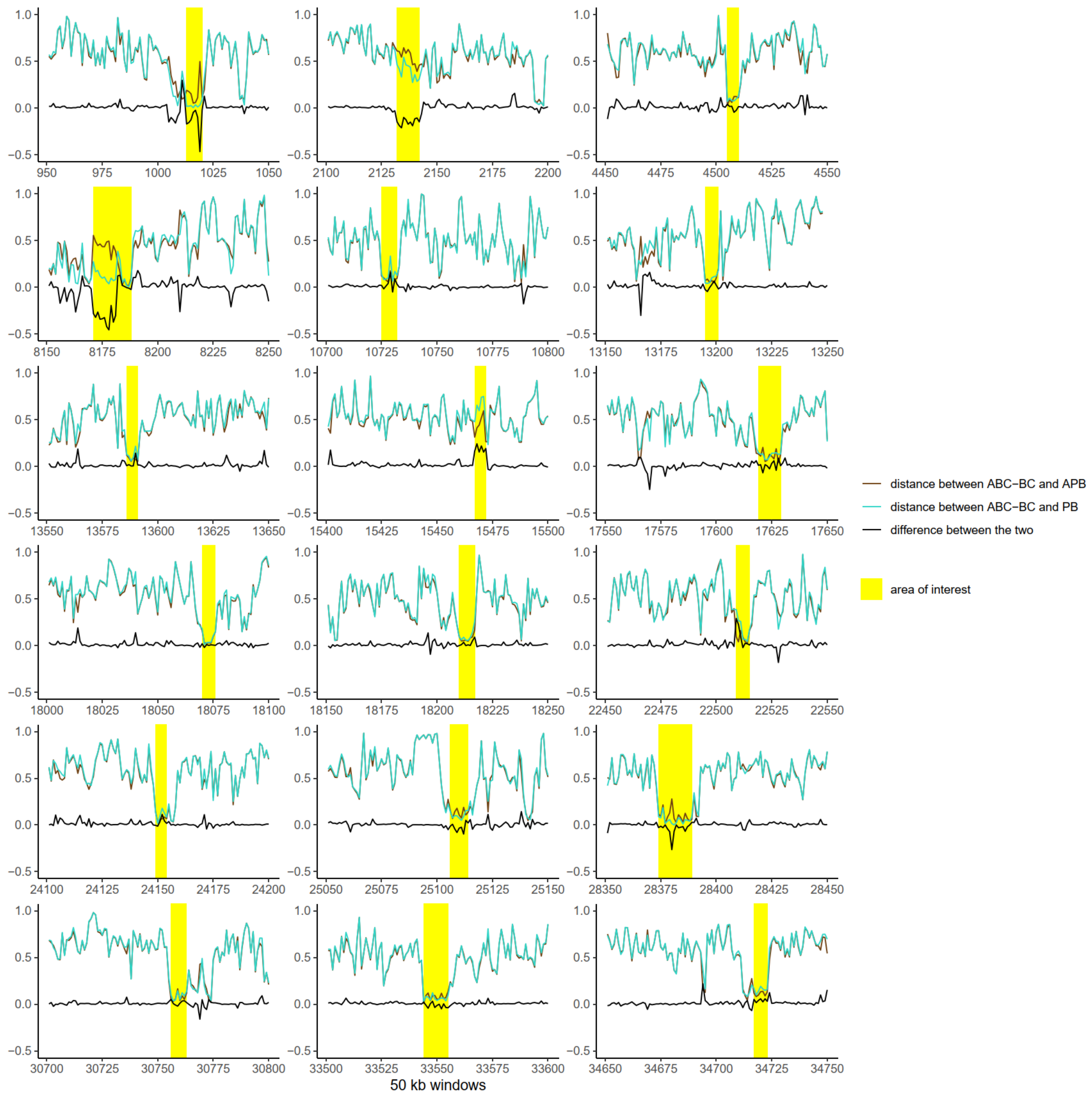


**Fig. S18.** Unusual genomic segments highlighted in Fig. S17. These potentially introgressed segments are less than 1 Mb in length, most of them only 250 kb.

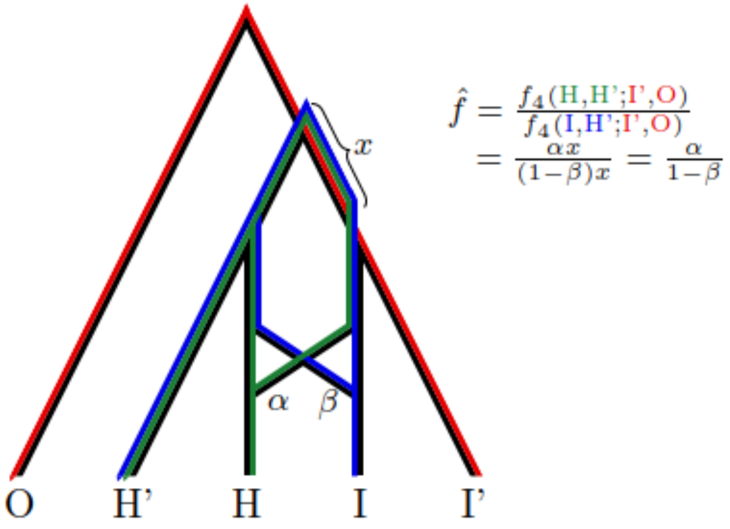


**Fig. S19.** By interpreting the $f_4$-statistics as weighted overlaps of paths in the admixture graph, we see that in the scenario depicted here $\hat{f}$ equals α /(1 - β). In particular, if the gene flow is unidirectional with β = 0, we have an estimate $\hat{f}$= α. On the same note, if the source of the gene flow into H was I' instead, the estimate would be too much, and if it was the ancestors of both I and I', it would be too small.

Case I)

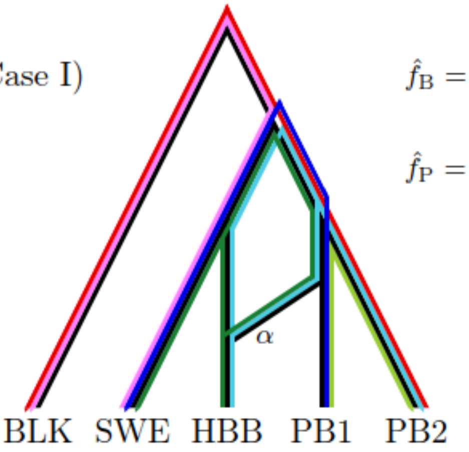


$$\hat{f}_\mathrm{B} = \frac{f_4(\mathrm{HBB},\mathrm{SWE};\mathrm{PB2},\mathrm{BLK})}{f_4(\mathrm{PB1},\mathrm{SWE};\mathrm{PB2},\mathrm{BLK})} = \alpha$$

$$\hat{f}_\mathrm{P} = \frac{f_4(\mathrm{PB1},\mathrm{PB2};\mathrm{SWE},\mathrm{BLK})}{f_4(\mathrm{HBB},\mathrm{PB2};\mathrm{SWE},\mathrm{BLK})} = 0$$

Case II)

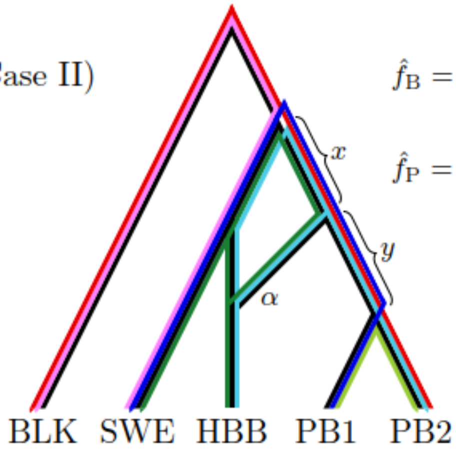


$$\hat{f}_\mathrm{B} = \frac{f_4(\mathrm{HBB},\mathrm{SWE};\mathrm{PB2},\mathrm{BLK})}{f_4(\mathrm{PB1},\mathrm{SWE};\mathrm{PB2},\mathrm{BLK})} = \frac{\alpha x}{x+y}$$

$$\hat{f}_\mathrm{P} = \frac{f_4(\mathrm{PB1},\mathrm{PB2};\mathrm{SWE},\mathrm{BLK})}{f_4(\mathrm{HBB},\mathrm{PB2};\mathrm{SWE},\mathrm{BLK})} = 0$$

Case III)

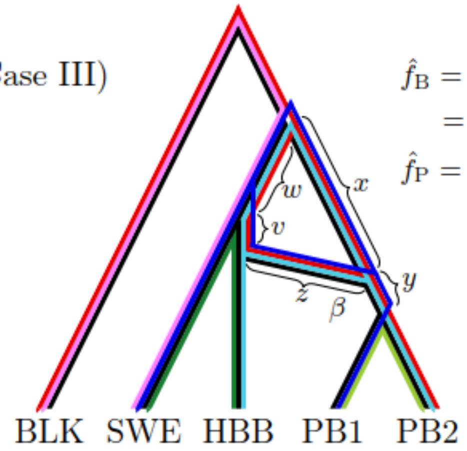


$$\hat{f}_\mathrm{B} = \frac{f_4(\mathrm{HBB},\mathrm{SWE};\mathrm{PB2},\mathrm{BLK})}{f_4(\mathrm{PB1},\mathrm{SWE};\mathrm{PB2},\mathrm{BLK})} = \frac{\beta v}{\beta^2(v+z)-\beta(1-\beta)w+(1-\beta)^2x+y}$$

$$\hat{f}_\mathrm{P} = \frac{f_4(\mathrm{PB1},\mathrm{PB2};\mathrm{SWE},\mathrm{BLK})}{f_4(\mathrm{HBB},\mathrm{PB2};\mathrm{SWE},\mathrm{BLK})} = 0$$

**Fig. S20.** Three alternative ways to explain the observed fractions $\hat{f}_\mathrm{B} \gg 0$, $\hat{f}_\mathrm{P} \approx 0$ using one unidirectional gene flow event. The direction of gene flow is ambiguous based on the two statistics only.

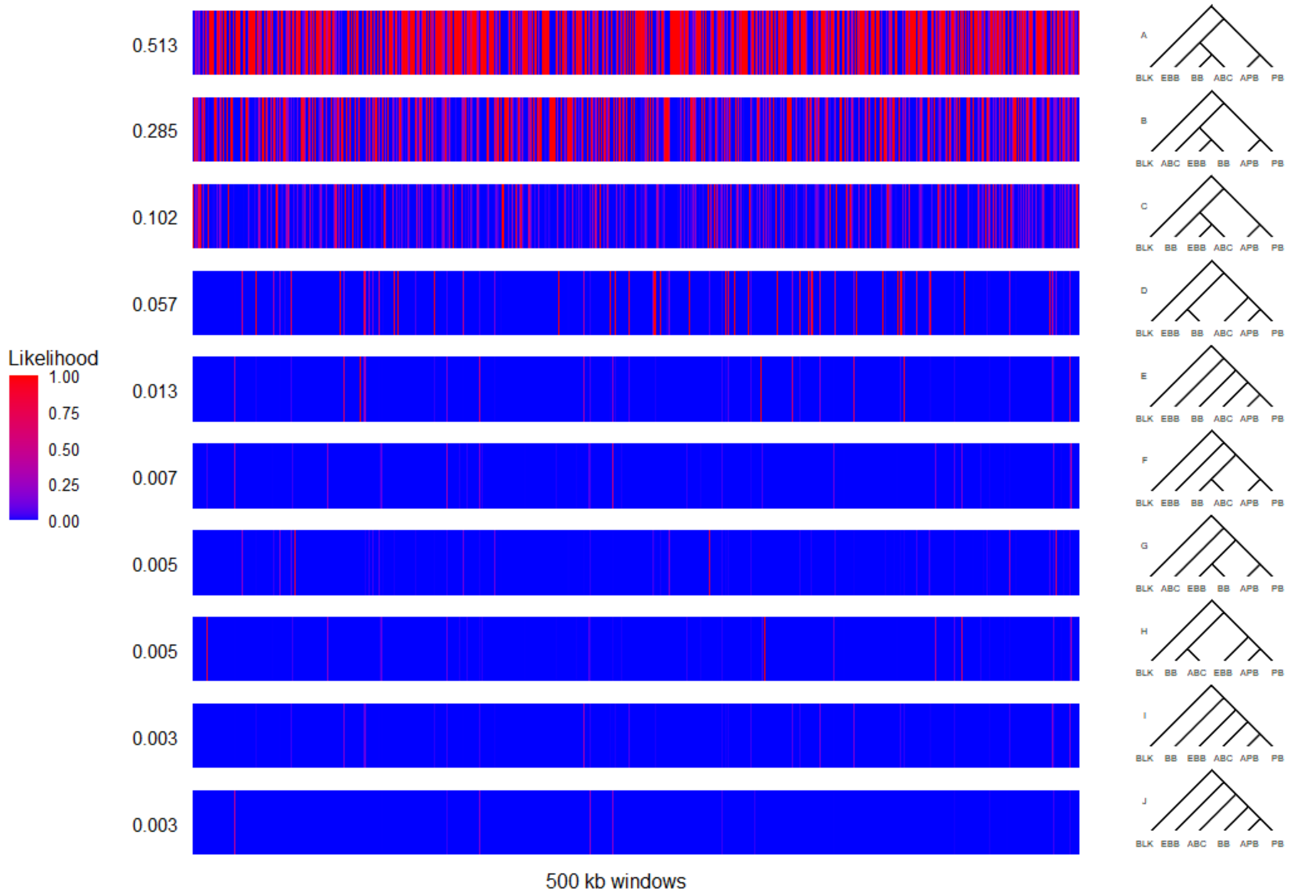


**Fig. S21.** The likelihood of the top ten best fitting trees among the 105 different trees according to admixturegraph analysis using $f_2$-statistics within 500 kb windows. The reported numerical value is the average over all the windows.

A.

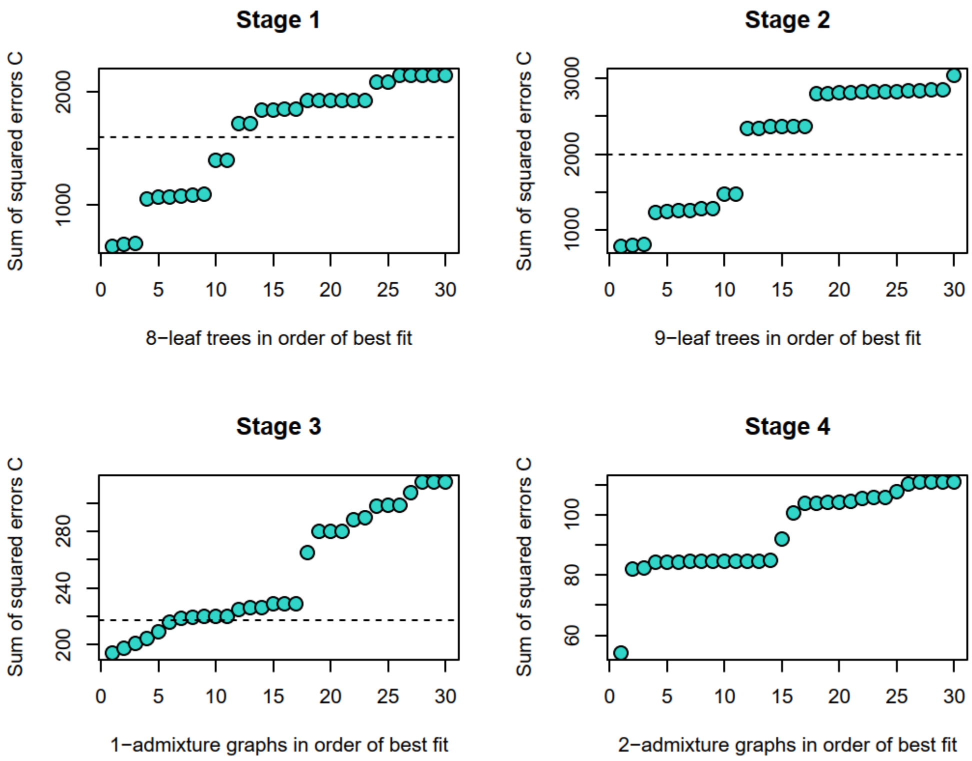
Stage 1
Sum of squared errors C
8−leaf trees in order of best fit
Stage 2
Sum of squared errors C
9−leaf trees in order of best fit
Stage 3
Sum of squared errors C
1−admixture graphs in order of best fit
Stage 4
Sum of squared errors C
2−admixture graphs in order of best fit

B.

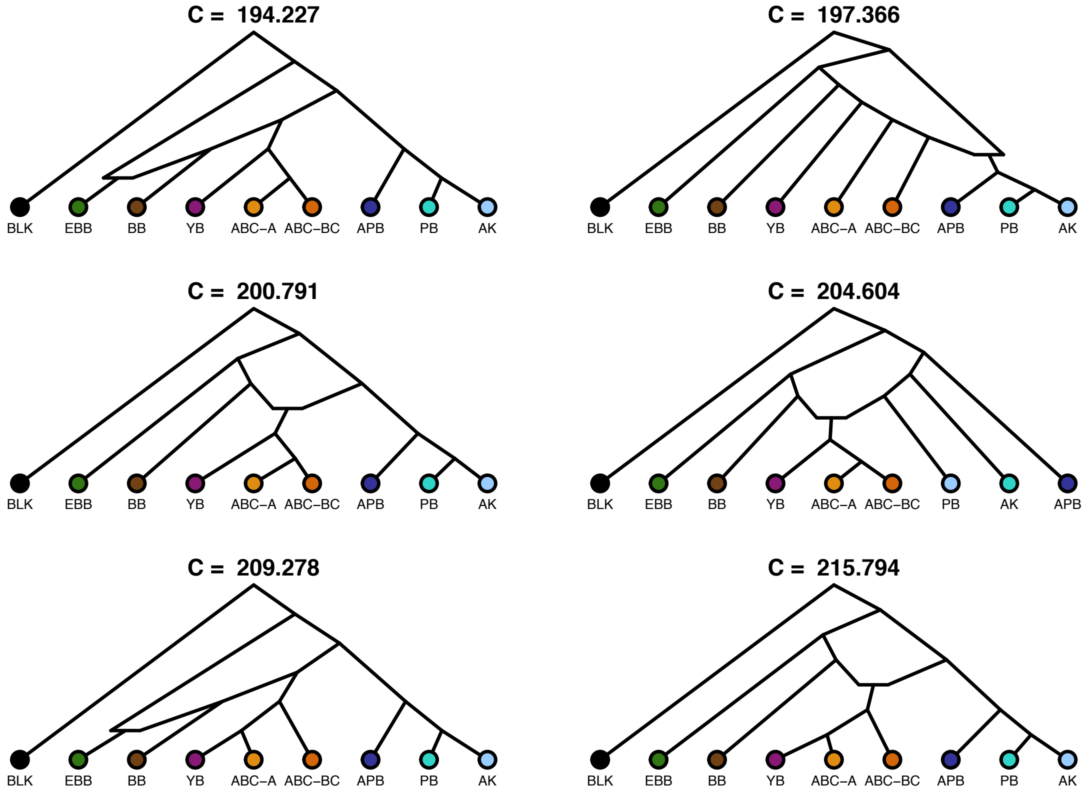
C = 194.227
BLK EBB BB YB ABC−A ABC−BC APB PB AK
C = 197.366
BLK EBB BB YB ABC−A ABC−BC APB PB AK
C = 200.791
BLK EBB BB YB ABC−A ABC−BC APB PB AK
C = 204.604
BLK EBB BB YB ABC−A ABC−BC PB AK APB
C = 209.278
BLK EBB BB YB ABC−A ABC−BC APB PB AK
C = 215.794
BLK EBB BB YB ABC−A ABC−BC APB PB AK

**C.**

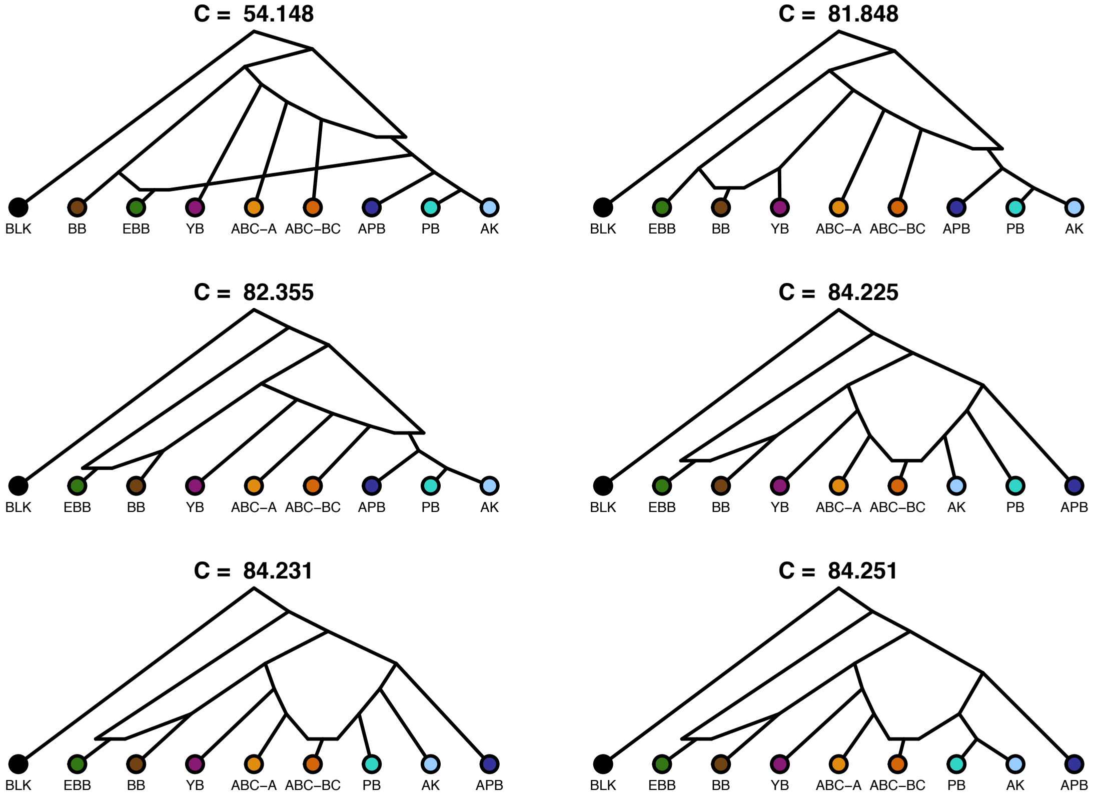


**Fig. S22.** The admixturegraph analysis using $f_2$-statistics. **A.** Stages 1–4. Note that graphs where BLK appeared as an admixed population were removed after Stage 4, the quality of fit among those was 57.092 at best. **B.** The best fitting graphs after Stage 3. **C.** The best fitting graphs after Stage 4.

**A.**

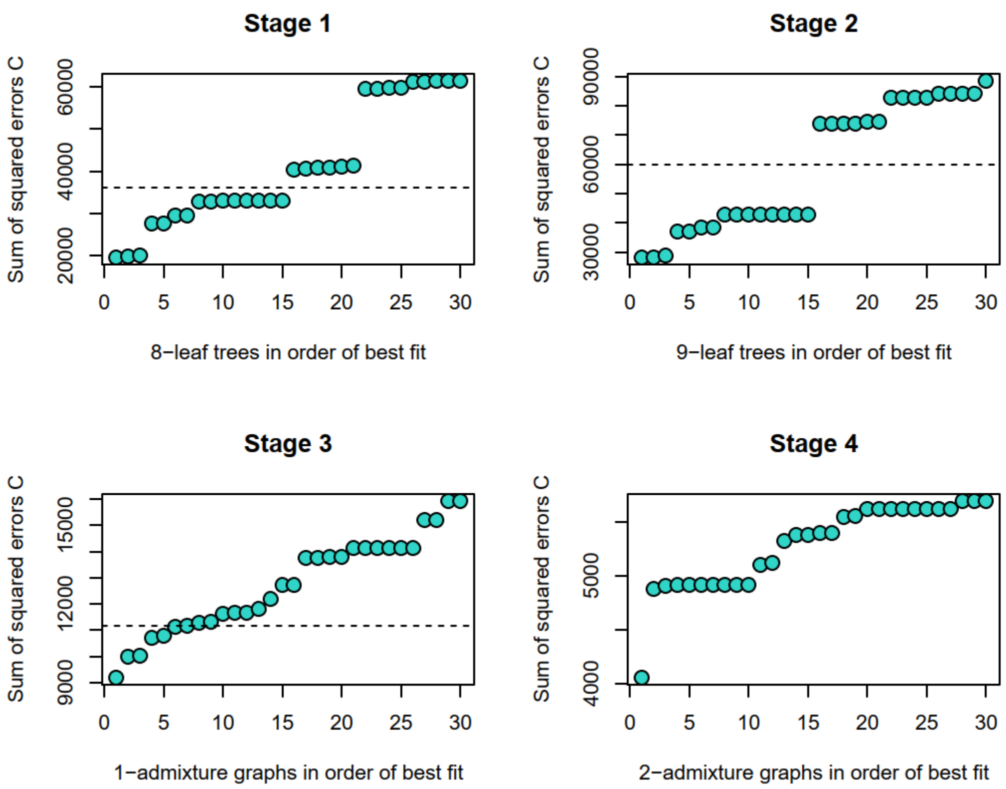
Stage 1
Sum of squared errors C
20000
40000
60000
0 5 10 15 20 25 30
8−leaf trees in order of best fit
Stage 2
Sum of squared errors C
30000
60000
90000
0 5 10 15 20 25 30
9−leaf trees in order of best fit
Stage 3
Sum of squared errors C
9000
12000
15000
0 5 10 15 20 25 30
1−admixture graphs in order of best fit
Stage 4
Sum of squared errors C
4000
5000
0 5 10 15 20 25 30
2−admixture graphs in order of best fit

**B.**

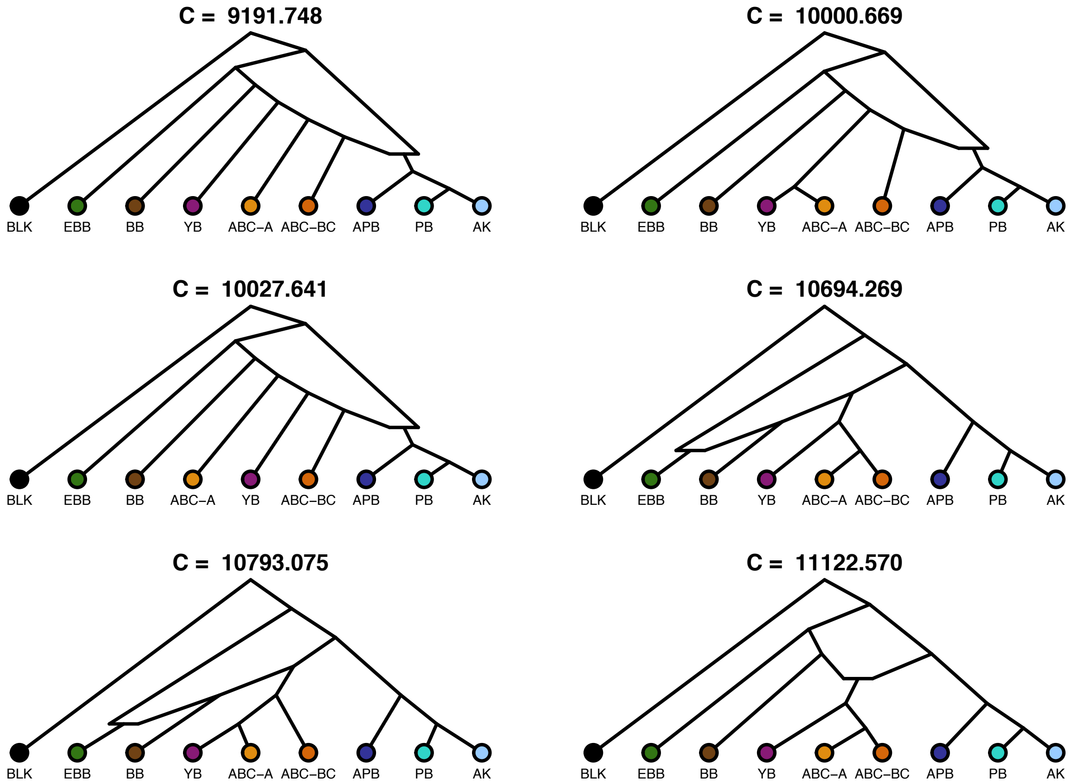
C = 9191.748
BLK EBB BB YB ABC−A ABC−BC APB PB AK
C = 10000.669
BLK EBB BB YB ABC−A ABC−BC APB PB AK
C = 10027.641
BLK EBB BB ABC−A YB ABC−BC APB PB AK
C = 10694.269
BLK EBB BB YB ABC−A ABC−BC APB PB AK
C = 10793.075
BLK EBB BB YB ABC−A ABC−BC APB PB AK
C = 11122.570
BLK EBB BB YB ABC−A ABC−BC APB PB AK

C.

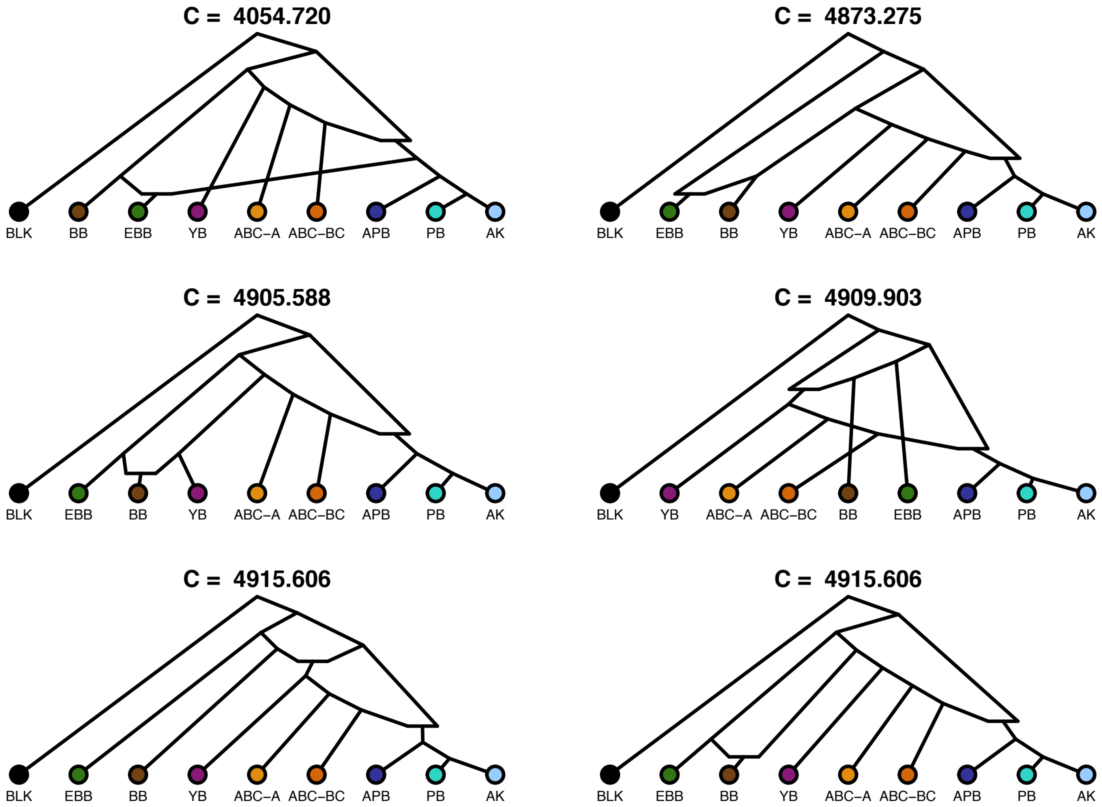


**Fig. S23.** The admixturegraph analysis using $f_4$-statistics. **A.** Stages 1–4. Note that graphs where BLK appeared as an admixed population were removed after Stage 4, the quality of fit among those was 3584.543 at best. **B.** The best fitting graphs after Stage 3. **C.** The best fitting graphs after Stage 4.

**A.**

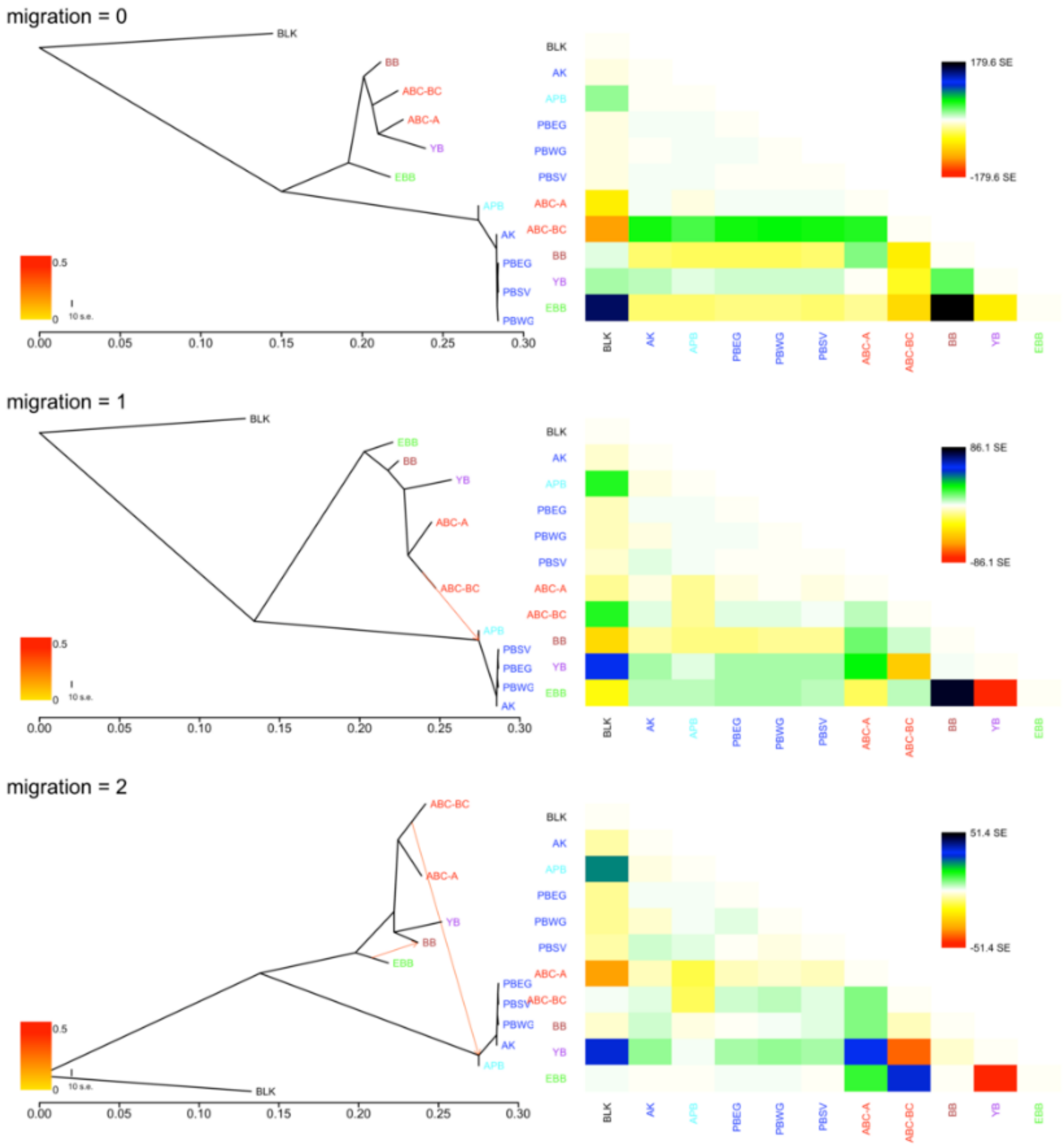

migration = 0
BLK
BB
ABC-BC
ABC-A
YB
EBB
APB
AK
PBEG
PBSV
PBWG
0.5
0
10 s.e.
0.00
0.05
0.10
0.15
0.20
0.25
0.30
179.6 SE
-179.6 SE
migration = 1
86.1 SE
-86.1 SE
migration = 2
51.4 SE
-51.4 SE

migration = 3

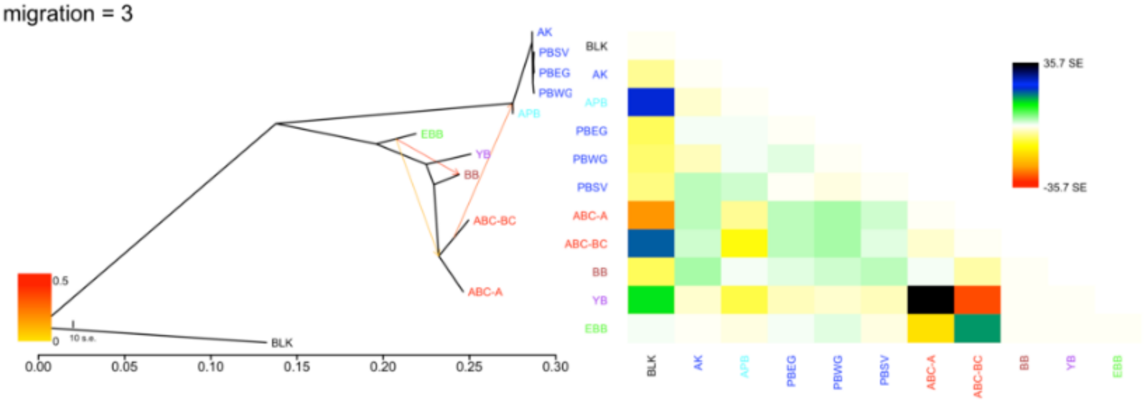


migration = 4

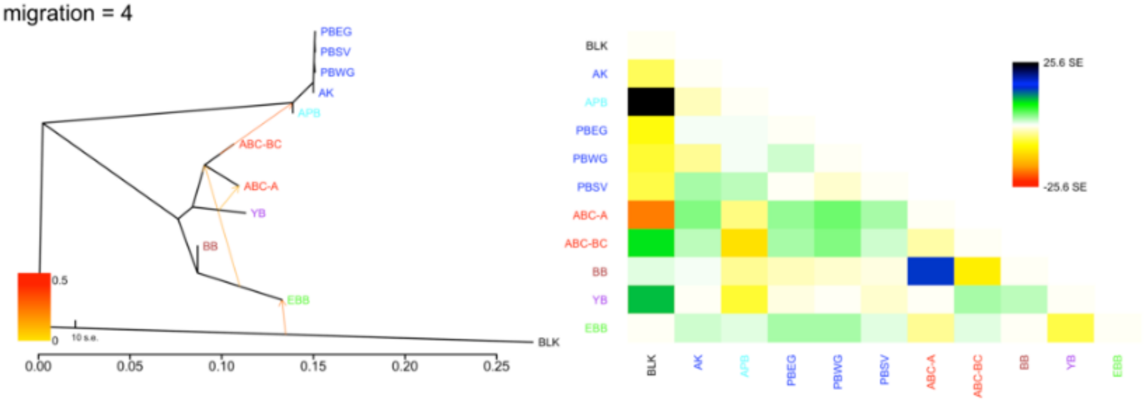


migration = 5

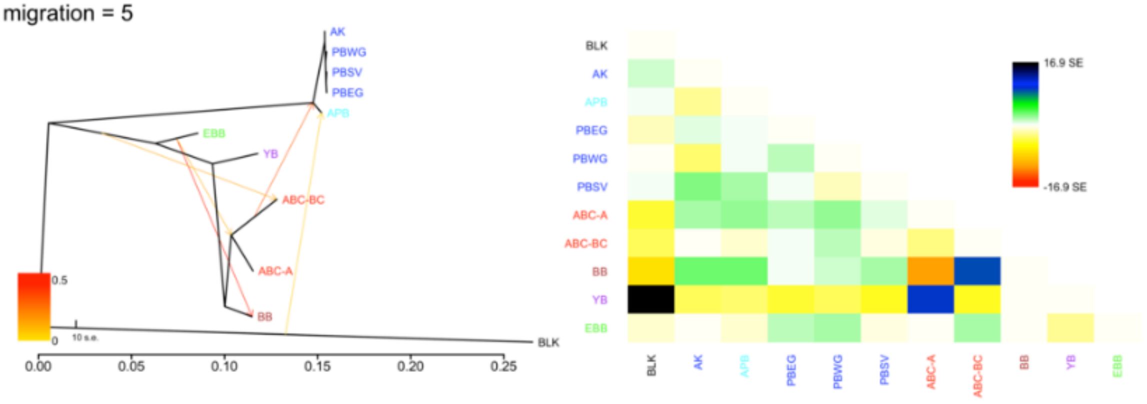

**B.**

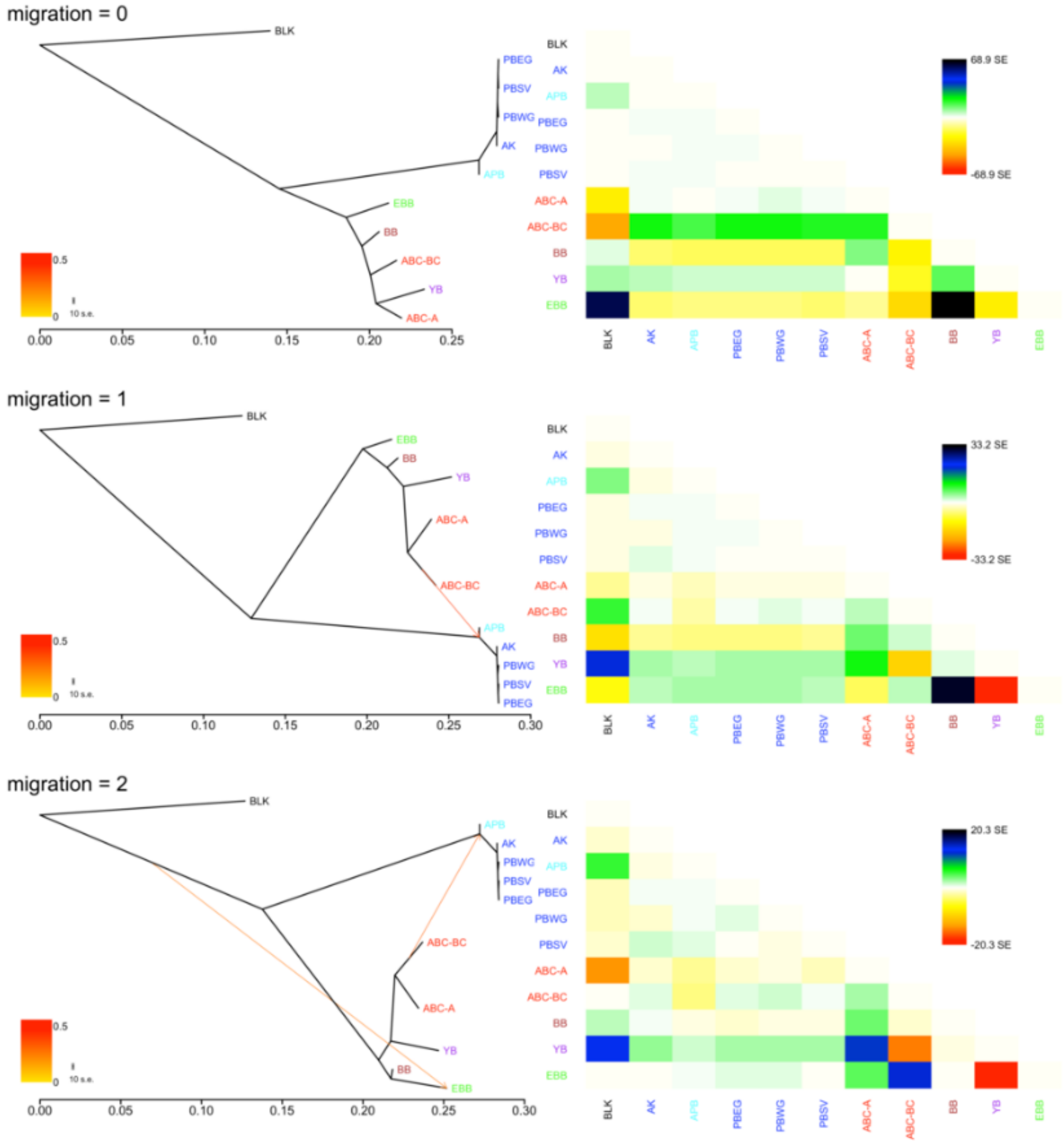

migration = 0
migration = 1
migration = 2
BLK
AK
APB
PBEG
PBWG
PBSV
ABC-A
ABC-BC
BB
YB
EBB
68.9 SE
-68.9 SE
33.2 SE
-33.2 SE
20.3 SE
-20.3 SE
0.5
0
10 s.e.
0.00
0.05
0.10
0.15
0.20
0.25
0.30

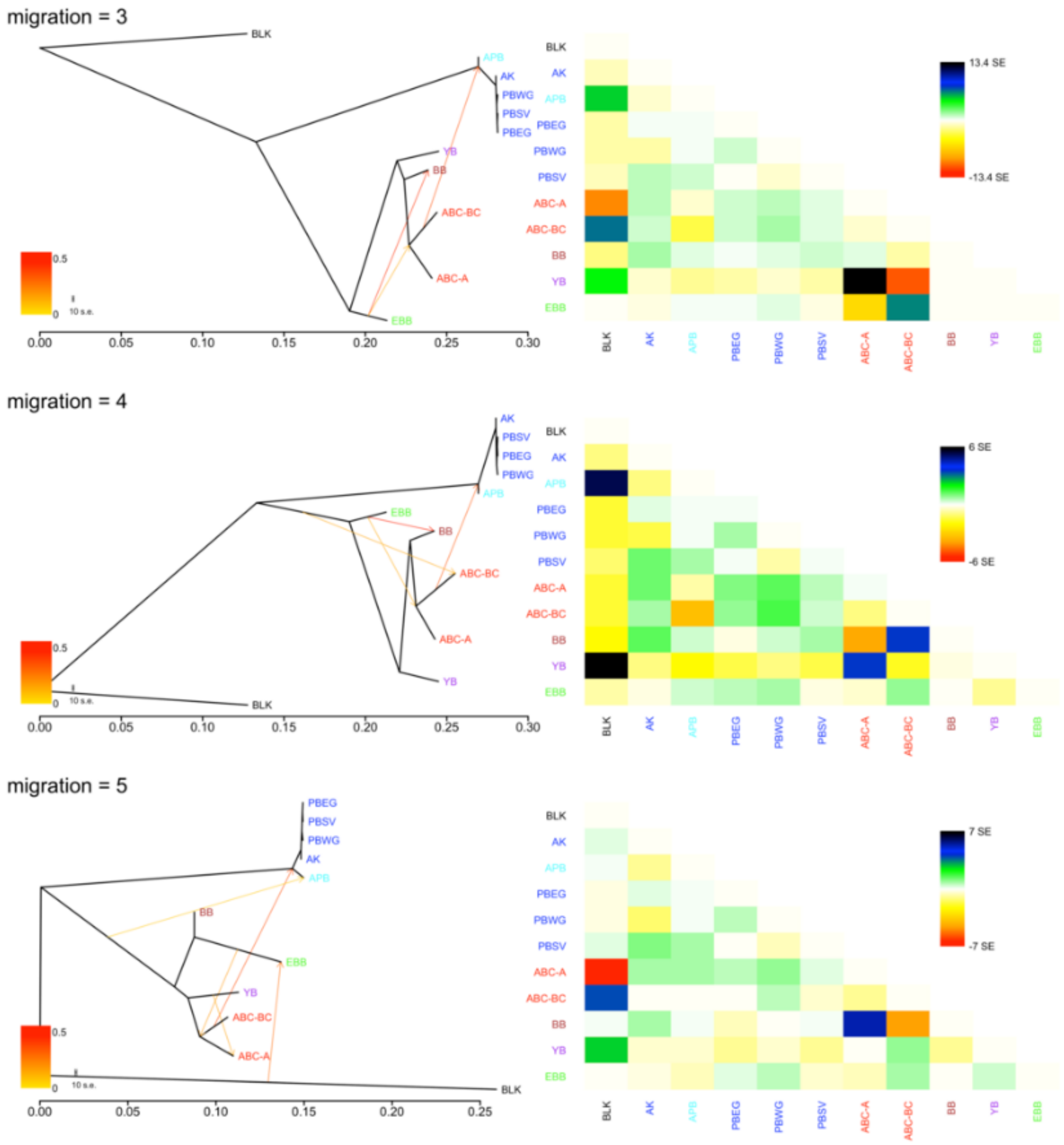


**Fig. S24.** TreeMix showing 0–5 migration events for populations of brown and polar bears based on: **A.** DS3 (all SNPs from genomes > 8X coverage, excluding private alleles). **B.** DS8 (DS3 but transversions only).

**A.**

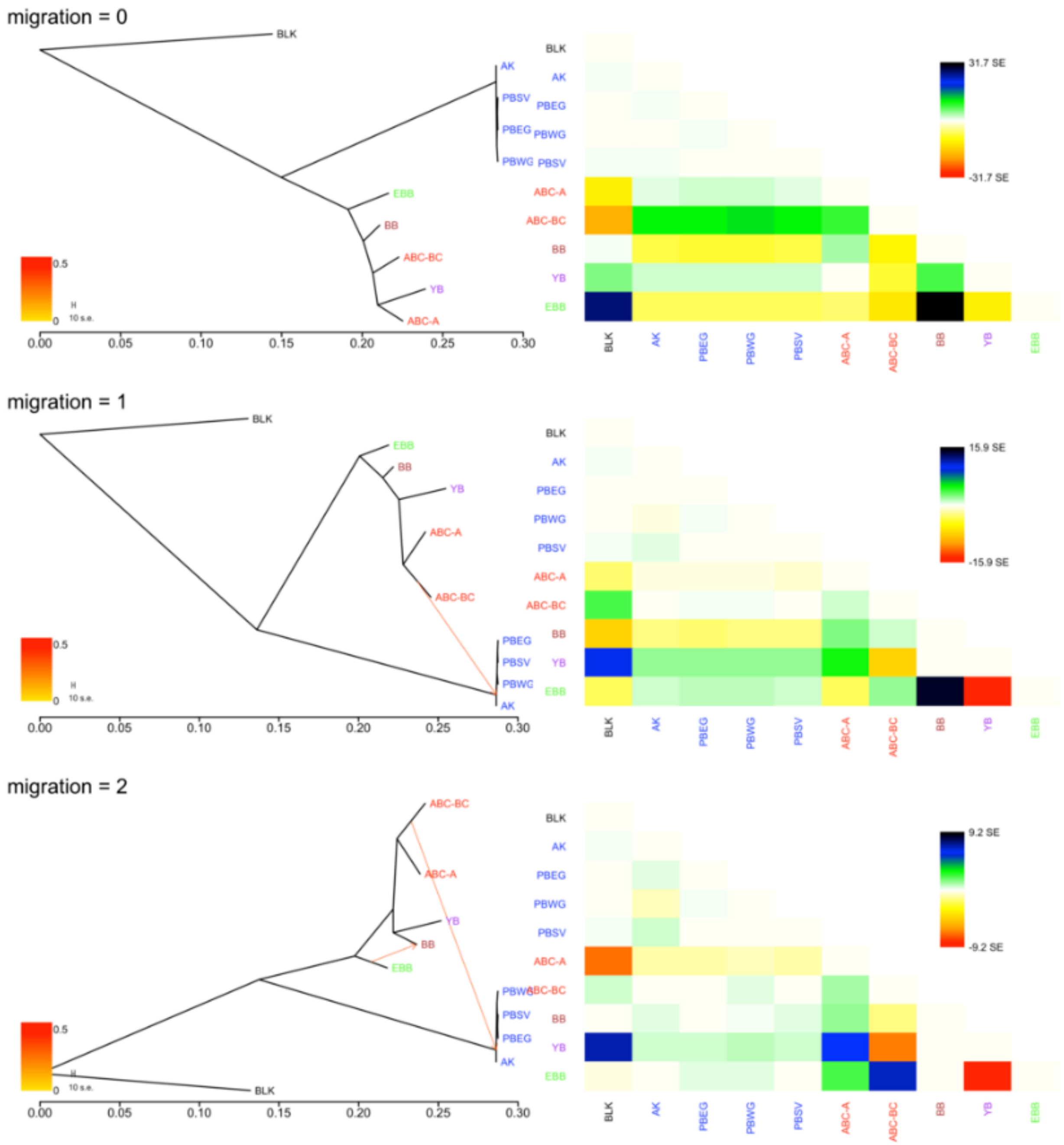
migration = 0
BLK
AK
PBSV
PBEG
PBWG
EBB
BB
ABC-BC
YB
ABC-A
0.5
0
10 s.e.
0.00
0.05
0.10
0.15
0.20
0.25
0.30
31.7 SE
-31.7 SE
migration = 1
15.9 SE
-15.9 SE
migration = 2
9.2 SE
-9.2 SE

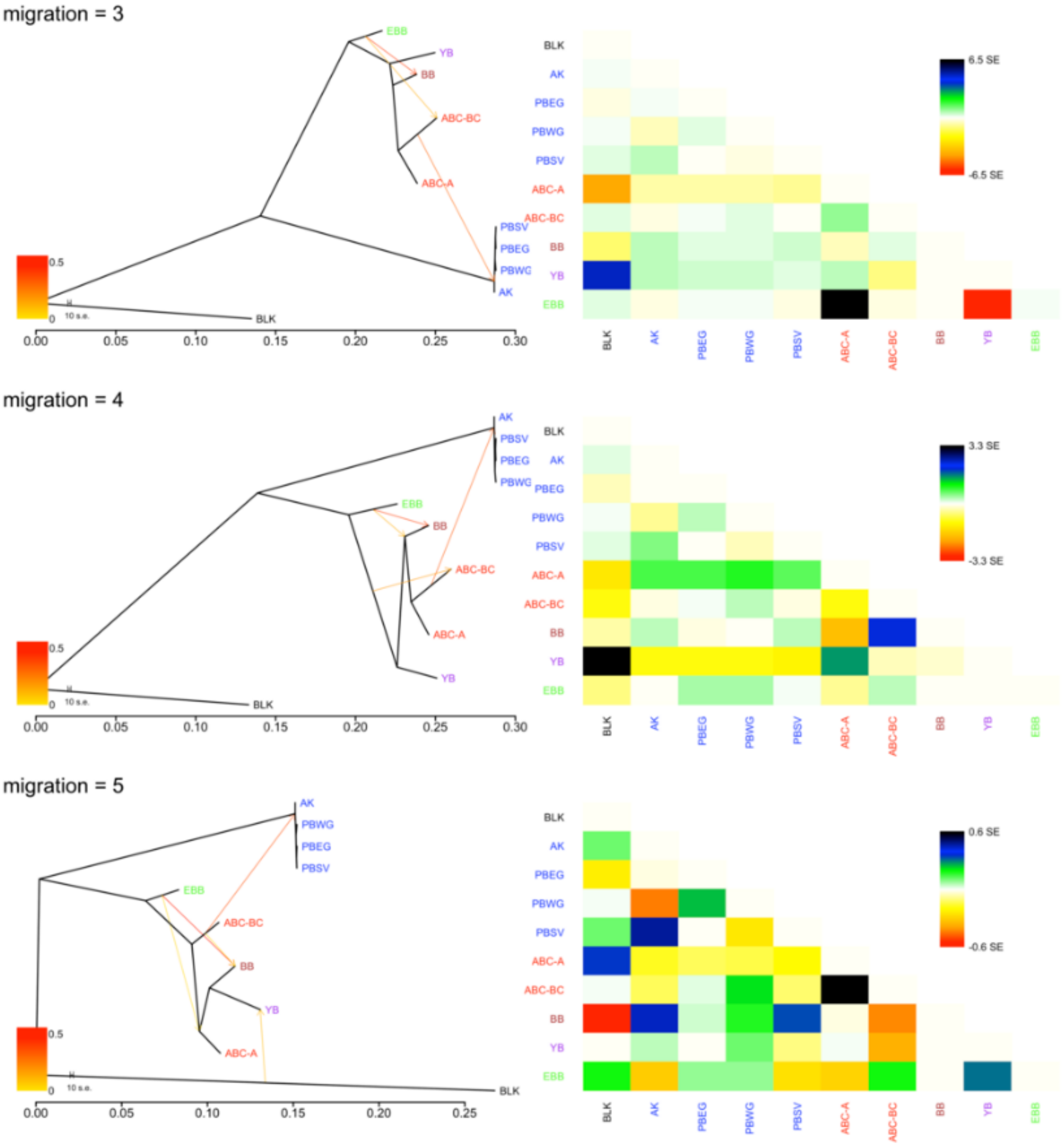
migration = 3
EBB
YB
BB
ABC-BC
ABC-A
PBSV
PBEG
PBWG
AK
BLK
0.5
0
10 s.e.
0.00
0.05
0.10
0.15
0.20
0.25
0.30
6.5 SE
-6.5 SE
migration = 4
3.3 SE
-3.3 SE
migration = 5
0.6 SE
-0.6 SE

**B.**

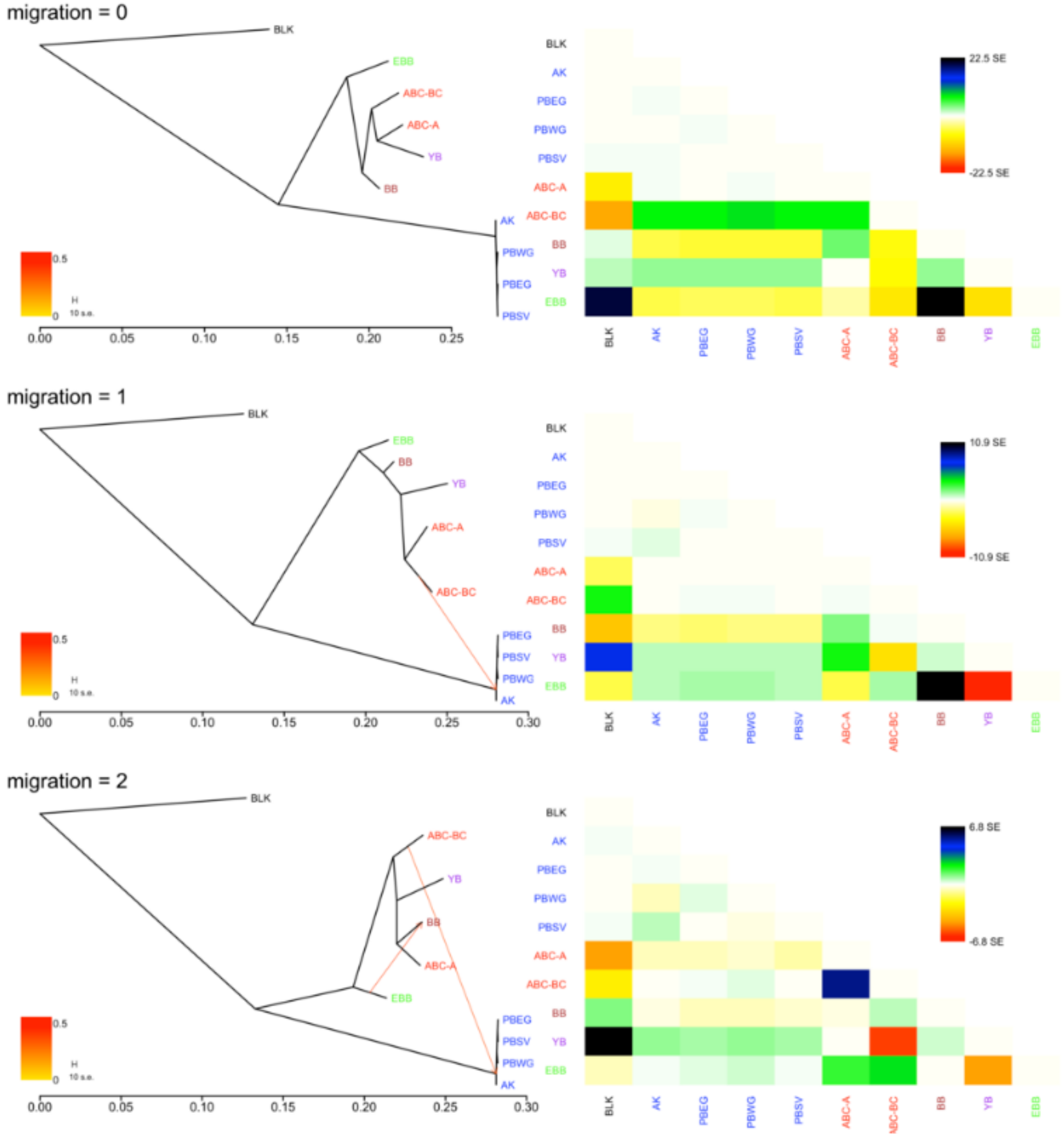
migration = 0
BLK
EBB
ABC-BC
ABC-A
YB
BB
AK
PBWG
PBEG
PBSV
0.5
0
H
10 s.e.
0.00
0.05
0.10
0.15
0.20
0.25
22.5 SE
-22.5 SE
migration = 1
10.9 SE
-10.9 SE
0.30
migration = 2
6.8 SE
-6.8 SE

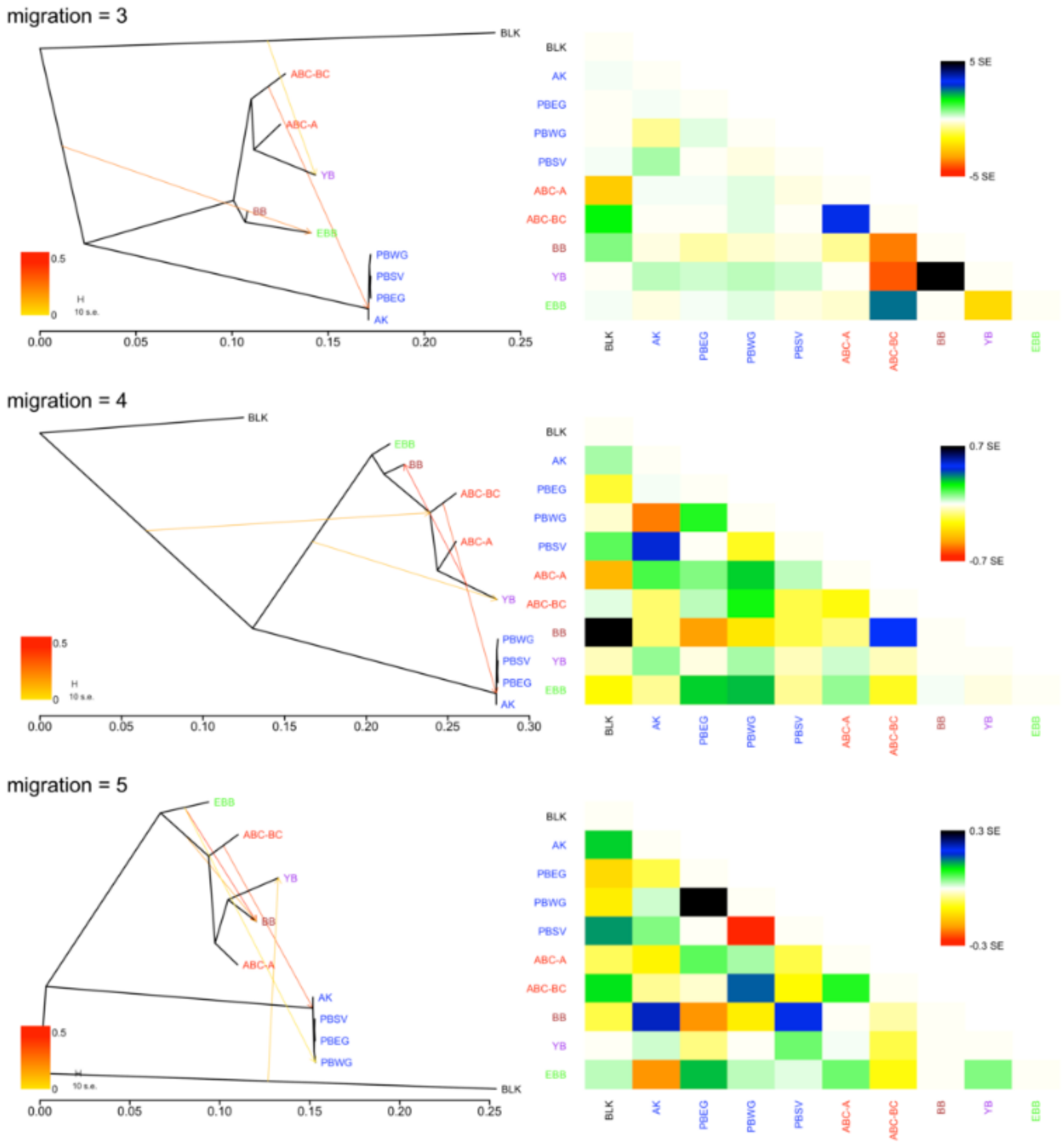


**Fig. S25.** TreeMix showing 0–5 migration events for populations of brown and polar bears, but excluding the ancient polar bear, based on: **A.** DS3 (all SNPs from genomes > 8X coverage, excluding private alleles). **B.** DS8 (DS3 but transversions only).
